# Evaluation of K-Ras4B dimer interfaces and the role of Raf effectors

**DOI:** 10.1101/2022.10.04.510804

**Authors:** Alexios Chatzigoulas, Ioannis Andreadelis, Stefan Doerr, Christos Lamprakis, Anastasia Theodoropoulou, John Manchester, Camilo Velez-Vega, Jose Duca, Zoe Cournia

## Abstract

K-Ras4B is one the most frequently mutated proteins in cancer, yet mechanistic details of its activation such as its homodimerization on the membrane remain elusive. The structural determinants of K-Ras4B homodimerization have been debated with different conformations being proposed in the literature. Here, we perform microsecond all-atom Molecular Dynamics (MD) simulations on the K-Ras4B monomer in solution, the K-Ras4B monomer on the membrane, and two experimentally-based K-Ras4B dimer models of the α4-α5 interface to investigate the stability of these structures bound to GTP on a model cell membrane. We then evaluate the complexes for their propensity to form stable dimers on the plasma membrane in the presence and absence of Raf[RBD–CRD] effectors. We find that Raf[RBD-CRD] effectors enhance dimer stability, suggesting that the presence of effectors is necessary for K-Ras4B dimers stabilization on the cell membrane. Moreover, we observe, for the first time, a dynamic water channel at the K-Ras4B dimer interface, and identify putative allosteric connections in the K-Ras4B dimer interface. To discover novel K-Ras4B interfaces, we perform coarse-grained MD simulations in two dissociated K-Ras4B monomers on the membrane, which reveal that the dominant dimer interface is the α4-α5 interface. Finally, a druggability analysis is performed in the different K-Ras4B structures in the monomeric states. Strikingly, all known binding pockets of K-Ras4B are identified only in the structure that is membrane-bound, but not in the solution structure. Based on these results, we propose that modulating the protein-membrane interactions can be an alternative strategy for inhibiting K-Ras4B signaling.

## INTRODUCTION

The Ras family is comprised of H-Ras, N-Ras, and the two variants K-Ras4A and K-Ras4B proteins, which are small GTPases mediating signaling pathways essential for cell proliferation and connecting various transmembrane receptors to activating transcription factors in the nucleus.^1, 2^ Approximately one-third of all human cancers harbor an oncogenic mutation in at least one *RAS* gene, rendering Ras proteins the most commonly mutated oncoproteins.^3^ Among them, *KRAS* is the most frequently mutated gene in human cancers (83% of all RAS mutations), while in some pancreatic adenocarcinomas it is mutated in nearly 90% of tumors. Even though Ras proteins are a highly conserved family, K-Ras is characterized by unique stem-like properties,^4^ higher gene expression,^5^ and poor rate of DNA repair^6^ compared to H-Ras and N-Ras.

Ras proteins are comprised of a catalytic domain (residues 1 to 166) and a hypervariable C-terminal region (HVR) (residues 167 to 188/189). The catalytic domain, also called G-domain, consists of two structurally distinct regions: the effector lobe (residues 1 to 86), which recruits downstream signaling effectors, and the allosteric lobe (residues 87-167), which is proposed to be implicated in membrane interactions.^7^ The catalytic domain includes Switch I and Switch II regions, which are connected through the GTP γ-phosphate and undergo significant conformational changes upon hydrolysis of GTP to GDP in the protein.^8^ The hypervariable C-terminal region is highly flexible and divergent amongst Ras family members. In the full-length GDP-bound wild type (WT) K-Ras4B, HVR interacts tightly with the catalytic domain, constraining K-Ras4B in an intrinsically autoinhibited state.^8^ On the contrary, looser HVR-catalytic domain interactions occur in full-length GTP-bound WT K-Ras4B, leading to the release of HVR and its subsequent association with the cell membrane.^9^

Ras proteins function as binary switches cycling between active (GTP-bound) and inactive (GDP-bound) conformational states. The γ-phosphate of GTP keeps Switch I and Switch II bound in a stable conformation via the main chain NH groups of the invariant Thr and Gly amino acids, while the release of the γ-phosphate after GTP hydrolysis allows the switch regions to relax into a different conformation.^10^ Although the intrinsic GTPase activity of K-Ras4B is low, it can be accelerated by orders of magnitude by GTPase activating/accelerating proteins (GAPs), leading to the inactive GDP-bound K-Ras4B state after hydrolysis.^11^ The hydrolysis cycle is also regulated by Guanine Nucleotide Exchange Factors (GEFs), which are proteins recruited as a response to several growth receptors that promote the release of GDP in favor of the 10-fold higher cellular concentrated GTP.^12^ Oncogenic mutations in all Ras proteins impair GAP-mediated hydrolysis of GTP, resulting in the stabilization of the active GTP-bound Ras state, hindering hydrolysis, and consequently in the hyper-activation of upstream signaling. Dominant K-Ras4B mutations cause tumorigenesis to occur at codons 12, 13, and 61,^3^ with each of them prone to engaging different downstream effectors.^13^

Active GTP-bound K-Ras4B can be anchored on the plasma membrane after its post-translational farnesylation at the C-terminal cysteine of the HVR, and subsequently engage a plethora of effectors at its effector lobe such as Raf kinase^14, 15^ and Ral guanine nucleotide dissociation stimulator (RalGDS).^16^ The interaction of K-Ras4B with these effectors activates signaling pathways related to cell growth and proliferation, including the Ras-Raf-MEK-ERK and Ras-PI3K-PDK1-AKT pathways.^17^ In the MEK-ERK signaling pathway, which is responsible for cell proliferation, K-Ras4B recruits Raf-1 kinase (hereafter mentioned as “Raf”) through the Raf Ras binding domain (RBD) and Raf cysteine-rich domain (CRD) to the plasma membrane (Figure S1).^18^ Cytosolic Raf exists in an inactive, auto-inhibited conformational state; however, it is activated upon recruitment on the plasma membrane by K-Ras4B.^19^ Ras membrane dimerization plays an important role in the Ras-Raf association and subsequent regulation of the MEK-ERK pathway. It has been proposed that monomeric Ras can bind Raf, but activation of Raf requires Raf dimerization, which can be achieved using Ras dimers as a scaffold on the membrane (Figure S1).^20, 21^ The increased local concentration of K-Ras4B on the membrane-mediated by anionic lipid binding specificity favors the formation of K-Ras4B dimers or higher-order structures,^22-24^ and consequently promotes K-Ras4B-Raf association and the homodimerization and transactivation of the Raf kinase domain.^25^ Activation of Raf through its dimerization leads to the phosphorylation and activation of MEK, transmitting the signal down to ERK.^26^

The debate whether K-Ras4B dimerizes on the cell membrane is still ongoing as another study could not observe K-Ras4B in a dimeric state.^27^ In this study, biochemical evaluation of K-Ras4B in a broad range of lipid bilayers indicated that WT K-Ras4B exists in a monomeric state that lacks intrinsic dimerization ability.^27^ However, it should be noted that these studies have been performed in cell-free environments lacking effectors such as B-Raf.^27^ This points to the question of whether K-Ras4B has an intrinsic dimerization capability or it needs effectors to dimerize. Recently, it was experimentally shown that the RBD domain induces robust Ras dimerization.^28^ It is well-established that the Raf RBD domain binds to the effector lobe of active membrane-bound Ras with nanomolar affinity,^29, 30^ while the CRD domain of Raf, which is connected to the RBD domain of Raf through a disordered linker, anchors to the membrane and interacts with phospholipids, specifically to negatively charged phospholipids such as phosphatidic acid (PA) and phosphatidyl serine (PS),^31^ although recently a different Ras/Raf dimer assembly was proposed with CRD distal from the membrane.^32^ The RBD domain of Raf is strongly attached to the K-Ras4B effector binding region, limiting the accessibility of this K-Ras4B domain to the membrane, while the CRD domain possesses a low affinity for K-Ras4B and is strongly tethered to the membrane through electrostatic interactions.^33^ Thus, the competition for protein-membrane contacts between the CRD domain of Raf and K-Ras4B is a key factor leading to limiting the K-Ras4B configurational ensemble on the membrane and could enhance dimer stability.^34-37^

The continuous quest for K-Ras4B targeting therapeutic strategies has insofar mainly concentrated on the inhibition of its binding to the various upstream and downstream effectors^38-41^ as well as indirect inhibition strategies of the K-Ras4B partners, e.g. inhibition of the PDEδ protein that transports farnesylated K-Ras in the cytosol leads to disruption of the PDEδ–K-Ras interaction, thereby hindering HVR from anchoring to the plasma membrane.^42^ The direct inhibition of K-Ras4B is strenuous due to the picomolar affinity of KRAS for its substrates, and the lack of deep hydrophobic pockets.^43, 44^ One such success story was the design of G12C GDP-bound KRAS selective inhibitors with promising cellular activity.^45^ These inhibitors suppress G12C KRAS mutant signaling and tumor cell growth,^46^ by inhibiting downstream signaling. Recently, a potent G12C covalent allosteric inhibitor named sotorasib (AMG510) obtained FDA approval and became the first therapy to directly target the KRAS oncoprotein.^47^ However, the aforementioned inhibition is mutant-specific and targets only the G12C KRAS, a mutation not frequently observed in the clinic, as it is present in the lung but not in pancreatic and colorectal cancers.^3^ Recently, another approach for targeting K-Ras4B arose from cumulative evidence that membrane-bound K-Ras proteins, including K-Ras4B, form dimers or even nanoclusters in order to activate signaling pathways, with the dimer proposed to be the basic clustering unit.^20, 24, 48-53^ K-Ras dimerization aids the regulation of signaling pathways by providing signaling specificity, allosteric effects, and effector activation and inhibition. Indeed, it has been shown that WT K-Ras acts as a tumor suppressor in *KRAS* mutant cancer cells^54^ by antagonizing the function of oncogenic K-Ras when forming WT-mutant dimers, which however no longer occurs when the WT *KRAS* allele is lost during tumor progression.^55, 56^

Several studies suggest the existence of K-Ras dimer,^20, 57-61^ trimer,^62^ and also pentamer conformations.^63^ Dimerization is critical for the activity of WT K-Ras as well as for the oncogenic ability of mutant G12D K-Ras, according to a study that investigated the K-Ras D154Q mutant does not have the ability to dimerize.^60^ This study showed that K-Ras dimerization is essential for activation of downstream signaling and cell growth *in vitro* and *in vivo*, and suggested a role for disruption of dimerization as a therapeutic strategy for *KRAS* mutant cancers. At the same time, Ras dimers have been observed in the biological assembly of an X-ray crystal structure^64^ and NMR-inspired structures.^61^ Different dimer interfaces have been suggested using both computational and experimental methods for members of the Ras family,^52, 53^ proposing several interfaces for membrane-bound Ras dimers such as the region between the α4-α5 helices and the loop between β2–β3 sheets for N-Ras,^48, 65^ the α4–β6–α5 helical interface for H-Ras,^51^ the α4–α5 interface for K-Ras,^60, 61, 64^ the β2 strands or the α3–α4 helical interface for K-Ras,^20, 66-68^ and dimer assemblies with GTP in the interface for K-Ras.^69, 70^ For the K-Ras4B G12D mutant similar interfaces have also been observed; the β2 interface in crystal structures of the K-Ras4B G12D mutant (PDB ID: 7ACF^71^, 6GJ7^72^) as well as the α4–α5 interface (PDB ID 5US4^73^, 5XCO^74^).

The critical importance of the α4-α5 interface in WT Ras dimerization and function was demonstrated in Ref.^60^ which studied the D154Q mutant shown to abolish both WT and mutant K-Ras homo-dimerization, without influencing its intrinsic GTPase activity, GEF or GAP sensitivity and C-Raf binding. Specifically, at the α4-α5 dimer interface, which is also present in the 5VQ2 PDB crystal structure,^64^ residues D154 and R161 of the two different K-Ras4B monomers form two intermolecular salt bridges; abolishing this interaction interrupts K-Ras4B dimerization, while at the same time its other biochemical properties are not affected.^60^ Furthermore, a study by Spencer-Smith et al. demonstrated that a designed monobody, namely NS1, interacts with the catalytic domain of both GTP- and GDP-bound K-Ras and disrupts K-Ras dimerization through the α4-α5 interface,^51^ also inhibiting oncogenic K-Ras in vivo.^75^ Another recent study also reported the α4-α5 region as the likely dimerization interface but proposed a different orientation of K-Ras4B membrane-associated homodimers bound to GTP and GDP.^61^ Therein, the authors studied K-Ras4B dimerization on nanodiscs via paramagnetic relaxation enhancement (PRE) NMR, and the structures of membrane-anchored KRAS dimers in the active GTP- and inactive GDP-loaded states were modeled using the 4DSO crystal structure as reference^39^ and conforming with PRE-derived distances. For the active GTP state model, K-Ras4B was found to dimerize through an α4-α5 dimer interface, albeit distinct from the one proposed in Ref.^60^ In Ref.^61^ helices α4 and α5 of opposing monomers form a 20° angle with each other, while in Ref.^60^ α5 is rotated 90° with respect to α4 and α5 helix of the opposite monomer, forming an α4-α5 interface, which includes α5-α5 and α4-α4 interfaces (Figure S2). Moreover, in Ref.^61^ the dimer interface is stabilized by hydrogen bonds from Q131 to D154 and R161 and a salt bridge between R135 of one monomer and E168 of the second monomer, while in Ref.^60^ an intermolecular salt bridge between R161 and D154 is stabilizing the interface (Figure S3). These observations suggest that the α4-α5 interface may be the one of the most viable for K-Ras dimerization; however, it remains unclear, which conformation is the biologically-relevant interface of the K-Ras4B dimer.

Molecular dynamics (MD) simulations of the K-Ras4B monomer in the presence and absence of model membranes have been studied in the literature, and distinct conformations with varying orientations of the protein on the membrane have been described.^34-36, 76-83^ The formation and stability of the K-Ras4B dimer have been also described in Refs ^20, 28, 50, 58, 59, 63, 68, 69^. To create different potential dimer interfaces, computational methods such as PRISM^84, 85^ and RosettaDock^86^ have been also employed. In a study evaluating K-Ras4B-GTP dimer interfaces, four different interfaces were generated with the PRISM^84, 85^ protein-protein complex structure prediction algorithm.^58^ These models were simulated with MD in solution and in DOPC:DOPS (4:1) membrane systems (200 ns each). In these simulations, the α3–α4 and β2 interfaces were the most stable.^58^ Based on these results, three different orientational states of the α3-α4 interface were further evaluated with MD simulations along with a symmetric α4–α5 interface used as a control system in DOPC:DOPS (4:1) membrane systems for 1 μs each, this time for the K-Ras4B-Raf[RBD-CRD] complex, resulting in stable α3–α4 interfaces throughout the simulations and separation of the α4–α5 interface.^68^ In another study aiming to characterize the K-Ras4B dimer interface, six different dimer interfaces were modeled with the RosettaDock^86^ protein-protein docking tool and simulated in DOPC:DOPS (∼3:1) membrane systems (∼200 ns each).^59^ The two most stable dimer models, α3–α4 and α4–α5 interfaces, were further simulated in solution for 400 ns each, being stable without dimer dissociation. These dimer models were then used for the creation of trimer, tetramer, and pentamer models which were simulated in a DOPC:DOPS (5:1) membrane for 1 μs with stable PPIs.^63^ Recently, the K-Ras4B-Raf[RBD] α4–α5 interface dimer was modeled by applying twofold crystallographic symmetry to the asymmetric unit of the H-Ras-Raf[RBD] complex (PDB ID 4G0N)^87^ and two independent 1 μs MD simulations were performed in a DOPC:DOPS (4:1) membrane.^28^ The dimer interface was stable in the simulations, and allosteric network analysis indicated that dimerization increases allosteric connections at opposite ends of the dimer formed by the α4−α5 interface. In a large-scale study, two K-Ras4B-GTP monomers without effectors were placed in arbitrary positions and without being in contact, and 20 MD simulations were performed in solution (total time: 680 μs) and another 23 MD simulations on the membrane in a POPC:POPS (70:30) membrane (total time: 363 μs).^69^ The spontaneous dimerization of the α4–α5 dimer interface was observed in a simulation, but in other simulations an interface involving GTP on the dimer interface emerged, that has not been previously reported. The GTP-binding region was also found at the K-Ras4B dimer interface of another millisecond MD simulations study, although with a different dimer configuration.^70^ In this study, 50 K-Ras4B structures were extracted from previous monomer K-Ras4B simulations,^83^ and then were placed distant from each other in different orientations to generate 144 different initial configurations. Each configuration was simulated for 7 μs without effectors and on a POPC:POPS (71:29) membrane, discovering 25 stable K-Ras4B dimer interfaces. A stable α4-α5 dimer interface was also found in the simulations, with the α4 and α5 helices being parallel to the membrane plane and the α4 helices being in contact with the membrane.

In the present study, we use atomistic MD simulations to investigate the stability of two experimentally-based K-Ras4B dimer structures forming different α4–α5 dimerization interfaces^60, 61^ bound to GTP on a model cell membrane with and without the presence of Raf[RBD-CRD] effectors. We find that Raf[RBD-CRD] effectors enhance dimer stability and the α4–α5 interface, suggesting that the presence of effectors is necessary for K-Ras4B dimers to stabilize on the cell membrane. Comparison of the two different dimerization interfaces based on structural and interaction energy analysis results in different strengths and weaknesses of the two α4–α5 interfaces. The α4-α5 interface also emerged in our unbiased coarse-grained MD simulations. This finding suggests that the α4-α5 interface is the dominant dimer interface and that both models could be metastable states of the K-Ras4B active conformation. Moreover, in our MD simulations, we observe a dynamic water channel that can be attributed to the screening of the electrostatic interactions between the intermolecular salt bridges. Also, we analyze the allosteric connections in the K-Ras4B dimer interface and provide candidate amino acids for mutational studies and experimental testing. Finally, we simulate the K-Ras4B monomer in solution and on a model membrane using atomistic MD simulations. Binding site identification in representative structures extracted using Markov state modeling reveals that known binding pockets of K-Ras4B reported in the literature are identified only in the membrane-bound structure and not in the solution structure. Based on these results, we propose that modulating the protein-membrane interactions can be an alternative strategy for inhibiting K-Ras4B signaling.

## METHODS

### Building the K-Ras4B dimer structure with and without effectors on a model membrane

To our knowledge, the structure of a K-Ras4B dimer bound to its effectors has not been yet experimentally identified; however, biochemical and NMR data point to the α4-α5 interface being the most viable for KRAS dimerization, based on experimental evidence showing that the D154Q mutation abolishes both the WT and mutant G12D KRAS dimerization, without influencing its intrinsic GTPase activity, GEF or GAP sensitivity and C-Raf binding,^60^ and also on the recently-published molecular model of K-Ras4B based on the 4DSO PDB structure, using PRE-derived distances and a multibody docking protocol via HADDOCK 2.2.^61^ In our study, we modeled both structures proposed in Refs.^60, 61^ using the procedure outlined below to generate the starting structures.

Because no complete K-Ras4B dimer structure is available with the Raf effectors, information from multiple sources had to be integrated to produce the final model, including the full structure of the monomer K-Ras4B with its farnesylated HVR tail, the interface of K-Ras4B with the RBD domain of Raf, the missing loop connecting the RBD and CRD regions and the interface of RBD to CRD. Briefly, the full structure of the K-Ras4B dimer was generated using PDB ID 5VQ2^64^ as the core, and PDB IDs 4G0N^87^ and 2MSC^88^ were used to build the missing regions of K-Ras4B. To build the Raf[RBD-CRD] effector models we used structures with PDB IDs 4G0N and 1FAQ^15^, and subsequently coupled them to K-Ras4B to generate the final complex. A detailed description of modeling this structure is presented in Ref.^37^.

The structures reported in Ref.^61^ (PDB ID: 6W4E) are based on a monomer structure of mutant G12D K-Ras4B (PDB ID: 4DSO) complexed with GTPγS on a nanodisc, and the dimer was constructed using the multibody docking protocol available in the HADDOCK 2.2 program.^61^ However, given that the experimental setup of Ref.^61^ was based on WT K-Ras4B, we had to back-mutate G12D to glycine to directly compare to experimental results. We then prepared the first four frames using the corresponding module from the Schrodinger Suite in Maestro.^89^ The same protocol was used for attaching the Raf[RBD-CRD] domains as in Ref. ^37^.

A model membrane was then created with a lipid composition of 78% mol. DOPC, 20% mol. DOPS, and 2% mol. PIP2. Computational and experimental results highlight the importance of PIP2 and PS lipid interaction with the HVR and G-Domain leading to K-Ras4B localization on the plasma membrane justifying the choice for this model membrane.^23, 24, 82, 83, 90, 91^ For the model based on PDB ID 5VQ2, the K-Ras4B-Raf[RBD-CRD] complex model was placed on the membrane, and the farnesyl groups were inserted into the membrane and oriented vertically to the membrane plane adopting six different orientations (see Ref.^37^ for more information). For the 6W4E structure, the first four frames from the computational model that were generated based on the NMR restraints were chosen for simulation and the K-Ras4B-Raf[RBD-CRD] complex was placed on the membrane. Moreover, the monomer K-Ras4B was also simulated in solution as well as in contact with the model membrane with a lipid composition of 78% mol. DOPC, 20% mol. DOPS, 2% mol. PIP2 (see SI for more information).

The protocol regarding all monomer and dimer systems of this study, excluding the CHARMM-GUI membrane construction and equilibration, was performed using HTMD^92^ with its integrated psfgen builder, and the resulting simulations were carried out with ACEMD3 (see SI for more information).^93^ All systems were parameterized using the CHARMM36m force field.^94^ All simulations performed are listed in Table S1.

### Simulation analyses

The protein-membrane contacts were calculated using the contactFreq VMD^95^ tcl script. A contact is counted when two atoms are distanced below 5 Å, without accounting for hydrogens. The analysis of the salt bridges formed on the K-Ras4B dimer interface was carried out using the Salt Bridges plugin of VMD, according to which a salt bridge is considered to be formed if the distance between any of the oxygen atoms of acidic residues and the nitrogen atoms of basic residues are within the cutoff distance (default 3.2 Å). RMSD and RMSF calculations were performed for both K-Ras4B monomers (KR0, KR1) and residues 1-166, thus excluding the HVR region, which is expected to have increased fluctuations. Interaction energies of the K-Ras4B monomers were calculated using the NAMD Energy plugin of VMD and a cutoff of 12 Å. Cluster representatives of the G-domain of each simulation were calculated with the Gromos method and a cut-off of 0.2 Å. Correlation matrices were calculated using Pearson correlation in Python. The frequency of the inter-monomer interactions between the amino acids of the two monomers was calculated using the newcontacts VMD tcl script. The amino acids of the two monomers were considered to be in contact if the minimum distance between their heavy atoms was less than 5 Å. The HVR tail is omitted from the analysis. The exchange rate of water molecules was calculated using an in-house Python script. The 3D box was limited to the area between D154 and R161 residues and an exchange was counted when a water molecule entered from the top of the box and left from the bottom or the opposite. The amino acid movement correlations were calculated using the new version of the dynamical network analysis tool.^96^ The amino acids of the two monomers were considered to be in contact if the minimum distance between their heavy atoms was less than 4.5 Å for at least 75% of the simulation.

### Unbiased Coarse-Grained simulations

Coarse-grained MD simulations of the K-Ras4B dimer were performed on a 78% mol. DOPC, 20% mol. DOPS, and 2% mol. PIP2 membrane using Martini 3. In the starting conformation, two K-Ras4B monomers were placed on the membrane at a distance of 40 Å apart. An unbiased CG simulation with Martini 3 open beta force filed was performed for 20 μs. During the first 1.4 μs, the two monomers diffused separately on the membrane. Hence, a distance restraint was imposed between the center of mass of α4-α5 of the two monomers, which restricts the monomer’s separation to less than 25 Å. Because the helicity content of the α5 at the N-terminal residues of HVR (167-173) impacts the conformation of the K-Ras4B on the membrane and consequently the conformation of the dimer structure,^97^ the Martini ElNeDyn restraints corresponding to the α-helix in region 167-173 were removed from the topology. As a result, the extent of the α5 helicity on the HVR was not imposed by default.

### AlphaFold predictions

The dimer structure of K-Ras4B was predicted using AlphaFold v2.2, with and without using templates (by using current --max_template_date), generating five models. The full sequence databases were used for the predictions.

### Druggability analysis

Binding site prediction on monomer K-Ras4B structures was performed using SiteMap^98^ and FTMap^99^ with default parameters unless mentioned otherwise. The structures selected for binding site identification on the K-Ras4B monomer were isolated from the most conserved macrostates of the Markov state models (see SI for more information about the Markov state models). For each protein structure, a list of up to five potential binding sites was identified and analyzed (see the SI for more information).

The methodology scheme followed in this study is visualized in Figure 1.

**Figure 1.**
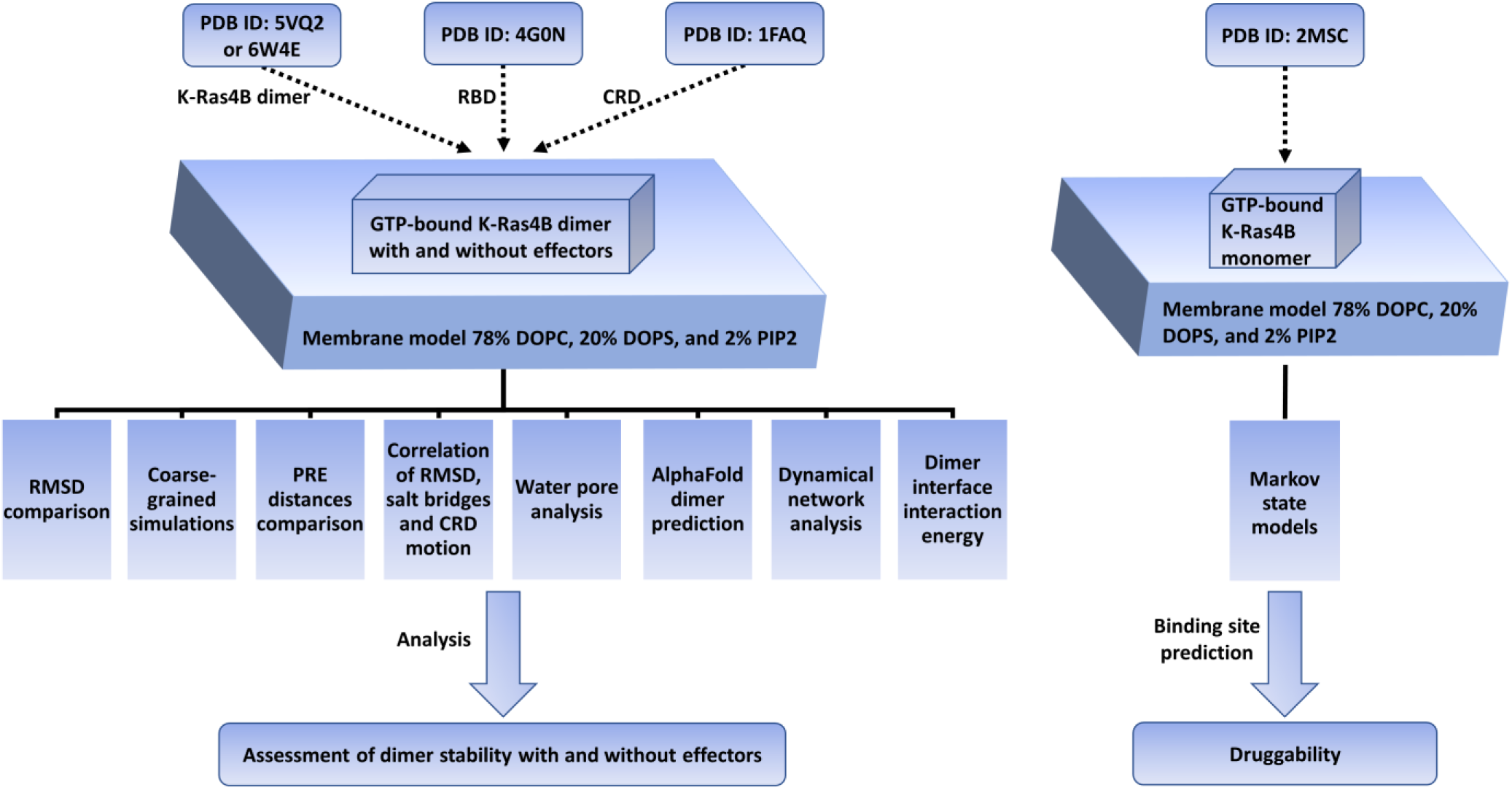
Workflow of system building and analyses.

## RESULTS

### Comparison of the two membrane-bound K-Ras4B structural models without effectors

We performed unbiased MD simulations for the dimer K-Ras4B structures based on PDB IDs 5VQ2 and 6W4E on a model cell membrane in the absence of effectors using 4 different replicas of 2 μs each to ensure reproducibility and proper statistics of the results. Henceforth, the simulations based on PDB ID 5VQ2 will be called Crystal #1-4 without effectors and the simulations based on PDB ID 6W4E will be called NMR frame #1-4 without effectors. First, we investigated the conformational flexibility of the two different dimer conformations in the absence of effectors, in association with a model cell membrane. We also monitored the stability of the intermolecular salt bridges formed on the two different dimer interfaces. As mentioned above, the K-Ras4B dimer interface based on the PDB ID 5VQ2 structure is stabilized by two intermolecular salt bridges between residues R161 and D154 belonging to two different monomers, while the K-Ras4B dimer interface based on the PDB ID 6W4E structure forms two intermolecular salt bridges between residues R135 and E168 of two different monomers (see Figure S3 for a schematic of the two different salt bridge cases).

The simulations based on the structure with PDB ID 6W4E have higher root mean squared deviation (RMSD) values compared to the simulations based on the structure with PDB ID 5VQ2 (Figure S4A). Visualization of the systems showed that in most cases K-Ras4B without effectors tilts relative to the membrane normal and lies towards the membrane plane initiating membrane contacts mainly with α2, α3, β6 K-Ras4B structural components (Figure S5 and Figure S6). We observed that K-Ras4B moves toward the membrane and interacts with the negatively charged lipid head groups. Intriguingly, for simulations 2 and 4 based on PDB ID 6W4E (NMR frame 2 and NMR frame 4), HVR and residues of β1 and β2 sheets, and α2 and α3 helices of one K-Ras4B monomer interact more frequently with the membrane, while the other K-Ras4B monomer maintains membrane contact only through the HVR tail (Figure S5 and Figure S6). A K-Ras4B dimer interface with an elevated monomer was also observed in MD simulations in Ref. ^70^, providing another pathway for the recruitment of Raf kinases. In all cases, the dimer complex based on PDB ID 5VQ2 without effectors shows enhanced stability with respect to the dimer based on PDB ID 6W4E.

The conformational flexibility of the K-Ras4B monomers within the dimer structure triggers an overall instability of the interactions in the dimer interface, including loss of the salt bridge between D154 and R161 (Figure 2, left panel) or R135 and E168 at the dimer interface (Figure 2, right panel). For the simulations based on PDB ID 5VQ2, a disruption of the α5-α5 dimer interface is observed. The system eventually becomes unstable and loses the salt bridges at the dimer interface except for one of the salt bridges in simulation 4 (Table S2). Similar behavior is also observed for the simulations based on PDB ID 6W4E, although one of the salt bridges is retained in simulation 2 (Table S2). These observations are in close agreement with reported studies suggesting that K-Ras4B gains dimerization capability only in the presence of additional factors, i.e., downstream effectors such as Raf.^27, 28^

**Figure 2.**
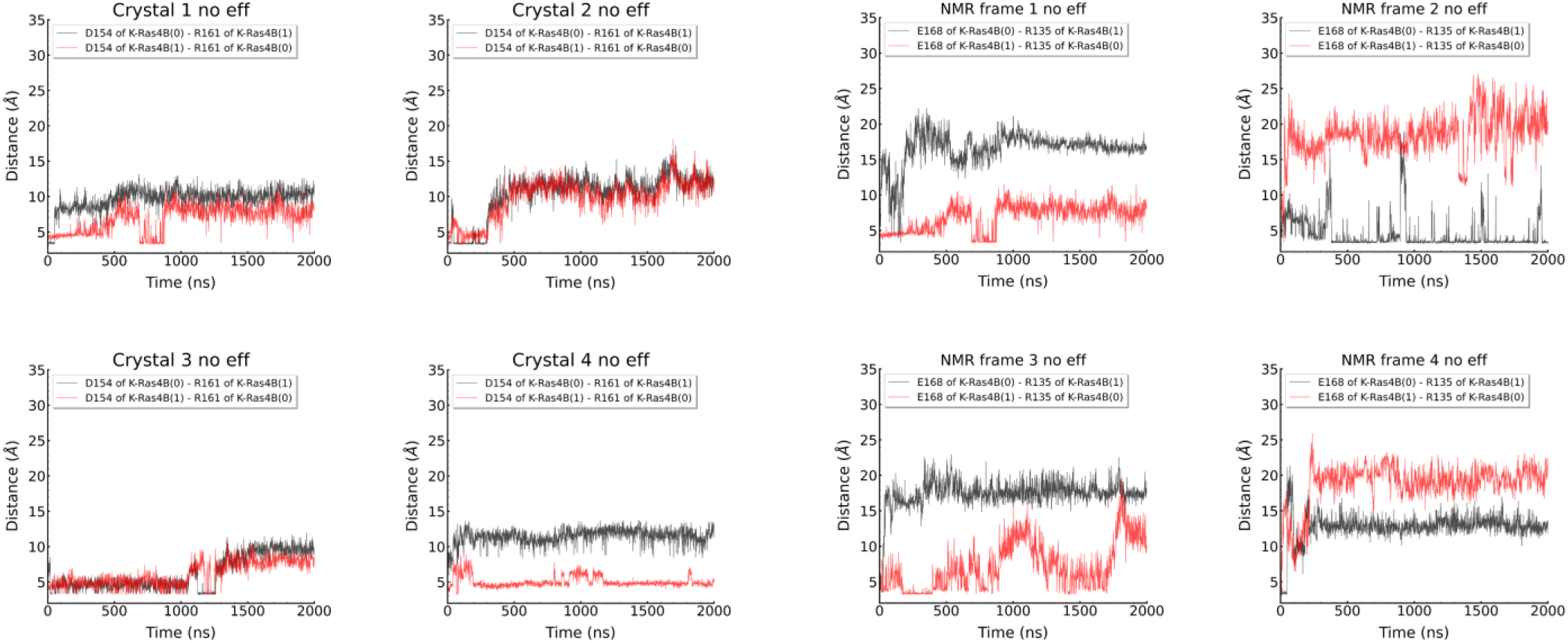
Distance of salt bridges between D154 and R161 for the simulations based on PDB ID 5VQ2 (left panel) and R135 and E168 for the simulations based on PDB ID 6W4E (right panel) without Raf[RBD-CRD] effectors.

### Comparison of the two membrane-bound K-Ras4B structural models with effectors

Next, we sought to investigate the conformational stability of K-Ras4B dimers in the presence of the Raf[CRD-RBD] effectors. We performed unbiased MD simulations for the dimer K-Ras4B structures based on PDB IDs 5VQ2 and 6W4E on a model cell membrane in the presence of effectors using 4 different replicas of 2 μs each to ensure reproducibility and proper statistics of the results. Henceforth the simulations based on PDB ID 5VQ2 will be called Crystal #1-4 with effectors and the simulations based on PDB ID 6W4E will be called NMR frame #1-4 with effectors.

The simulations based on the structure with PDB ID 6W4E have higher RMSD values compared to the simulations based on the structure with PDB ID 5VQ2 (Figure S4B). Overall, K-Ras4B dimers bound to Raf[RBD-CRD] demonstrate more conformational stability relative to dimers without Raf[RBD-CRD] (Figures S4), suggesting that the presence of the Raf effectors provides stability to K-Ras4B and limits K-Ras4B-membrane interactions by limiting the accessible K-Ras4B areas that may contact the cell membrane by steric hindrance. The percentage of interactions between the G-domain of K-Ras4B and the membrane validates this hypothesis because no K-Ras4B monomer tilting is observed, in contrast to some K-Ras4B systems without effectors (NMR frame 2 and NMR frame 4) (Figures S5-S8).

The K-Ras4B dimer stability is also reflected in the dimer interface interactions and particularly on the salt bridges formed between D154 and R161 of the two K-Ras4B monomers for the structure based on 5VQ2 except for replica 1 where the salt bridge breaks after ∼800 ns (Figure S9). On the contrary, the salt bridges formed between R135 and E168 for the structure based on 6W4E are unstable (Figure S10 and Table S2).

We then performed a correlation analysis for each simulation with effectors to observe which structural factors confer dimer stability. The crucial salt bridge distances (D154-R161 and R135-E168 center of mass distance), the distance of K148 of the CRD domains attached to each of the K-Ras4B monomers from the center of the membrane, and the RMSD from the initial PDB structure were taken into account. The calculations were performed for K148 of the two CRD domains and for both pairs of intermolecular salt bridges. Analysis of the correlation matrices reveals that the 6W4E systems and the Crystal 1 system that demonstrate higher RMSD than the Crystal 2, 3, and 4 systems, demonstrate also higher absolute correlation values (Figures S11 and S12). The driving force for RMSD stability for the 6W4E systems was mainly the crucial salt bridge distances, indicating that the high RMSD results from the separation of the salt bridges. Surprisingly, for the Crystal 2, 3, and 4 systems the crucial D154-R161 salt bridge exhibits minimum correlation to RMSD, while a moderate negative correlation is observed between RMSD stability and CRD mobility. The structures based on 6W4E are also influenced by the Raf[RBD-CRD] domain dynamics. These results are consistent with the findings of Ref.^68^, where increased CRD displacements were observed in the 5VQ2 interface albeit also with dimer disassociation. The relationship between HVR-CRD interactions and the stability of the dimer interface was also investigated but no correlation was found between the number of CRD-HVR contacts and the dimer interface disruption or the CRD leaving the membrane for any of the membrane dimer systems.

These observations suggest that the presence of Raf effectors may confer additional stability to the formation of stable K-Ras4B dimers, and provide insights into the CRD-membrane interactions influencing the dimer stability. The location of the Raf[RBD-CRD] in the structures with PDB ID 5VQ2 and 6W4E structures leaves the α4-α5 interface available for forming a dimer interface; moreover, the attachment of the CRD domain at the side of the K-Ras4B G-domain (Figure S1) could enhance dimerization by limiting the conformational space available for K-Ras4B dynamics.

### Relative stability of the two dimer structures

The experimental data that reveal the α4-α5 interface as the most promising for dimerization describe two closely related conformations.^60, 61^ Undoubtedly, both of these conformations could be relevant to the dimerization procedure, since K-Ras4B exhibits high flexibility on the membrane. However, characterizing the thermodynamic stability of the two different conformations would require longer-scale computations utilizing biased MD. Although the latter is a subject for future work, here we assessed the relative stability by calculating the enthalpic terms of the protein-protein interactions. Interaction energies of the K-Ras4B monomers were calculated taking into account electrostatic and vdW energies as previously suggested in the literature.^100^ From Figure 3, it is evident that the dimer structures based on PDB ID 6W4E establish stronger interactions. However, in terms of RMSD (Figure S4), RMSF (Figure S13), and distances between the center of mass of amino acids D154-R161, R135-E168, and Q131-R161 (Figures S14-S17), the 5VQ2 systems shows enhanced stability compared to 6W4E systems throughout the 2 μs simulations. Moreover, in Figure 3 we observe that the 6W4E systems have different interaction energies between them, while the interaction energies for most of the 5VQ2 systems are similar. This difference can be attributed to the fact that the 6W4E frame systems have different initial coordinates, while the 5VQ2 systems have the same initial coordinates with different initial velocities.

**Figure 3.**
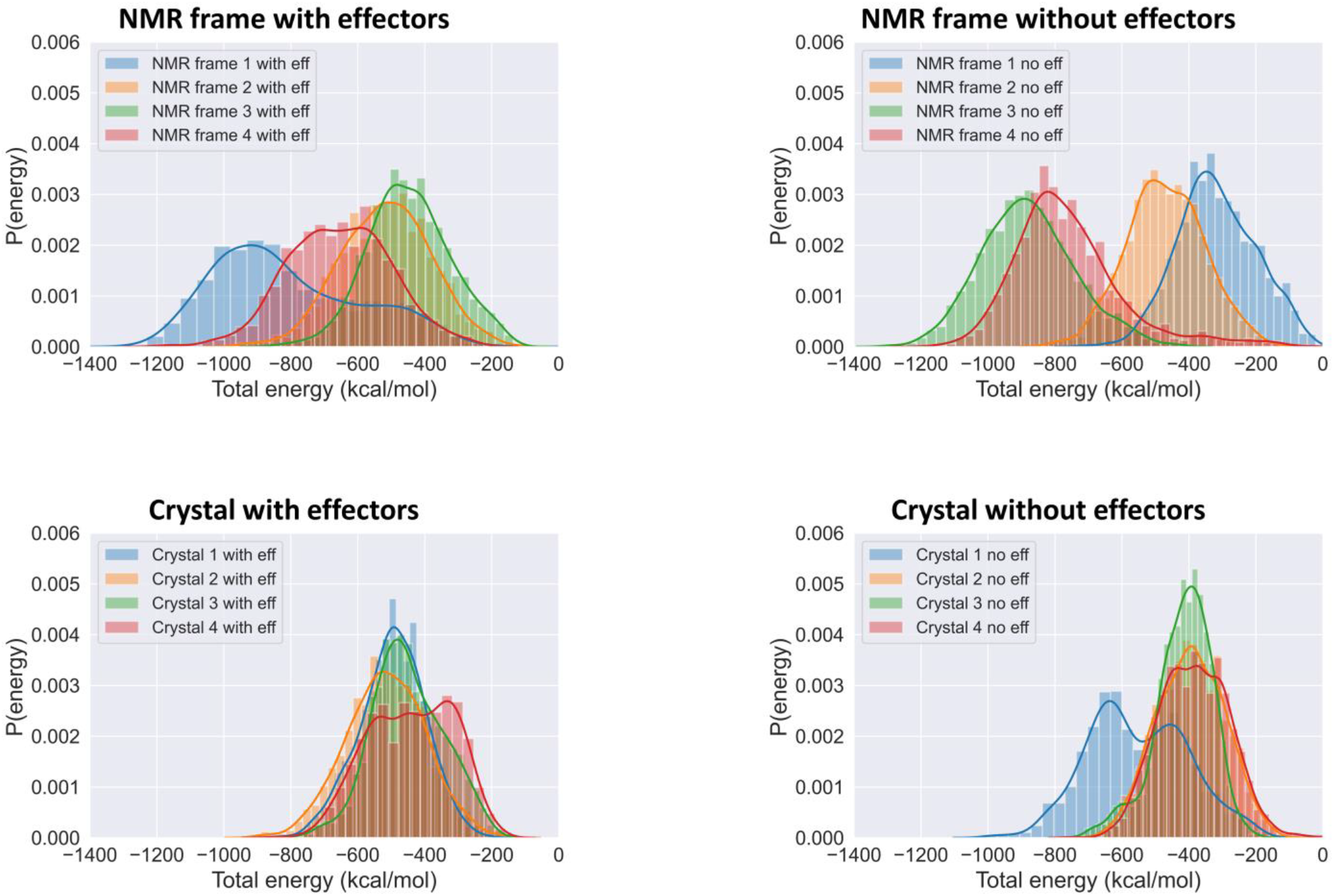
Normalized distributions of the interaction energies between two monomers in the simulations with the structure based on PDB ID 6W4E (upper panel, NMR frame with and without effectors) and the structure based on PDB ID 5VQ2 (Crystal with and without effectors, lower panel).

### Inter-monomer interactions of the two dimer structures

Analysis of the frequency of the inter-monomer interactions between the amino acids of the two K-Ras4B monomer G-domains showed that most interactions are formed around the α4 and α5 regions for the 6W4E and 5VQ2 systems (Figure S18-S21). Moreover, we observe interactions between the β2-β3 loop (D47, G48, and E49) and the β6 sheet, α4, and α5 regions for both 6W4E and 5VQ2 systems, suggesting that the region D47-E49 might be important for dimer stability. Interestingly, in the first 6W4E system without effectors (NMR frame 1 no eff) a significant conformational change occurs (Figure S4A), where the D47-E49 regions between the two monomers come in contact and interact until the end of the simulation.

The distance of the center of mass of critical dimer interface amino acids D154-R161, R135-E168 and Q131-R161 were plotted against the simulation time for both K-Ras4B monomers (Figures S14-S17). The R135-E168 salt bridge is present in both interfaces with a quick shift towards the 5-7 Å distance region for the 6W4E structures (Figures S14, S15). The D154-R161 interaction was not observed for the 6W4E systems. In most cases, the Q131-R161 distance deviated slightly or preserved the initial distance.

Furthermore, we searched for allosteric connections in the 5VQ2 and 6W4E interfaces based on the generalized correlation of amino acid motions using the new version of the dynamical network analysis tool.^96^ Table S3 shows the generalized correlation values for the K-Ras4B dimer interface of all 5VQ2 and 6W4E trajectories with and without effectors. The allosteric communication in the 5VQ2 interface is mainly mediated through the regions around R161-K165, R135-E143, and D47-G48. For the 5VQ2 system without effectors, significant movement correlations were found for the R135-E168 interface pair (0.64 generalized correlation), although the R135-E168 salt bridge is important for the 6W4E interface stability. For the 6W4E systems, in many cases, the HVR is found to be implicated in movement correlations between the two monomers, but in general, no common correlated interface pairs were found between the different 6W4E simulations. This observation can be attributed to the fact that the 6W4E systems had different initial conformations since they were retrieved from different structures of the 6W4E NMR ensemble, or because the 6W4E interface is unstable providing the opportunity to explore different interface orientations.

### Cross-RMSD analyses

Next, we investigated whether the simulations of the 5VQ2 systems reach the 6W4E interface and the opposite. We measured the RMSD of the G-domain of the 5VQ2 simulations taking as a reference structure the first structure of the 6W4E NMR ensemble, and the RMSD of the 6W4E simulations taking as a reference structure the 5VQ2 crystal structure (Figure S22). The 5VQ2 systems which are more stable than the 6W4E systems display a stable RMSD from the 6W4E structure of approximately 6 Å, while the 6W4E systems display an increase in the RMSD from the 5VQ2 structure up to 14 Å.

To further analyze the RMSD between the two interfaces of interest, we acquired cluster representatives for each of the simulations of the G-domain. Cross-RMSD analysis of the 5VQ2 and 6W4E systems was performed using the cluster representatives of the most populated clusters to directly compare the tendency toward the two proposed interfaces (Figure S23 and Table S4). The cluster representative of Crystal 1 simulation (5VQ2, with effectors) compared to NMR frame 3 simulation (6W4E, no effectors) has the minimum cross-RMSD observed (5.8 Å, Figure S24). This RMSD value can be attributed to the conformational change of Crystal 1 with effectors system (Figure S4B and Figure S9). In Crystal 1 with effectors simulation, two major clusters were found: the first cluster contains the conformations from the beginning of the simulation up to ∼800 ns, and the second and most populated cluster contains the conformations from ∼800 ns up to the end of the simulation.

### Comparison with experimental PRE restraints

In Ref.^61^, two distinct K-Ras4B dimer structures (GDP-bound or bound to the non-hydrolyzable GTP analogue GTPγS) anchored on a nanodisc membrane were determined by Paramagnetic Relaxation Enhancement (PRE) NMR experiments. PRE has long been recognized as a method for providing long-range distance information that can complement NOE restraints, which are limited to distances of up to ∼6 Å.^101^ To stabilize K-Ras4B on the membrane, they employed a maleimide-conjugated KRAS (MC-KRAS) combining it with the farnesylated KRAS (FP-KRAS). PRE effects on isotopically tagged atoms (probes) on MC-KRAS before versus after the spin labels (nitroxide tags attached on FC-KRAS) were quenched and assessed by measuring the relative reductions intensity of the paramagnetic to diamagnetic peaks (Ipara/Idia). These values were converted into the ^1^H transverse PRE rate (^1^H-Γ2, spin-spin PRE relaxation time), and examined proton distances were calculated. To account for errors in the estimate of ^1^H-Γ2, the authors set the distance restraint bounds to ± 3 Å from the experimentally-measured distance. A direct comparison of the PRE distances presented in Ref.^61^ versus our simulations was performed and evaluated for each system. 48 intermolecular atomic distances were calculated retaining the same order as presented in Lee et al.^61^ with numbers 1-15 corresponding to the top of the interface, 16-40 the α4-α5 intermolecular distances, and 41-48 the HVR region (Figures S25).

The experimentally measured PRE distances were on average lower than the simulated distances in simulations “NMR frame #1-4” and “Crystal #1-4” with and without effectors (Figure S26 and Table S5). A reason for the lower experimental PRE distances compared to the MD simulations measured-PRE distances could be the stability that the maleimide group confers to K-Ras4B in Ref.^61^. The maleimide group attaches irreversibly to the membrane, which could artificially lower the PRE distances and which is absent in our simulations. We additionally performed a time series analysis of the PRE distances for each simulation in the production run (Figures S27-30) and the difference of simulated PRE distances between the last and the first frame of the simulation time was calculated on all 48 pairs, and the mean and std values of these differences were measured for all 48 PRE distances of each replica simulation for each system. For the simulated structures based on PDB ID: 6W4E, a shift toward higher PRE distances was observed (2.3 ± 3.2 Å, 2.8 ± 4.5 Å, 10.3 ± 8.7 Å, and 4.6 ± 3.7 Å for simulations NMR frame #1-4 with effectors and 8.4 ± 11 Å, −0.9 ± 3.2 Å, 2.6 ± 3.8 Å, and 4.5 ± 4.9 Å for NMR frame #1-4 without effectors), while the structures based on PDB ID: 5VQ2 remained stable in most simulations (4.6 ± 3.5 Å, 1.7 ± 5.9 Å, −2.1 ± 5.4 Å, and 0.8 ± 2.7 Å for the simulations of Crystal #1-4 with effectors and 0.1 ± 1.9 Å, 0.5 ± 4.6 Å, 1.2 ± 5.7 Å, and −1.2 ± 5.3 Å for simulations of Crystal #1-4 without effectors).

These results suggest that the systems based on PDB ID 6W4E on the model membrane, alter their KRAS dimer conformation with respect to the PRE NMR study of Lee et al^61^, whereas the simulations based on PDB ID 5VQ2 preserve the KRAS dimer domain internal distances.

### A dynamic water channel is observed in the 5VQ2-based K-Ras4B dimer model

On the monomeric K-Ras4B crystal structure (PDB: 6GOF),^41^ as well as on our monomer K-Ras4B simulations (see the SI), it is observed that D154 and R161 of the K-Ras4B α5 helix interact through the formation of an intramolecular salt bridge (Figure S31). However, this interaction is replaced in the dimer K-Ras4B simulations by two intermolecular salt bridges at the dimer interface between D154 of one K-Ras4B monomer and R161 of the other monomer. In our simulations, we consistently observe the existence of a stable water channel that flows between the R161 residues of the two monomers, separating the two intermolecular salt bridges (Figure 4a). This water channel mediates intermolecular interactions between R161 and D154 while preventing the formation of a direct intramolecular salt bridge between R161 and D154.

**Figure 4.**
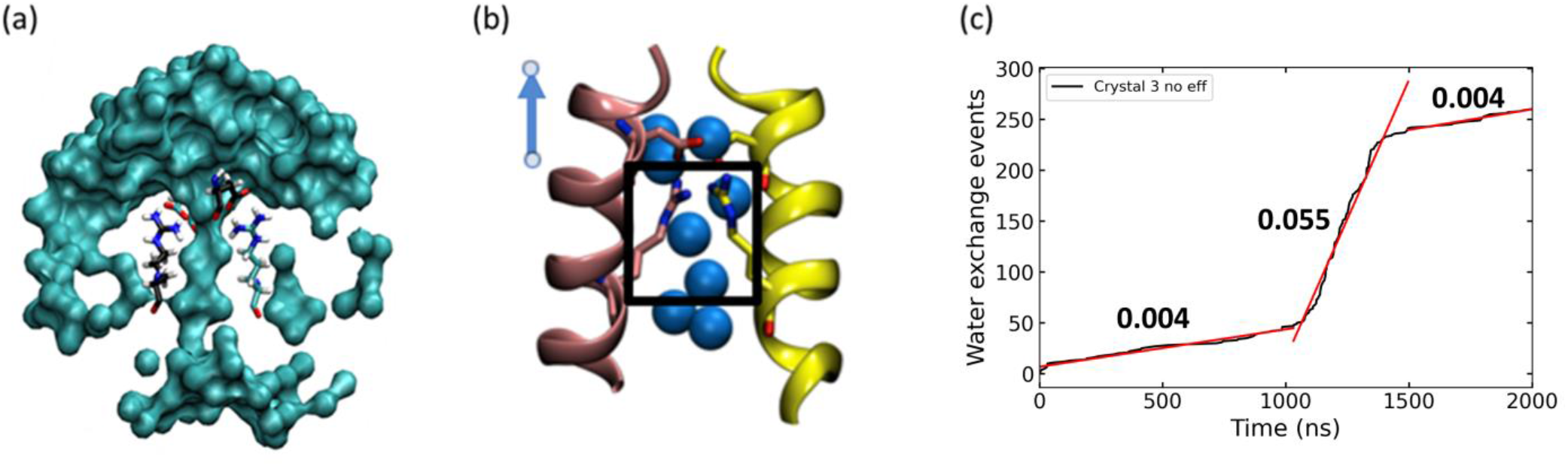
(a) Water channel observed at the K-Ras4B dimer interface. (b) Water exchange events in the vicinity of the R161-D154 salt bridge. (c) Water exchange event counts within the water channel for NMR frame 3 without effectors. Three different water exchange rates were calculated as the slope of the corresponding time-dependent events.

To quantify whether there is any water flow from this channel, we calculated the rate of the exchange of water molecules. The 3D box was limited to the area between D154 and R161 residues (Figure 4b). This procedure then counts water exchanges in this pore, which can result in different water exchange rates counted as the slope of the corresponding time-dependent events (Figure 4c); the larger the slope, the faster the exchange takes place. We observe that there is an anti-correlation of the water exchange events to the salt bridges stability at the Kras4B interface (R161-D154). As expected, the more stable the salt bridges between the two monomers are (which is achieved in 5VQ2 more than 6W4E models), the less water exchange takes place (Figure S32). Furthermore, K-Ras4B dimers with effectors appear to have fewer exchanges compared to the no-effector systems, effectively preserving the water channel. When the water channel opens, the water exchange events are that of bulk water. The functional role of this water channel can be attributed to the screening of the electrostatic interactions between the intermolecular salt bridges R161 and D154 (Figure 4a,b). In contrast to the 5VQ2 systems, for the structures based on PDB ID 6W4E, such a water channel/pore is not observed in our simulations, being replaced by a water interface (Figure S33).

### Coarse-Grained dimer simulations

Atomistic simulations provide a rigorous way to study biological processes at the molecular level. However, the spatiotemporal bandwidth of atomistic MD simulations hinders studying processes such as spontaneous dimerization of proteins. The use of coarse-grained (CG) simulations could provide insights into the K-Ras4B dimerization. In a previous study,^102^ we have identified that Martini 3 open beta force field identifies in all cases the near-native dimer complexes as minima in the free energy surface, albeit not always as the lowest minima. Thus, we chose to perform CG K-Ras4B/Raf[RBD-CRD] complex simulations to identify dimer structures at the α4-α5 interface.

An RMSD analysis of the unbiased coarse-grained simulations with respect to the 5VQ2 and 6W4E PDB structures (starting from a structure in which the α4-α5 helices of each monomer are 25 Å apart) showed that the system can reach a conformation as close as ∼5.5 Å from 6W4E, but the α5 helices never associate to achieve a 6W4E-like structure (Figure S34). On the other hand, the interactions at the α4-α5 interface of 6W4E are strong enough and the RMSD reveals a stable conformation for ∼10 μs. The dimer conformation after the 10 μs was backmapped to atomistic resolution (Figure 5a,b). The α5 helices are in a parallel conformation resembling the 6W4E structure. The R161, D154, and Q131 coordination however do not correspond to the 6W4E structure except for the interaction of one pair of the intermonomer D154-Q131. The relative positioning of the β2-β3 of the experimentally obtained structures and the simulated one reveal that a rigid rotation of the two monomers on the XY plane would suffice to obtain the two structures (Figure 5c,d).

**Figure 5.**
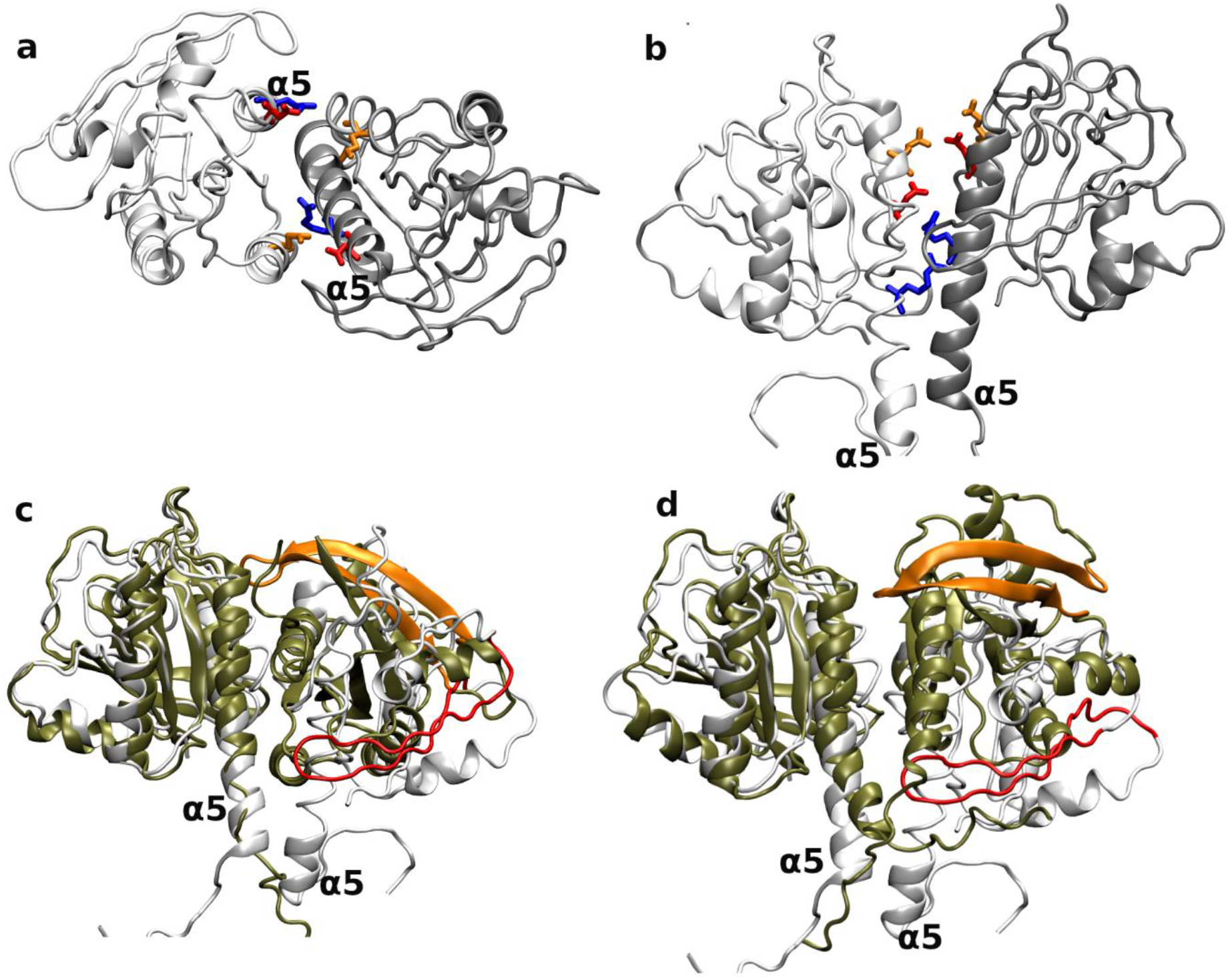
(a) and (b) The atomistic resolution structure that corresponds to the final frame of a 10 μs unbiased CG simulation. The D154 (red), R161 (blue), and Q131 (orange) are shown in licorice. The backmapped CG structure (white) was aligned to the 5VQ2 (c) and 6W4E (d) structures leading to Cα RMSDs of 11.45 Å and 7.45 Å, respectively. The crystal and NMR structures are colored in tan while the β2-β3 are colored in orange. The corresponding β2-β3 obtained from the simulations are colored in red. The Raf[RBD-CRD] domains are not shown for clarity.

The final frame of the 10 μs unbiased CG simulation showed a different conformation compared to 5VQ2 and 6W4E which could potentially be a metastable state connecting the monomers to the dimeric state as it also implicates the α4-α5 interface. As such, it was directly compared to all our simulated systems using RMSD (Figure S35). Only the K-Ras4B G-domain was taken into account for these comparisons as the HVR and the effector exhibit a high degree of flexibility. The time series of RMSD comparing the CG final frame structure to the dimer atomistic simulations show that most 5VQ2 systems are stable without a preference for the CG final frame structure (Figure S35). On the contrary, the 6W4E structures reached a minimum RMSD of ∼3.3-5.5 Å compared to the CG G-domain, being stable throughout the simulations except for NMR frame 3 with effectors and NMR frame 1 without effectors simulations. If we exclude these NMR simulations, the dimer interface proposed by this CG structure is closer to the structure with PDB ID 6W4E, with an average RMSD varying between ∼4.8-7.1 Å in contrast to the ∼10-12.2 Å deviation presented in the 5VQ2 dimers (Figure S35). These findings suggest that the dimerization process could be mediated by a structure resembling the CG structure leading to a structure with a dimer interface of PDB ID 6W4E, while the structure that corresponds to PDB ID 5VQ2 could assemble slower.

### K-Ras4B dimer prediction using AlphaFold

With the advent of AlphaFold2, the 3D structure of proteins can be predicted from the amino acid sequence with high accuracy.^103^ Moreover, the new version of AlphaFold, called AlphaFold-Multimer, is specifically trained for the prediction of protein complexes and the prediction of the protein-protein interfaces.^104^ Therefore, we utilized AlphaFold-Multimer to predict the K-Ras4B dimer with and without effectors and with and without the HVR tail. However, in the predictions without effectors, although the K-Ras4B monomers were predicted with high pLLDT (Figure S36A), the predicted alignment error between the two monomers was below average (Figure S36B), suggesting that the dimer interface was predicted unsuccessfully. For the predicted structures of K-Ras4B dimer with effectors, the two K-Ras4B monomers are not in contact and the complex is formed by the Raf[RBD-CRD] effectors.

### Druggability analysis of K-Ras4B

Despite the numerous attempts over the decades, targeting KRAS remains a challenging task. The difficulty in drugging KRAS is mostly attributed to the high affinity of KRAS for its GTP and GDP substrates, as well as to the lack of deep hydrophobic pockets on the protein surface.^43^ Strategies focusing on targeting the KRAS GTP pocket by inhibition of its catalytic site are hindered by the picomolar affinity of KRAS for its substrates.^44^ One such success story was the design of a GDP analogue, inhibiting KRAS signaling by interacting covalently with C12 on the active site of the G12C KRAS mutant.^105^ Ever since, alternative approaches have been employed, focusing mainly on targeting allosteric sites on KRAS. Several studies reveal the intrinsic allosteric nature of KRAS^106-110^ providing insightful ideas for targeting this oncoprotein via modalities other than active site inhibition. Such an approach was implemented by the Shokat group in 2013, targeting G12C GDP-bound KRAS mutant with small inhibitors that bind covalently on a pocket located over the Switch II loop, impairing GTP and Raf binding.^111^ This neighboring KRAS active site pocket, called the Switch II pocket, encompasses residues belonging to the β1 sheet, the α3 helix, and also to the Switch II loop. The Switch II pocket was also studied by Patricelli *et al*., who discovered G12C GDP-bound KRAS selective inhibitors with promising cellular activity.^45^ These inhibitors suppress G12C KRAS mutant signaling and tumor cell growth,^46^ by inhibiting downstream signaling. Recently, a potent G12C covalent inhibitor named sotorasib (AMG510) obtained FDA approval and became the first therapy to directly target the KRAS oncoprotein.^47^ However, the aforementioned inhibitory strategies are mutant-specific and target only the G12C KRAS, a mutation not frequently observed in the clinic, as it is present in the lung but not in pancreatic and colorectal cancers.^3^ A non-mutant-specific approach was employed by Fesik and colleagues, who targeted a conserved binding site in the WT, G12D, and G12V KRAS, spotted between Switch I and II, with small molecules that inhibit the activation of Ras by SOS.^38^ Furthermore, recently, this shallow Switch I-II pocket was targeted in G12D KRAS with nanomolar affinity inhibitors that disrupt the interaction of inactive KRAS with SOS, and active KRAS with SOS and its effectors,^72^ and also in G61H mutant KRAS_169_ by Cruz-Migoni *et al*.^41^ Additionally, Gorfe and colleagues employed a computational approach and predicted four allosteric binding sites on KRAS, based on the conformational transitions between the GDP- and GTP-bound protein.^109, 110^ Two out of the four pockets, pocket p1 (located between the β1, β2, β3 sheets and helix α2) and pocket p3 (located between helix α5, loop 7 and loop 9), were characterized as conserved amongst the tested conformational ensembles and were targeted with small inhibitors.^110, 112^ Finally, several studies have focused on targeting the interaction of KRAS with its effectors, such as Raf^113, 114^ or PDEδ^42^ as well as on disrupting KRAS membrane localization.^115-117^

In our study, we performed binding site identification on K-Ras4B monomer structures resulting from our simulations, using SiteMap^98^ and FTMap.^99^ Specifically, the structures used for the binding site identification on WT K-Ras4B GTP-bound monomer on the membrane, WT K-Ras4B GTP-bound monomer in solution, and WT K-Ras4B GDP-bound monomer in solution were sampled from the most populated macrostates resulting from the Markov state models (for more information see the SI and Figure S37d). For the membrane-bound monomer GTP-WT structure, SiteMap identified three binding sites, two of which are located near the Switch regions and one on the K-Ras4B-membrane interface (Figure 6). FTMap also predicted the same binding sites as mentioned above for SiteMap.

**Figure 6.**
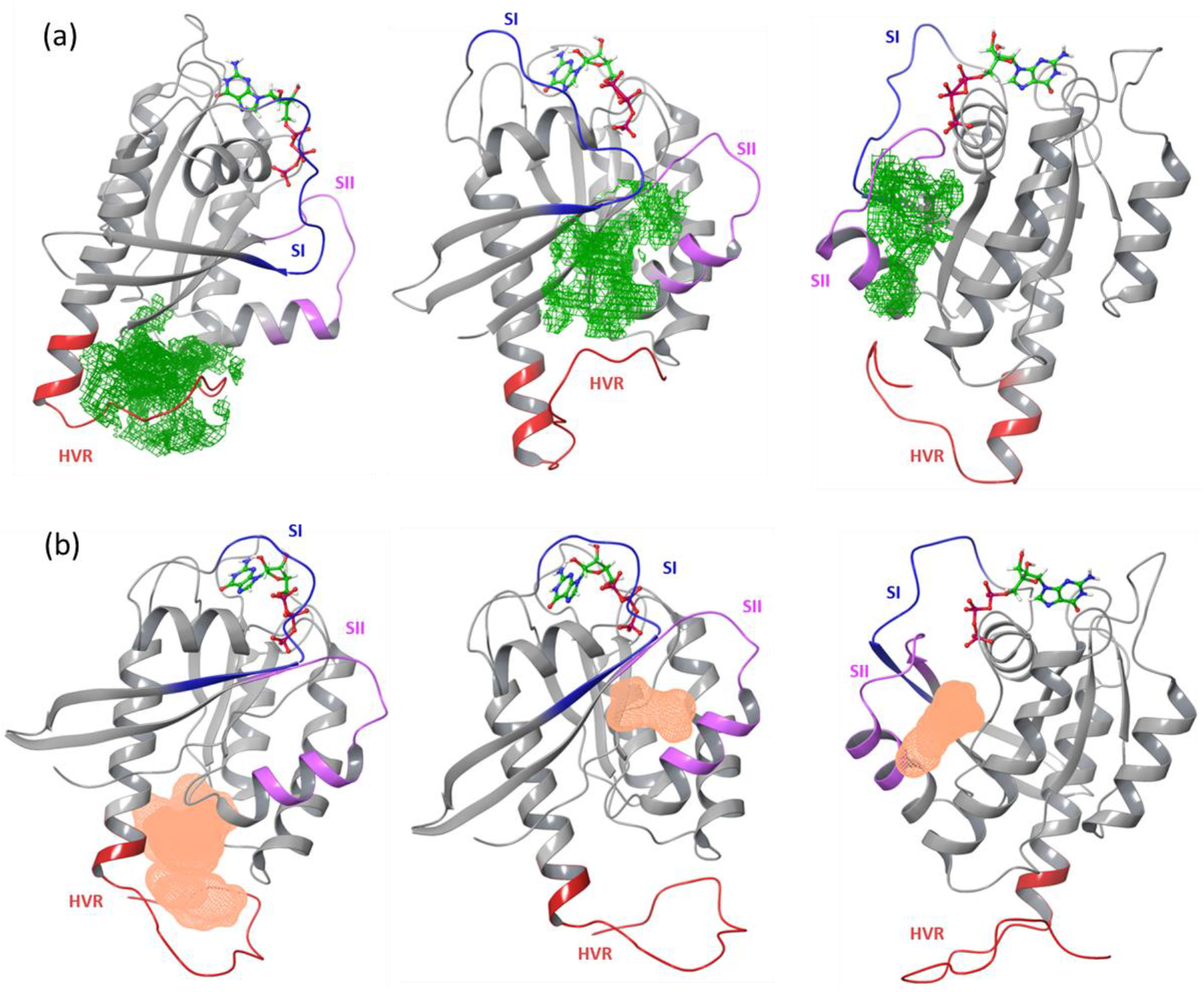
Binding sites on WT K-Ras4B GTP-bound monomer on the membrane, identified by (a) SiteMap and (b) FTMap. K-Ras4B monomers are presented in grey ribbons. Predicted binding sites in (a) are presented in green mesh and in (b) in pink mesh. Residues belonging to the Switch I region are colored blue, residues in Switch II are colored purple, residues in HVR are colored red, while GTP is in stick representation.

This analysis led to the identification of a binding site on the protein-membrane interface, located near residues of the β1 sheet, the loop before β4, α3 helix, the loop before β5, the α5 helix, and the N-terminal part of HVR. This binding site has a high SiteScore value of 0.924 and a druggability score (DScore) of 0.817. Another binding site with a good druggability score was also identified, located amongst the Switch I and II regions (see SI for more information). This site corresponds to the Switch I-II pocket mentioned in literature^38, 41, 72^ with residues belonging to the β1 sheet, Switch I and II loops of K-Ras4B. The third binding site identified corresponds to the Switch II pocket^45, 111^ and is mostly comprised of residues on the Switch II region, as well as residues from the β1 sheet, α2 and α3 helix (SiteScore = 0.982 and Dscore = 0.980). Notably, the identified binding pockets resemble the pockets identified in the literature^41, 112, 118^ only for the membrane-bound structures but not the solution structures. This observation may suggest that upon protein-membrane binding, the protein conformational ensemble alters toward the state that includes these binding sites, however, the absence of these binding sites in the solution structures may be due to limited sampling of the conformational space in our MD simulations, the starting protein conformation, and the protein conformation that was sampled from the distribution of the most populated macrostate in the MSMs. These results imply that either extensive sampling should be performed in solution to obtain the macroscopic picture observed in biophysical studies or the molecular simulations should be performed in the presence of the membrane which can reveal these pockets in less computational time.

## Discussion and Conclusions

The continuous quest for K-Ras4B targeting therapeutic strategies led to a novel approach for targeting K-Ras4B dimers based on cumulative evidence that membrane-bound K-Ras proteins, including K-Ras4B, form dimers or even nanoclusters in order to activate signaling pathways, with the dimer proposed to be the basic clustering unit.^20, 24, 48-53^ Dimerization is critical for the activity of WT K-Ras as well as for the oncogenic ability of mutant G12D K-Ras.^60^ Ras dimers have been observed in the biological assembly of an X-ray crystal structure^64^ and NMR-inspired structures,^61^ and different dimer interfaces have been suggested using both computational and experimental methods for members of the Ras family,^52, 53^ but still the exact structure of the KRAS dimer has not been experimentally resolved in the presence of its effector proteins.

Here, we perform atomistic MD simulations to compare for the first time two recently-proposed experimental dimer K-Ras4B structures in the presence and absence of the Raf effectors on a realistic model anionic cell membrane including the crucial lipids for K-Ras4B function PIP2 and DOPS: one based on biochemical evidence compatible with PDB ID 5VQ2 and one based on NMR experiments modeled as PDB ID 6W4E. An NMR structure of a monomeric K-Ras4B in the presence of Raf[RBD-CRD] in a nanodisk which recently appeared in the literature,^119^ was not compatible with a membrane orientation of having both HVRs positioned inside the membrane (Figure S38), indicating that K-Ras4B may adopt multiple configurations on the membrane surface in its monomeric state.

Based on MD simulations of the two structures with and without the Raf[RBD-CRD] effectors, we find that in most cases the RMSD of both dimers relative to their respective PDB structures showed a steady increase and that K-Ras4B without effectors tilts relative to the membrane normal and lies towards the membrane plane. This conformational flexibility of the K-Ras4B monomers triggers an overall instability of the interactions in the dimer interface, including the loss of crucial salt bridges reported in the experimental studies as major structural determinants of these conformations. On the contrary, in the presence of the Raf[RBD-CRD] effectors, K-Ras4B dimers bound to Raf demonstrate more conformational stability relative to dimers without Raf, suggesting that the presence of the effector provides stability to K-Ras4B and hampers K-Ras4B-membrane interactions by limiting the accessible K-Ras4B areas that may contact the cell membrane by steric hindrance. Correlation analysis of structural factors conferring dimer stability revealed that the driving force for RMSD stability was either the crucial salt bridges distances for the systems based on the 6W4E structures or a weak to moderate correlation of CRD distance from the membrane for the 5VQ2 structures. We thus argue that in order for KRAS to stabilize on the cell membrane, it needs the presence of effectors that may act as steric locks to its membrane-anchored conformation.

In terms of RMSD, RMSF, and distances of the amino acids D154-R161, R135-E168, and Q131-R161, the 5VQ2 systems show enhanced stability compared to the 6W4E systems. It has been argued that the nanodisc prevents the K-Ras4B homodimer from populating all possible orientations, resulting in biased dimer interfaces.^70^ In our unbiased coarse-grained MD simulations starting from the dissociated K-Ras4B complex we found that the α4-α5 interface is indeed the dominant dimer interface, reaching a conformation as close as ∼5.5 Å from 6W4E. This finding suggests that the dimerization process could be mediated by a structure resembling the coarse-grained final frame structure and that both 5VQ2 and 6W4E could be metastable states of the K-Ras4B active conformation. More advanced computational techniques, such as metadynamics and adaptive MD simulations could provide further insights into discovering the K-Ras4B dimer interface.

Moreover, recent studies propose dimer interfaces that allow one K-Ras4B monomer to interact with the GTP of the other K-Ras4B monomer.^69, 70^ One of these interfaces was also supported by mutational studies and FRET analysis,^69^ where D30R, E31R, E62R, and K147D mutations that are close to GTP disrupted the FRET signal of K-Ras4B dimer interactions, but this disruption can be an effect of allosteric modulation impaired from these mutations. D154Q and R161E mutations which are in the α5 helix were also experimentally found to disrupt K-Ras dimer interactions,^60, 69^ although recently it was shown that D154Q K-Ras4B mutant can dimerize in 3 different assays.^120^ Our analysis for allosteric connections in the dimer interface resulted in amino acids that might be significant for the dimer stability, such as amino acids D47, D105, R135, E143, and K165. These amino acids may act as promising candidates for mutational studies and experimental testing.

Furthermore, the validity of our simulations is checked upon comparison with the experimental PRE NMR restraints. Our MD simulations measured-PRE distances were higher compared to the experimental PRE distances, especially in the HVR region. The lower PRE distances in the NMR restraints can be attributed to the stability that the maleimide group confers to K-Ras4B, which is absent in our simulations. The maleimide group attaches irreversibly to the membrane, which could artificially lower the PRE distances.

Finally, we performed binding site identification on K-Ras4B GTP-bound monomer structures resulting from Markov state model analysis. Our analysis led to the identification of pockets of the K-Ras4B monomer that have been previously observed and studied in the literature and were absent in the solution structures. It is possible that the K-Ras4B-membrane binding is a conformational selection binding mechanism that favors the K-Ras4B conformation that includes these binding sites, although the limited sampling of the conformational space in our MD simulations of the solution structures may be another explanation for not finding these binding sites in the solution structures. In the future, exploring the conformational landscape of mutant K-Ras4B, e.g., G12D, would aid the drug design of K-Ras4B by finding binding sites in the mutant form that are absent in the WT form and may allosterically inhibit the mutant function.

In conclusion, we found that Raf[RBD-CRD] effectors provide stability to the K-Ras4B, acting as a steric lock to its membrane-anchored conformation. Calculation of protein-protein interaction energies showed that the structure based on PDB ID 6W4E forms more stable interactions than the model based on PDB ID 5VQ2, however, the 5VQ2 structure shows enhanced stability in terms of RMSD and RMSF throughout the 2 μs of the study. Moreover, the a4-a5 interface also emerged in the CG simulations. These findings suggest that both of these structures are viable and that the α4-α5 interface is the most probable. Binding pocket analysis in different KRas-4B structures showed that binding pockets of KRas-4B located in the effector binding region and the membrane interacting area of the protein are identified only in the structure that is membrane-bound, but not in the solution K-Ras4B structure. This finding is expected to significantly aid researchers working with K-Ras4B drug design. Based on these results, we propose that modulating the protein-membrane interactions can be an alternative strategy for inhibiting K-Ras4B signaling.

## ASSOCIATED CONTENT

### Supporting Information

Detailed methods, data analyses with associated figures, and tables (PDF).

## AUTHOR INFORMATION

### Author Contributions

JM, CVV, JD, ZC conceived and designed the study, AC, IA, SD, CL, AT performed the simulations, analyzed the data, and wrote technical details. AC, IA, ZC, AT wrote the manuscript.

### Notes

The authors declare no competing financial interest.

## ACKNOWLEDGEMENTS

This research work was supported by the Hellenic Foundation for Research and Innovation (H.F.R.I.) under the “First Call for H.F.R.I. Research Projects to support Faculty members and Researchers and the procurement of high-cost research equipment grant” (Project Number: 1780) awarded to ZC and the Operational Programme «Human Resources Development, Education and Lifelong Learning» in the context of the project “Strengthening Human Resources Research Potential via Doctorate Research” (MIS-5000432), implemented by the State Scholarships Foundation (IKY) to AC. We acknowledge computational time granted from the Greek Research & Technology Network (GRNET) in the National HPC facility – ARIS under project IDs pr008031_thin/KRAS-HVR, pr006033/kras, pr007019/kras. We acknowledge that the results of this research have been achieved using the DECI-16/KrasMem resource Kay based in Ireland at ICHEC with support from the PRACE aisbl. We acknowledge PRACE for awarding us access to Marconi100 hosted by CINECA, Italy.

