## Supplementary information for "Evaluation of K-Ras4B dimer interfaces and the role of Raf effectors"

### METHODS

The following protocols were employed for the preparation of the K-Ras4B membrane-bound monomer and dimer structures with Raf[RBD-CRD] effectors.

#### Membrane models

Two membrane models were obtained from the bilayer membrane builder of CHARMM-GUI.[1] The first membrane was constructed for the monomer simulations with a lipid composition of 78% DOPC, 20% DOPS, and 2% PIP2 (henceforth called 78/20/2 membrane) and x and y dimensions of 100 Å. The second membrane was constructed for the dimer K-Ras4B simulations with the same lipid composition but with x and y dimensions of 140 Å, which is wide enough to accommodate both monomers with and without effectors and prevent periodic box interactions, with the same lipid concentration as the bellow monomer membrane. Both membrane models were equilibrated following the OpenMM[2] CHARMM-GUI protocol described in Ref.[1].

#### Simulation Conditions for the Dimer Simulations on 78/20/2 membrane

#### Equilibration restraints

The equilibration restraints for the systems based on 5VQ2 (henceforth called Crystal systems) are analytically described in Ref. [3]. For the four systems based on 6W4E (henceforth called NMR frame systems), restraints on Cα atoms of K-Ras4B G-domains were applied for the first 50 ns and then were gradually scaled from 1 kcal/mol to 0 for the next 20 ns of the equilibration (out of 100 ns). K-Ras4B G-domain heavy atoms were gradually scaled from 0.1 kcal/mol to 0 over the first 20 ns of the simulation. Lys148 of the CRD domain was restrained close to the membrane with 5 kcal/mol gradually scaled to 0 in the first 50 ns of the equilibration. The CYSF residues were restrained with

0.5 kcal/mol gradually scaled to 0 in the first 5 ns of the equilibration. Crystallographically determined waters were also restrained with a 0.5 Å width 0.1 kcal/mol flat-bottomed potential scaling to 0 over the first 20 ns of the simulation.

**Dimer K-Ras4B simulations**

Four replica simulations were run for the Crystal systems with Raf effectors as described in Ref. [3]. Herein, we extended these simulations for another 1 μs, reaching in the end 2 μs for each replica system. Similarly, for the four replica simulations of the Crystal systems without Raf effectors, we extended the simulations for another 1 μs, reaching in the end 2 μs for each replica system.

Moreover, we simulated for 2 μs the four NMR systems with and without Raf effectors using the ACEMD3 package without restraints, with NPT ensemble, at 310 K temperature using a Monte Carlo x/y isotropic barostat for membranes and a timestep of 4 fs. The cut-off distance for van der Waals interactions and real-space electrostatic interactions was set to 12 Å and the cutoff of the switching function to 10 Å.

All simulations of the dimer K-Ras4B are summarized in Table S1.

**K-Ras4B monomer simulations**

**Solution K-Ras4B monomer simulations**

The 2MSC chain B structure was used. First, it was protonated and residues VAL, ILE, and MET were appended to the end of the chain using the PROPKA algorithm. Two K-Ras4B structures were then built with CHARMM36m; one with GTP bound and one with GDP bound. Both systems were solvated in a water box of 107 $Å^3$. The systems were equilibrated for 20 ns at 300 K temperature with decreasing protein heavy atom and Cα restraints (0.1 kcal/mol for heavy atoms and 1 kcal/mol for Cα atoms) scaling them to 0 kcal/mol with the first 10 ns of the equilibration. Eight production runs were then performed for the GTP-bound K-Ras4B and another eight production runs for the GDP-bound K-Ras4B for 300 ns each. The ACEMD3 package was used to perform the production simulations in the NPT ensemble at 300 K and a timestep of 4 fs. The cut-off distance for van der Waals interactions and real-space electrostatic interactions was set to 9 Å and the cutoff of the switching function to 7.5 Å.

**Membrane-bound K-Ras4B monomer simulations**

It is known that the K-Ras4B monomer can interact with the lipid bilayer through its HVR hexalysine moiety interacting with the polar headgroups and farnesyl moiety interacting with the hydrophobic core of the bilayer.[4] For the construction of the K-Ras4B monomer bound to the membrane without Raf effectors, the structure of the wild-type (WT) monomer K-Ras4B was obtained from chain B of the NMR structure 2MSC.[5] The terminal cysteine 185 was farnesylated with the HTMD[6] CHARMM builder to the CYSF residue, with parameters and topology extracted from the CHARMM toppar_all36_lipid_prot.str file.[7] K-Ras4B monomer was modeled with GTP, inserted into a pre-equilibrated 78/20/2 membrane, and then was solvated, ionized, and built using HTMD with a 0.15 M NaCl concentration.

The monomer K-Ras4B system bound to the membrane was equilibrated with Cα and heavy atom restraints on the whole protein for 90 ns using the ACEMD3 package in the NPT ensemble at 300 K using a Monte Carlo x/y isotropic barostat for membranes and a timestep of 2 fs. After the equilibration, six conformations were selected from the last 20 ns of the simulation, resulting in six different simulations of the monomer K-Ras4B bound to the membrane. The six systems continued to production runs with ACEMD3 for 600 ns each, with the NPT ensemble at 300 K and 2 fs time step. The cut-off distance for van der Waals interactions and real-space electrostatic interactions was set to 9 Å and the cutoff of the switching function to 7.5 Å.

Simulations were analyzed using Markov state models (see below) to investigate the interactions of the K-Ras4B catalytic domain (residues 1 to 169) and HVR (residues 172 to 183) with the membrane lipids. The minimum distance of each head group of the lipids from the HVR tail and the whole K-Ras4B provided insights into the K-Ras4B-lipid interactions (Figure S37 a, b, and c).

Figure S37 shows that PIP2 lipids are moving closer to HVR and interact with the residues of the HVR polybasic loop. Nonetheless, DOPC lipids have a non-specific interaction with the residues of HVR or the K-Ras4B and continuously attach and detach from the protein throughout the simulation time. DOPS lipids have different behavior, forming occasional stable interactions with K-Ras4B.

**Markov State Models**

The monomer K-Ras4B membrane-bound system was analyzed using Markov state models using the HTMD package.[8] Six different production simulations were projected using as metrics the contacts of the Cα atoms of the catalytic domain of the protein (defined as residues 1 to 169) with a) the membrane lipids and b) the HVR tail (defined as the Cα atoms of residues 172 to 183). Contacts were defined as distances smaller than 8 Å as long-range interactions are cut off at 10 Å. This projection allows the Markov model to differentiate between states where the catalytic domain is interacting directly with the membrane or indirectly by interacting with the HVR tail which lies horizontally on top of the membrane. We used TICA[9, 10] with a lag time of 2 ns on the projections to further reduce the dimensionality to 3 dimensions, while maintaining information on the slow processes. Next, the TICA projections were clustered using the k-means algorithm. Various numbers of clusters were tested by evaluating the implied timescales and finally, 800 clusters were selected, as they produced the most converged Markov models. The model was then estimated at a 50 ns lag time and macrostate clustering was performed with the PCCA[11] algorithm to produce two macrostates. Finally, the equilibrium probabilities of the two states were obtained from the model.

From the two macrostates produced by the Markov state models (Figure S37d), macrostate 1 residues of the HVR tail interact very strongly with helix α2 of the catalytic domain and with the PIP2 lipids. In all states, analysis of these systems demonstrated that PIP2 lipids aggregate around the HVR loop. In contrast, macrostate 0 is characterized mostly by nonspecific interactions, with little interaction of helix α4 with PIP2 lipids. This state has a very low population, as proved by its equilibrium probability ($8 \cdot 10^{-4}$ %), compared to macro 1 (99.999% probability).

The same MSM analysis procedure was performed for the eight production simulations of the GTP-bound K-Ras4B in solution and the eight production simulations of the GDP-bound K-Ras4B in solution, with the only difference that the simulations were projected using as metric the contacts of the Cα atoms of the catalytic domain of the protein (defined as residues 1 to 169) with the HVR tail (defined as the Cα atoms of residues 172 to 183).

**Binding Site identification with SiteMap and FTMap**

Binding site prediction on monomer K-Ras4B structures was performed using SiteMap[12] and FTMap[13] tools, with all default parameters unless mentioned otherwise. The structure for the binding site identification on the K-Ras4B monomer was isolated from the most conserved macrostates of the Markov state models. For each protein structure, a list of up to five potential binding sites was identified and analyzed.

**SiteMap**

SiteMap performs binding site identification on protein structures in three stages. First, it locates the sites and defines them by a set of "site points" on a grid. Second, it generates the "maps" that define the sites and uses them for visualization. Finally, it uses the site-point groups produced in the site-finding stage and the grids produced in the mapping stage, to evaluate the sites in terms of several properties. Some of these properties are, a) the number of site points, a measure of the size of the site or the number of grid points required to define the predicted site, b) Exposure and Enclosure properties measuring how open the site is to the solvent: a tight binding site has low exposure rate (average value = 0.49) and high enclosure rate (average value = 0.78), c) Hydrophobic and Hydrophilic character, properties measuring the relative hydrophobic/hydrophilic character of the cavity (both properties are calibrated to have average values of 1 for tight binding sites), and d) SiteScore and Dscore, defined by the same descriptors, but different coefficients:

$$SiteScore = 0.0733\, sqrt(n) + 0.6688\, e - 0.20\, p$$

$$DScore = 0.094\, sqrt(n) + 0.60\, e - 0.324\, p$$

where, n = the number of site points (capped at 100), e = enclosure, and p = hydrophilic score (capped at 1.0 to limit the impact of hydrophilicity in charged and highly polar sites).

Scores greater than 1 suggest a site of particular promise. Additionally, a SiteScore of 0.80 has been found to accurately distinguish between drug-binding and non-drug-binding sites.

**FTMap**

FTMap server available at https://ftmap.bu.edu/ identifies binding hot spots on proteins, i.e., areas of the protein with a major contribution to ligand-binding free energy. The algorithm uses 16 small organic molecules as probes of varying size, shape, and polarity and finds their favorable positions using empirical free energy functions. Specifically, it identifies the most favorable positions for each probe, clusters the probes, and ranks the clusters based on average free energy. Protein areas that bind different probe clusters (consensus sites) are identified as binding hot spots.

### Supporting Figures

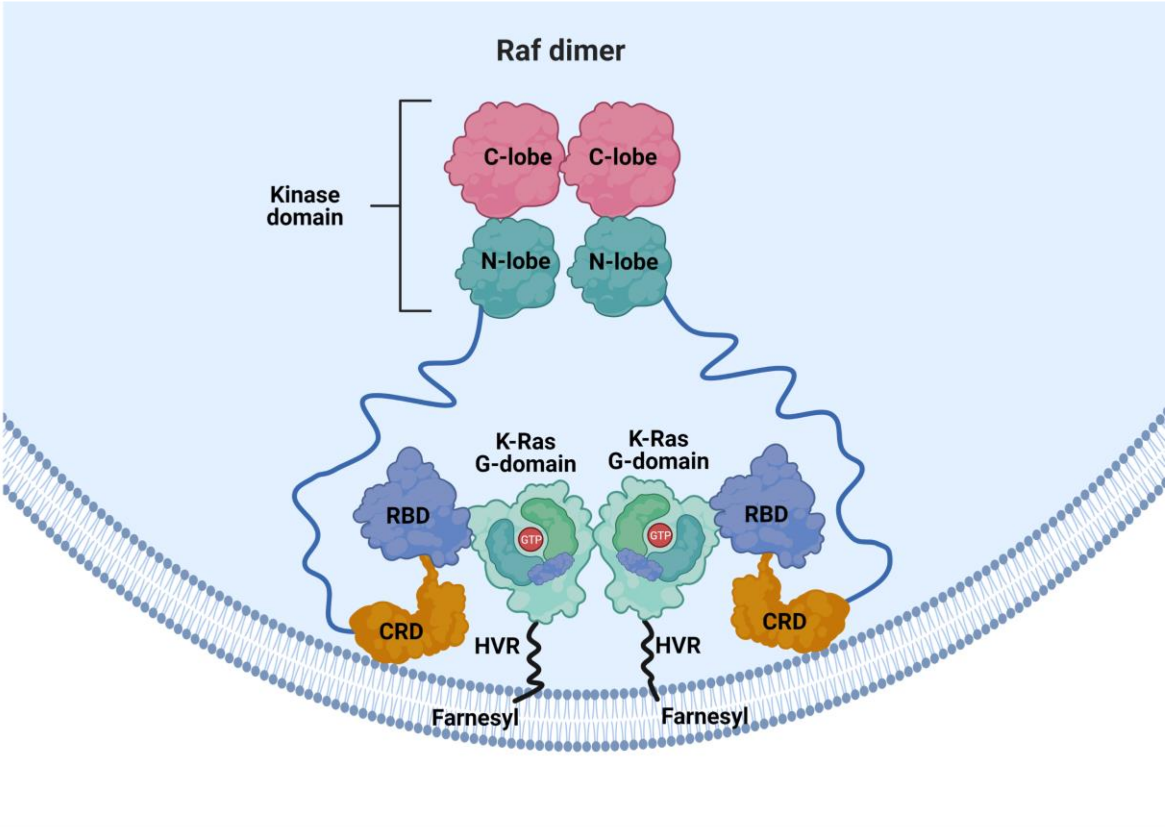


**Figure S1.** A schematic representation of the Ras dimerization in the membrane, which facilitates the dimerization of Raf, which is essential for Raf activation in cells. The RBD domain of Raf is bound to Ras with high affinity and connected to the CRD domain through a disordered linker. The CRD is attached to the inner leaflet of the cell membrane and connected to the kinase domain of Raf through a long flexible linker.

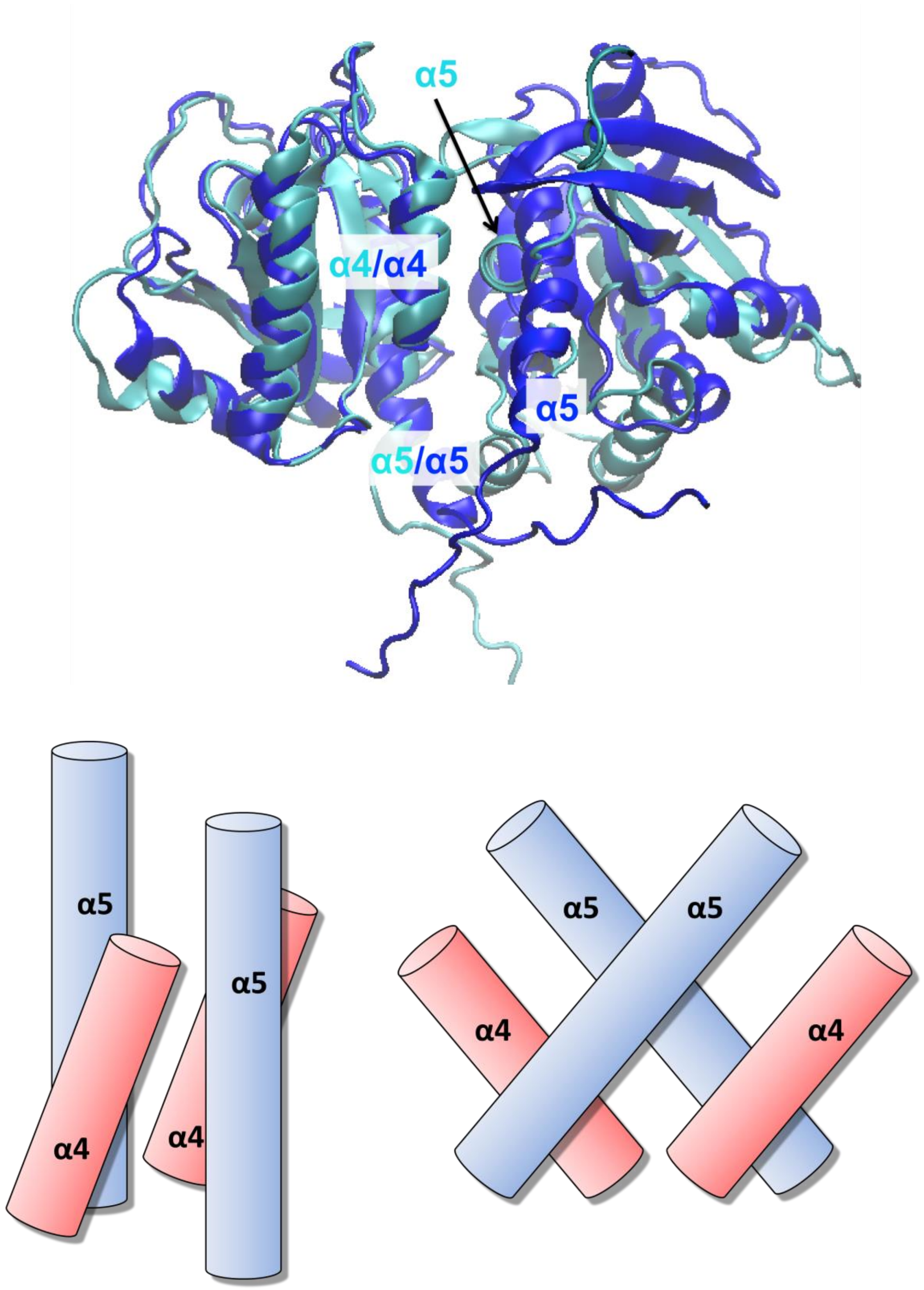


**Figure S2.** Upper panel: The model structure of Ref.[14] (PDB ID: 6W4E colored in blue) overlapped with the structure Ref.[15] (PDB ID: 5VQ2 colored in cyan). Lower panel: Helices α4 and α5 of opposite monomers (colored in red and blue) are almost parallel to each other (forming a 20° angle) in Ref.,[14] while in Ref.[15] helix α5 is rotated 90° with respect to the α4 of the opposite monomer.

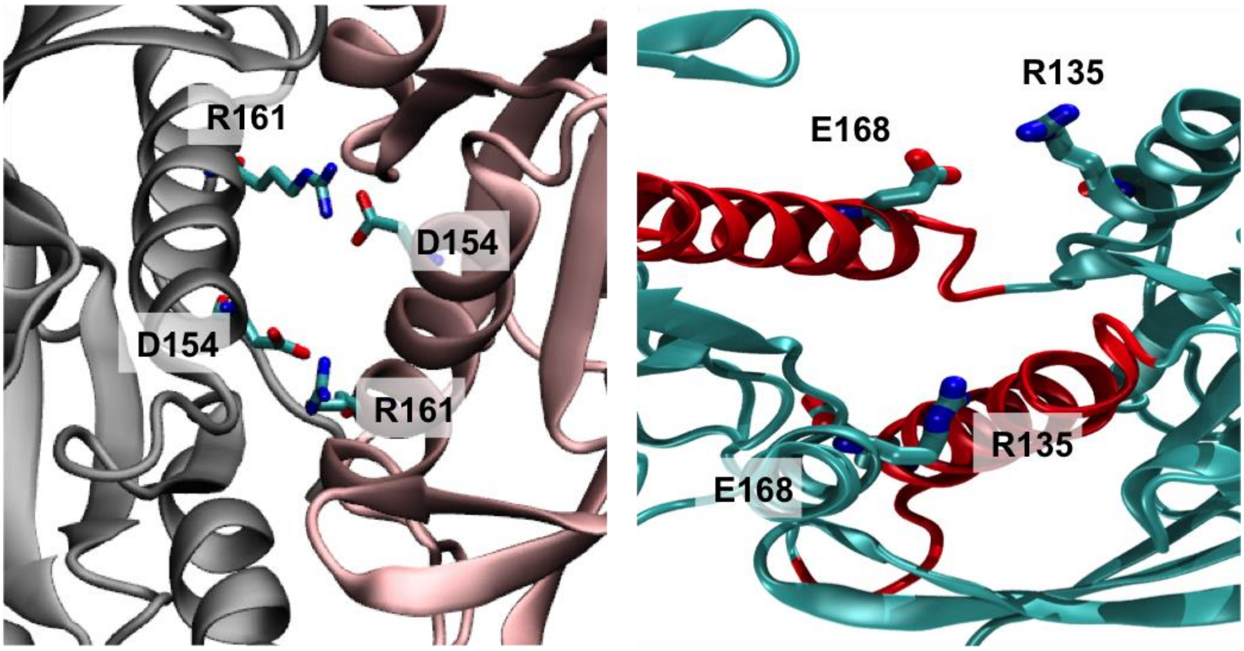


**Figure S3.** The α5-α5 interface of the K-Ras4B based on the PDB ID 5VQ2 structure is mediated by R161-D154 intermolecular salt bridges (left), while the α4-α5 interface of the K-Ras4B based on the PDB ID 6W4E structure is mediated by the R135-E168 salt bridge (right).

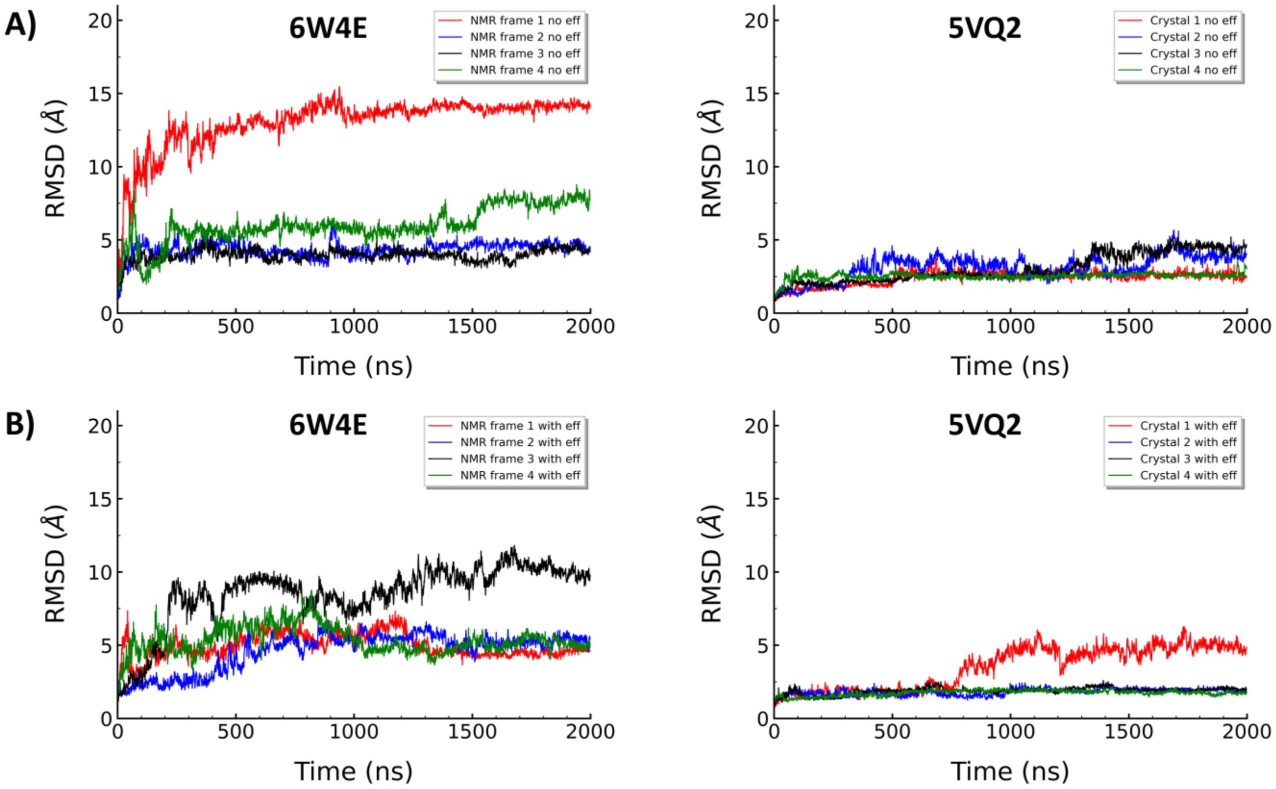


**Figure S4.** A) RMSD of the four replicas of K-Ras4B dimer simulations without effectors based on the 6W4E (upper left) and 5VQ2 structure (upper right) and B) RMSD of the four replicas of K-Ras4B dimer simulations with effectors based on the 6W4E (bottom left) and 5VQ2 structure (bottom right).

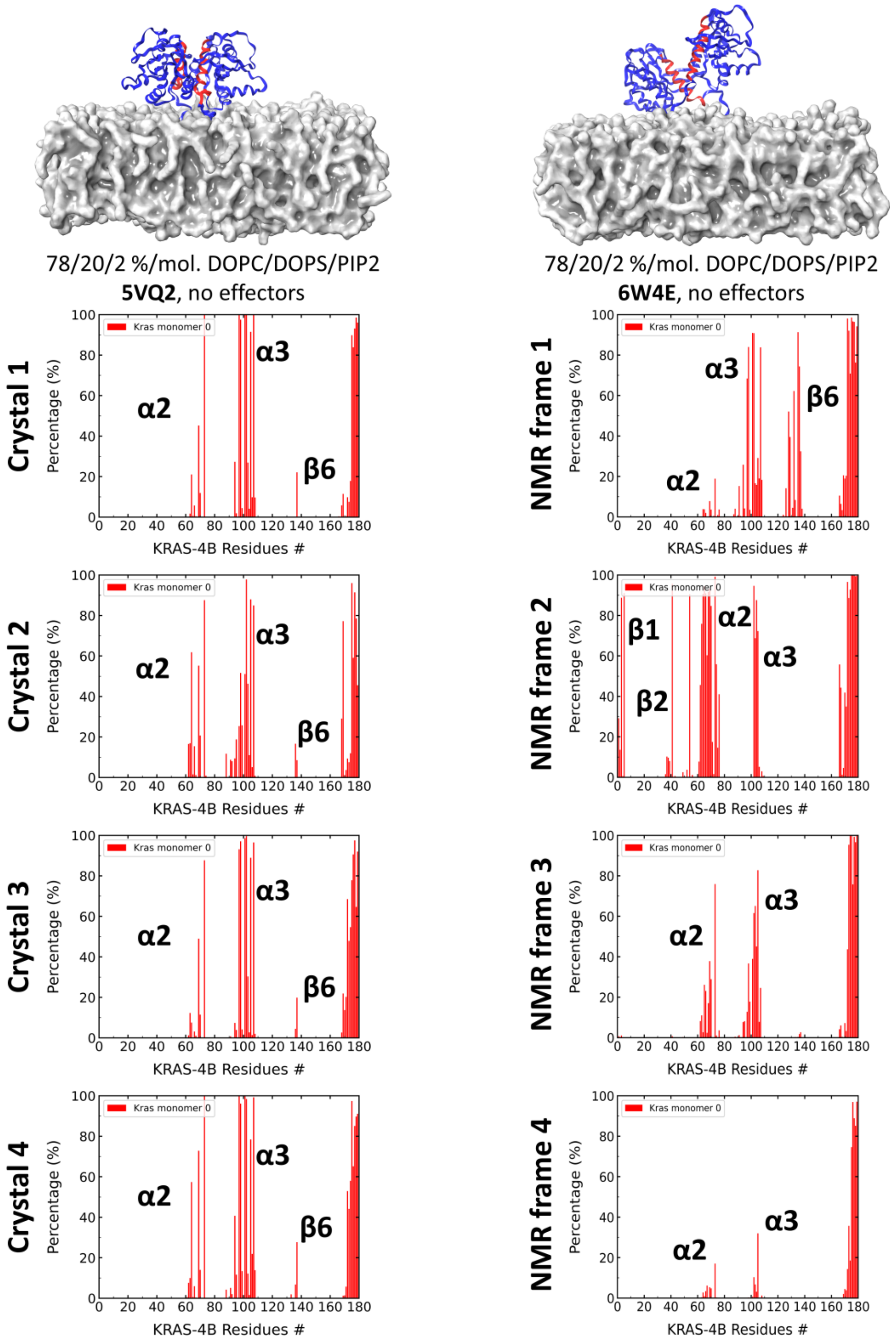


**Figure S5.** Percentage of interactions between the G-domain of K-Ras4B and the membrane for the 5VQ2 and 6W4E structures without effectors. Interactions of the first K-Ras4B (monomer 0) are depicted here.

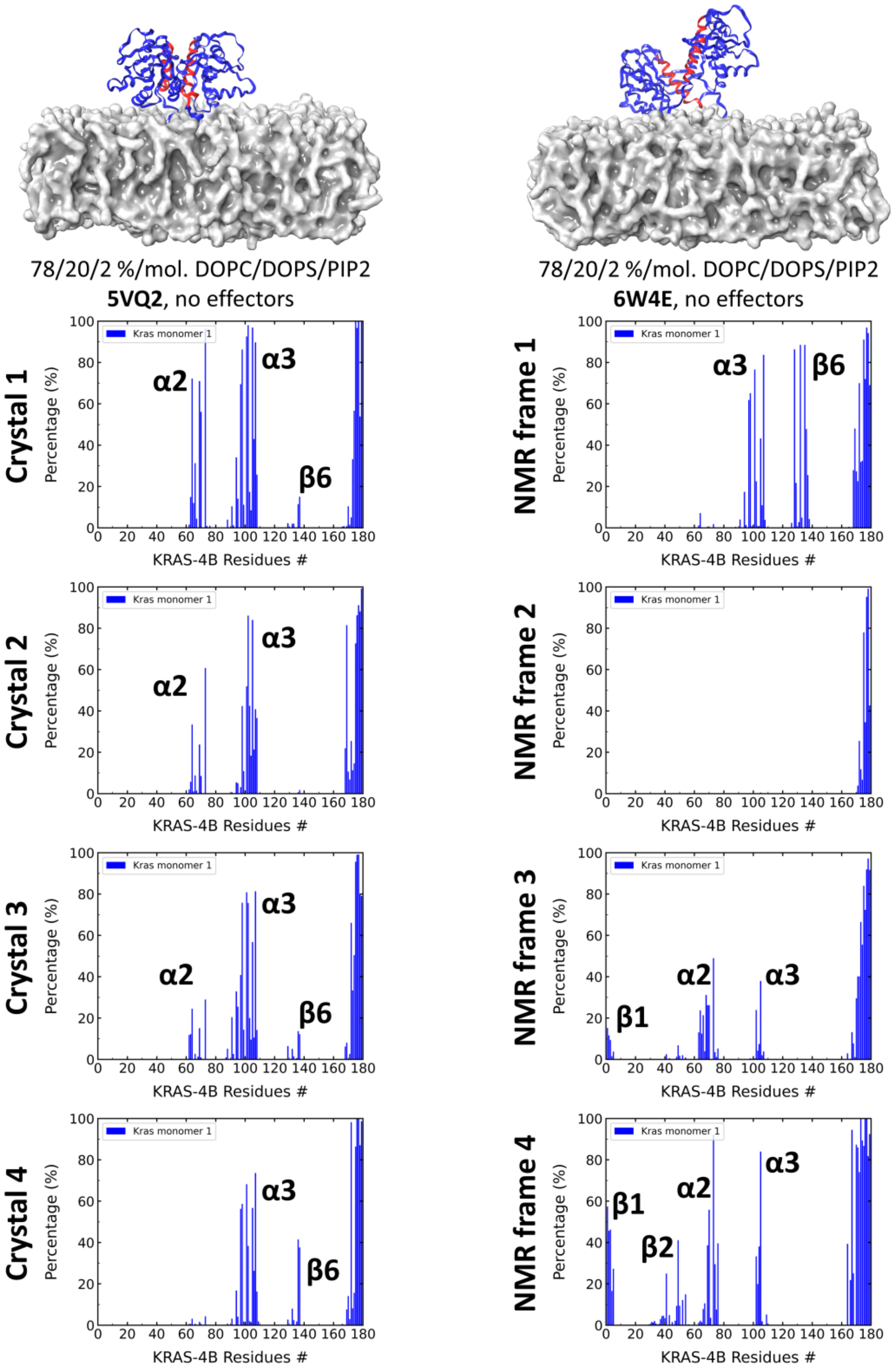


**Figure S6.** Percentage of interactions between the G-domain of K-Ras4B and the membrane for the 5VQ2 and 6W4E structures without effectors. Interactions of the first K-Ras4B (monomer 1) are depicted here.

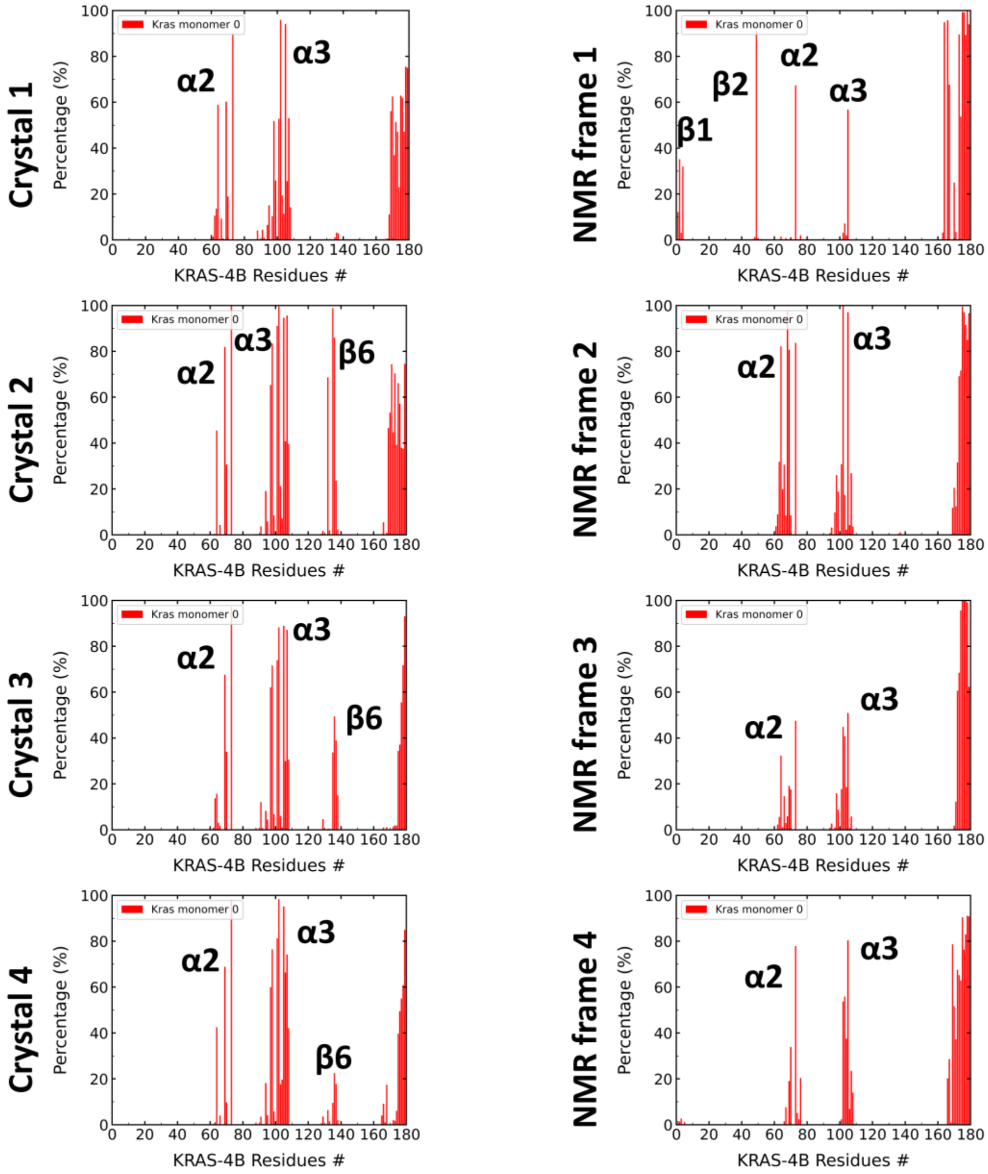


**Figure S7.** Percentage of interactions between the G-domain of K-Ras4B and the membrane for the 5VQ2 and 6W4E structures with effectors. Interactions of the first K-Ras4B (monomer 0) are depicted here.

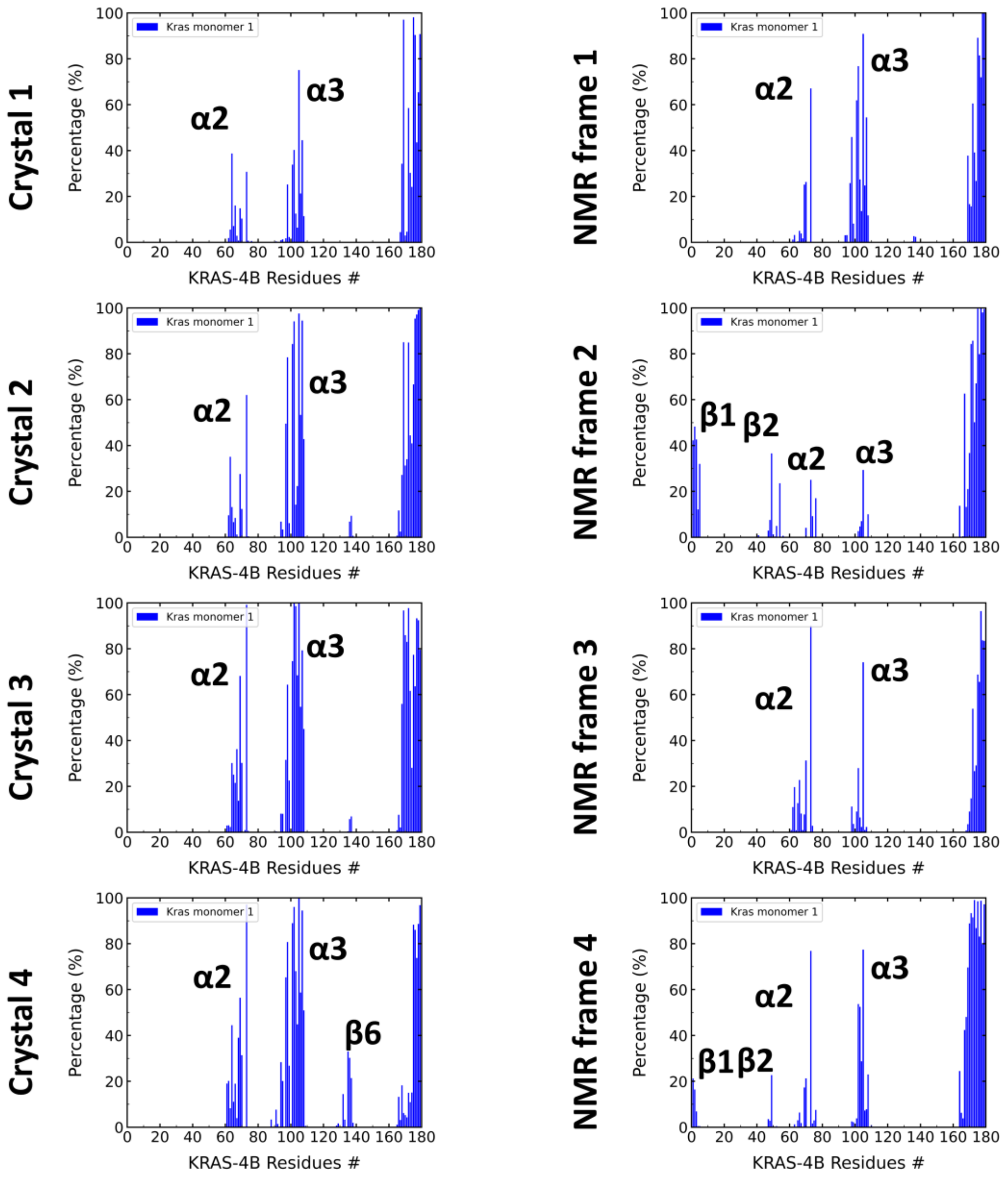


**Figure S8.** Percentage of interactions between the G-domain of K-Ras4B and the membrane for the 5VQ2 and 6W4E structures without effectors. Interactions of the first K-Ras4B (monomer 1) are depicted here.

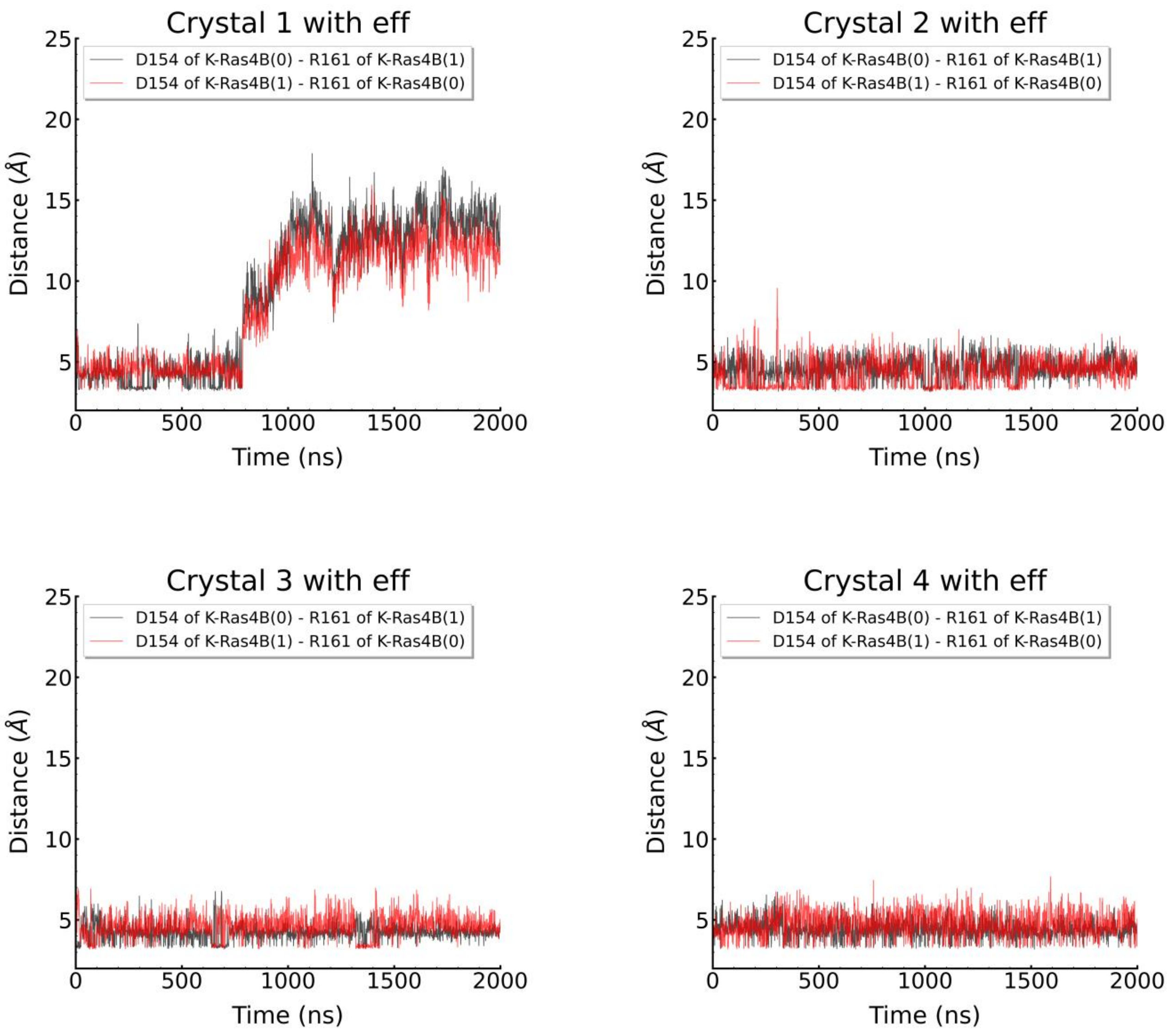


**Figure S9.** Distance of salt bridges between D154 and R161 of K-Ras4B for the structure with PDB ID 5VQ2 with effectors (Crystal #1-4).

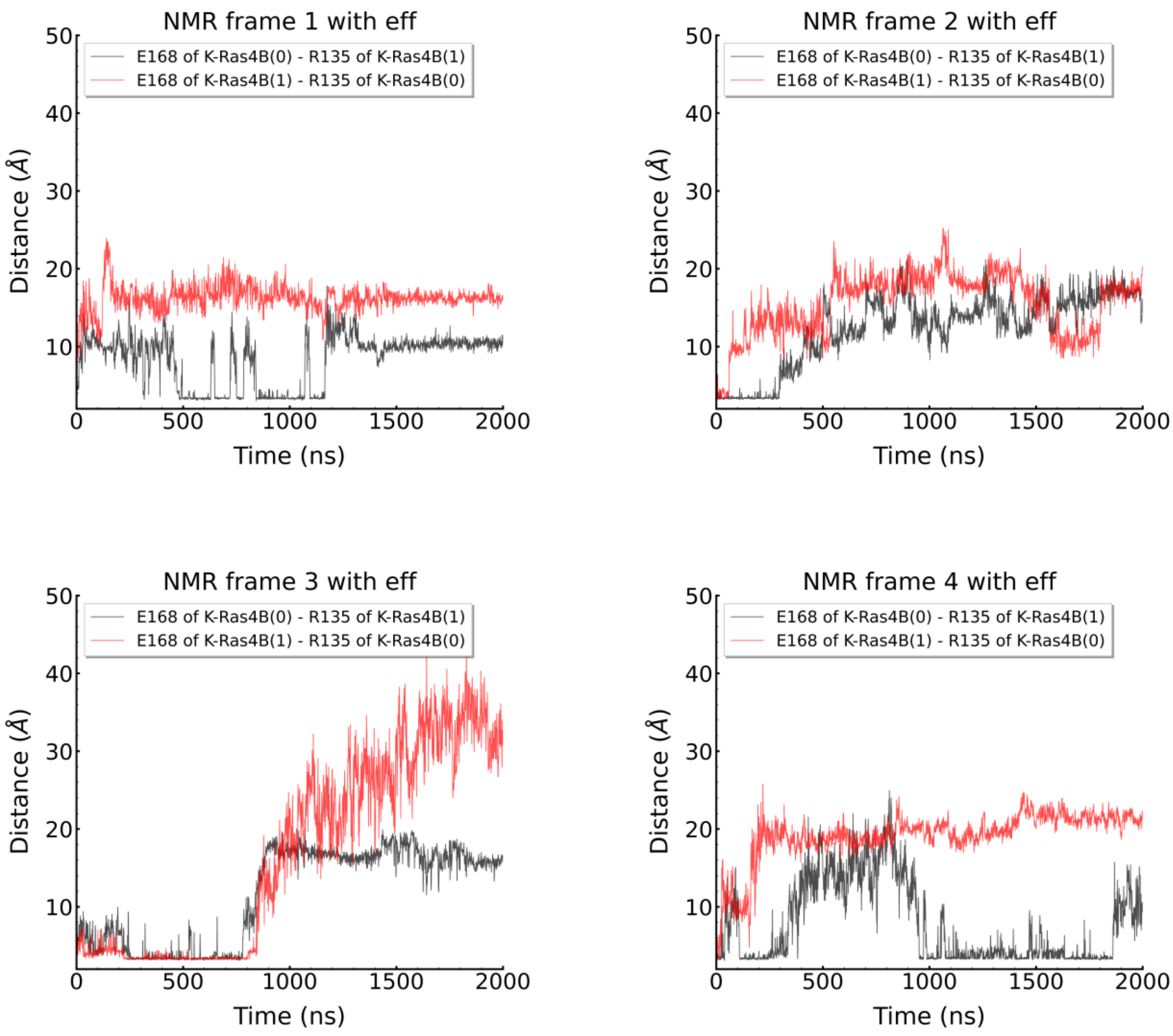


**Figure S10.** Distance of salt bridges between R135 and E168 of K-Ras4B for the structure with PDB ID 6W4E with effectors (NMR frames #1-4).

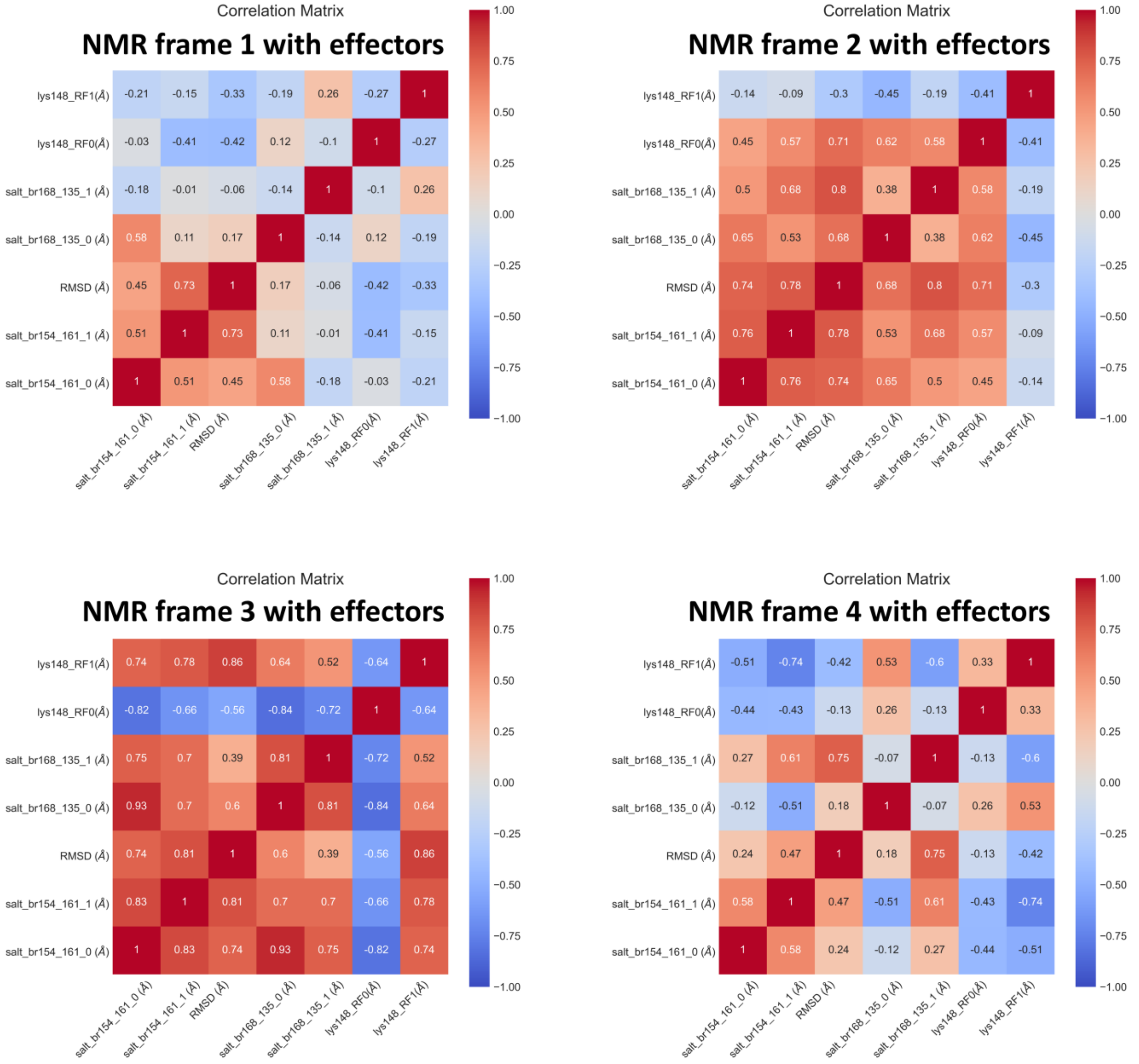


**Figure S11.** Visualization of the correlation matrices between RMSD, crucial salt bridges' distances, and CRD's distances from the center of the membrane for the structures with effectors based on PDB ID 6W4E. The calculation was performed in both K-Ras4B monomers as well as Raf monomers.

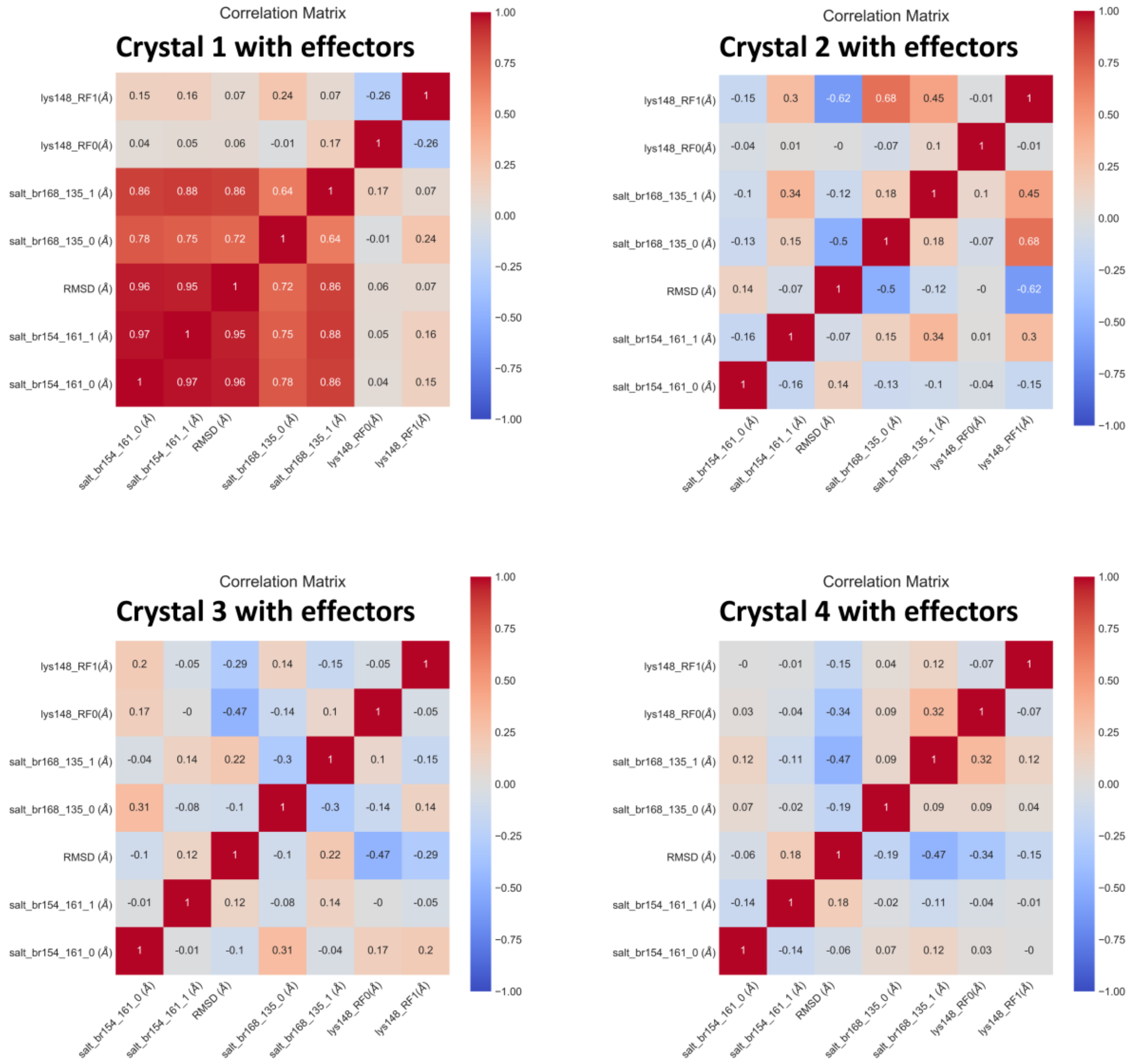


**Figure S12.** Visualization of the correlation matrices between RMSD, crucial salt bridge's distances, and CRD's distances from the center of the membrane for the structures with effectors based on PDB ID 5VQ2. The calculation was performed in both K-Ras4B monomers as well as Raf monomers.

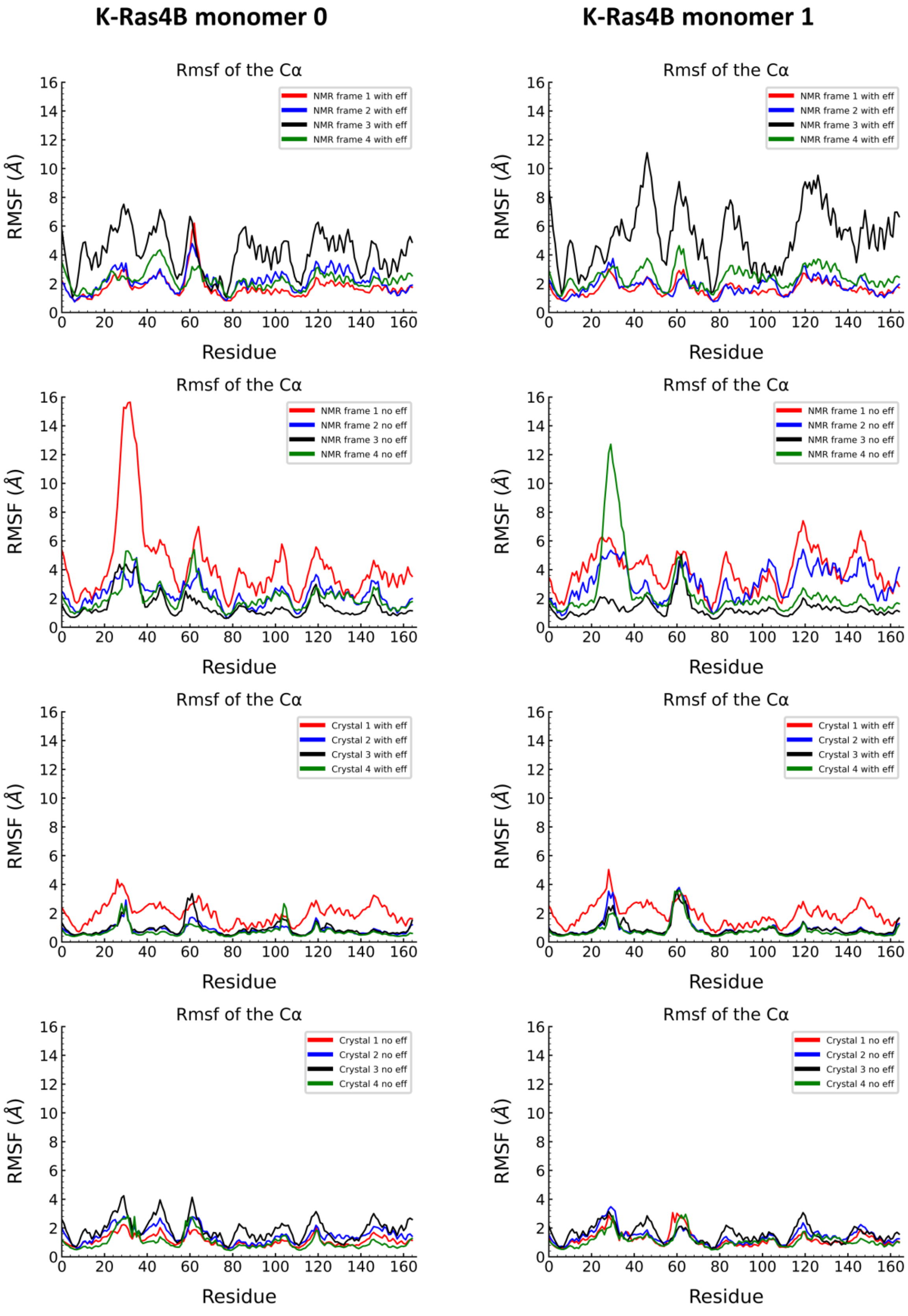


**Figure S13.** RMSF of each amino acid of the K-Ras4B monomer 0 and K-Ras4B monomer 1 with and without the effectors for the structures based on PDB ID: 5VQ2 and 6W4E.

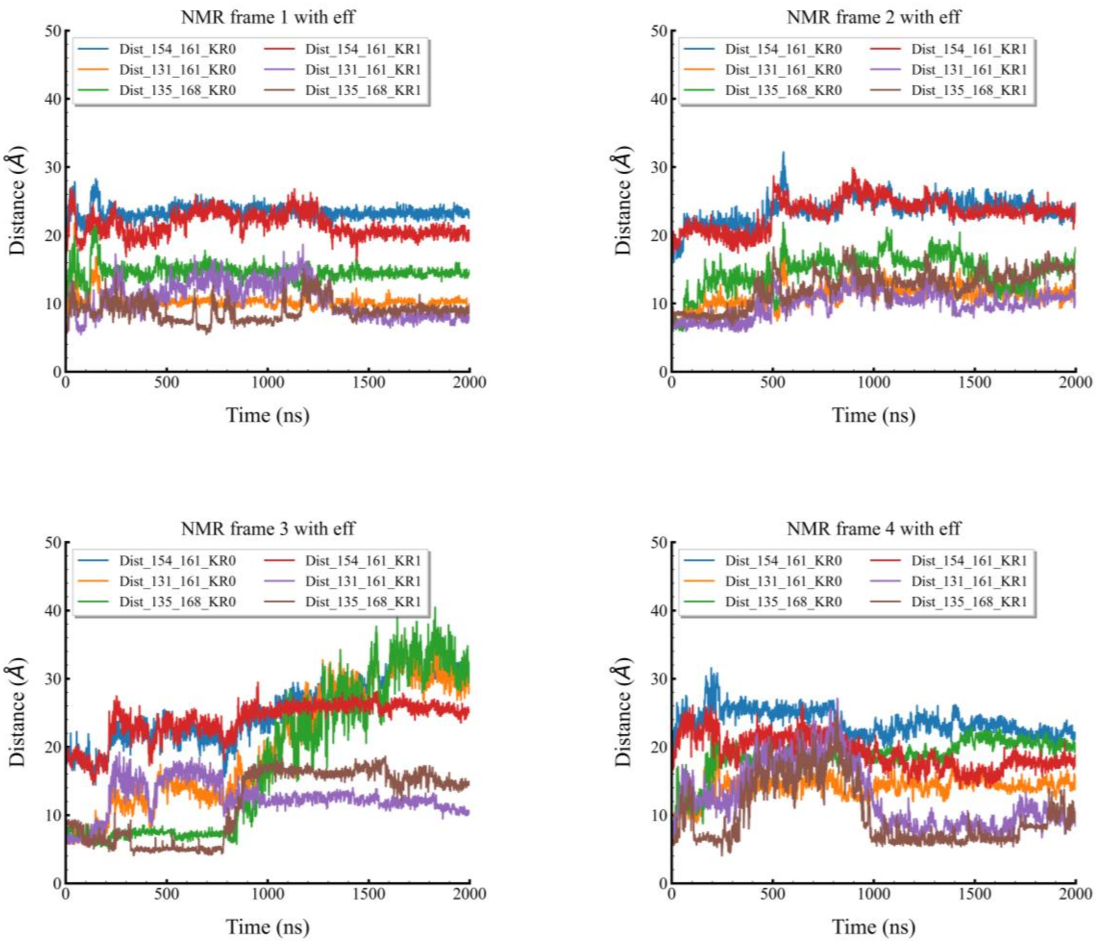


**Figure S14.** Distance of COM of D154-R161, Q131-R161, and R135-E168 amino acids for 6W4E systems with effectors (NMR frame with effectors #1-4).

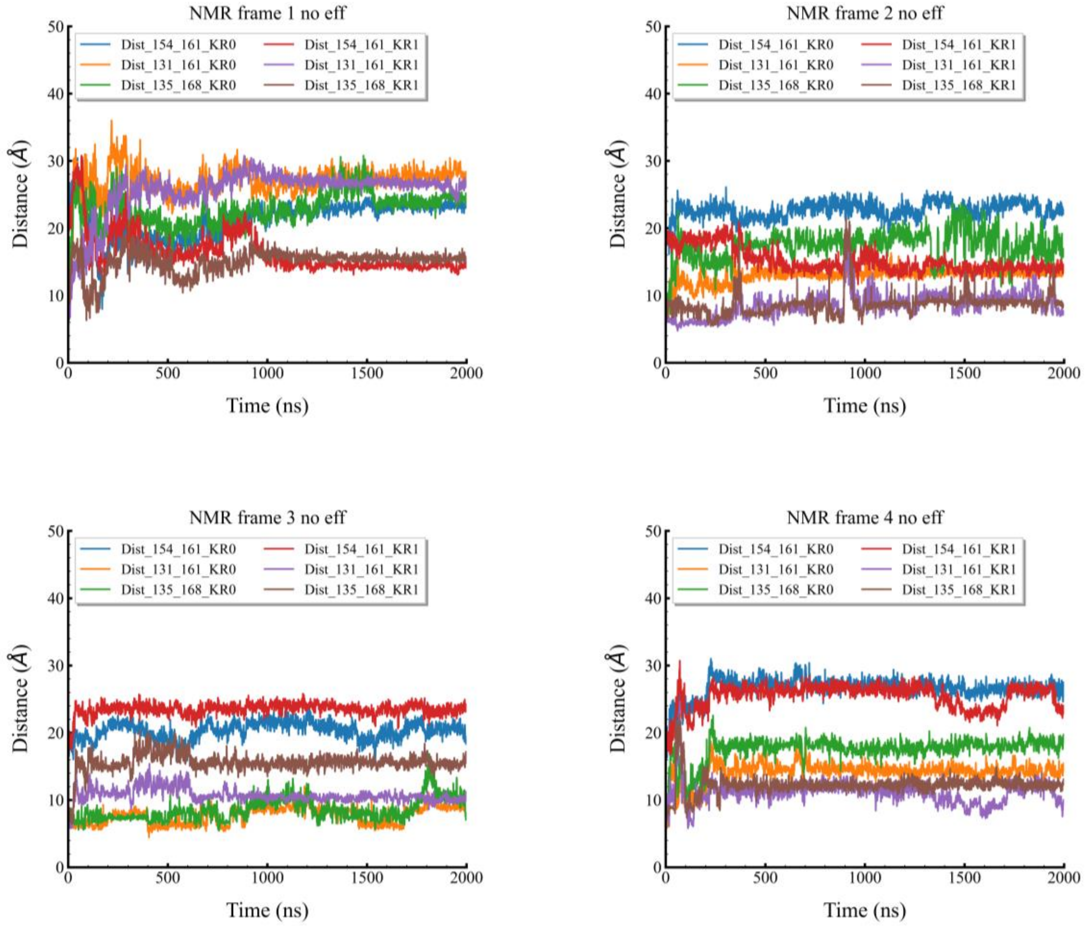


**Figure S15.** Distance of COM of 154-161, 131-161, and 135-168 amino acids for 6W4E systems without effectors (NMR frame without effectors #1-4).

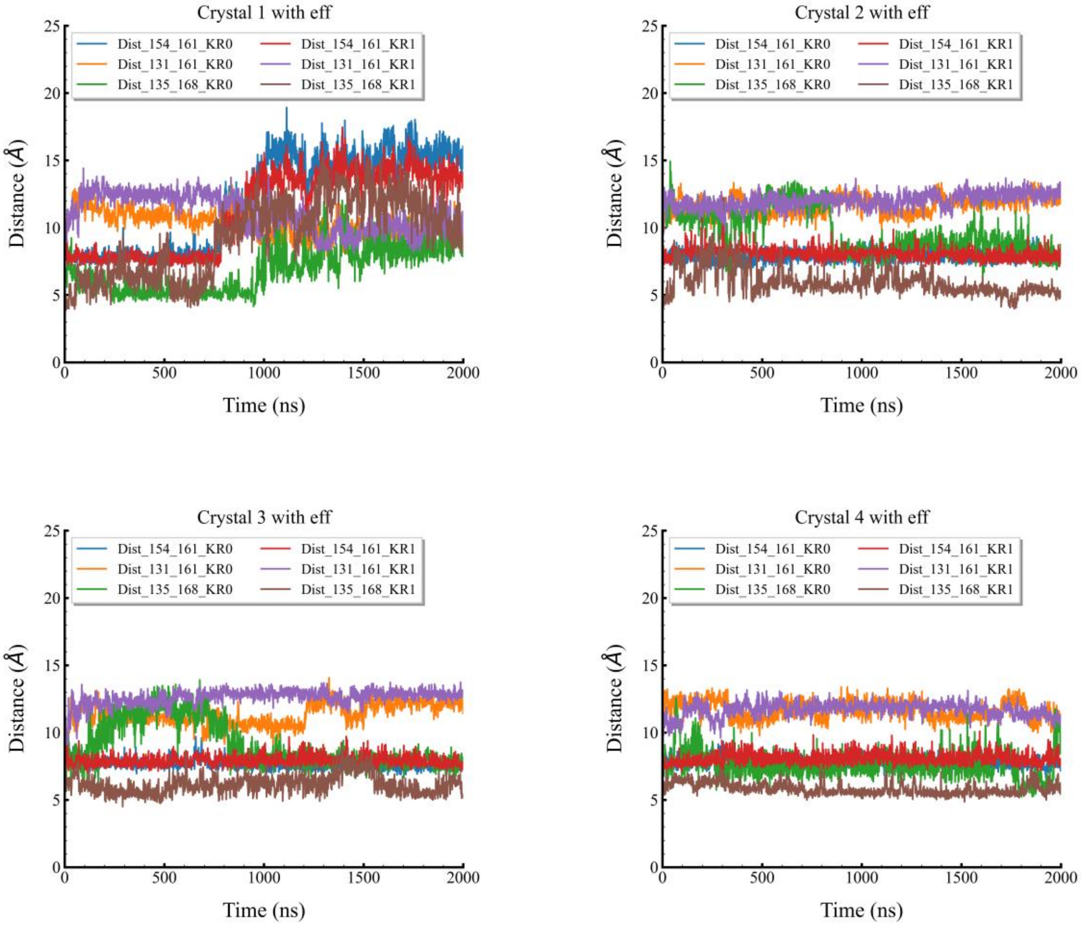


**Figure S16.** Distance of COM of 154-161, 131-161, and 135-168 amino acids for 5VQ2 systems with effectors (Crystal with effectors #1-4).

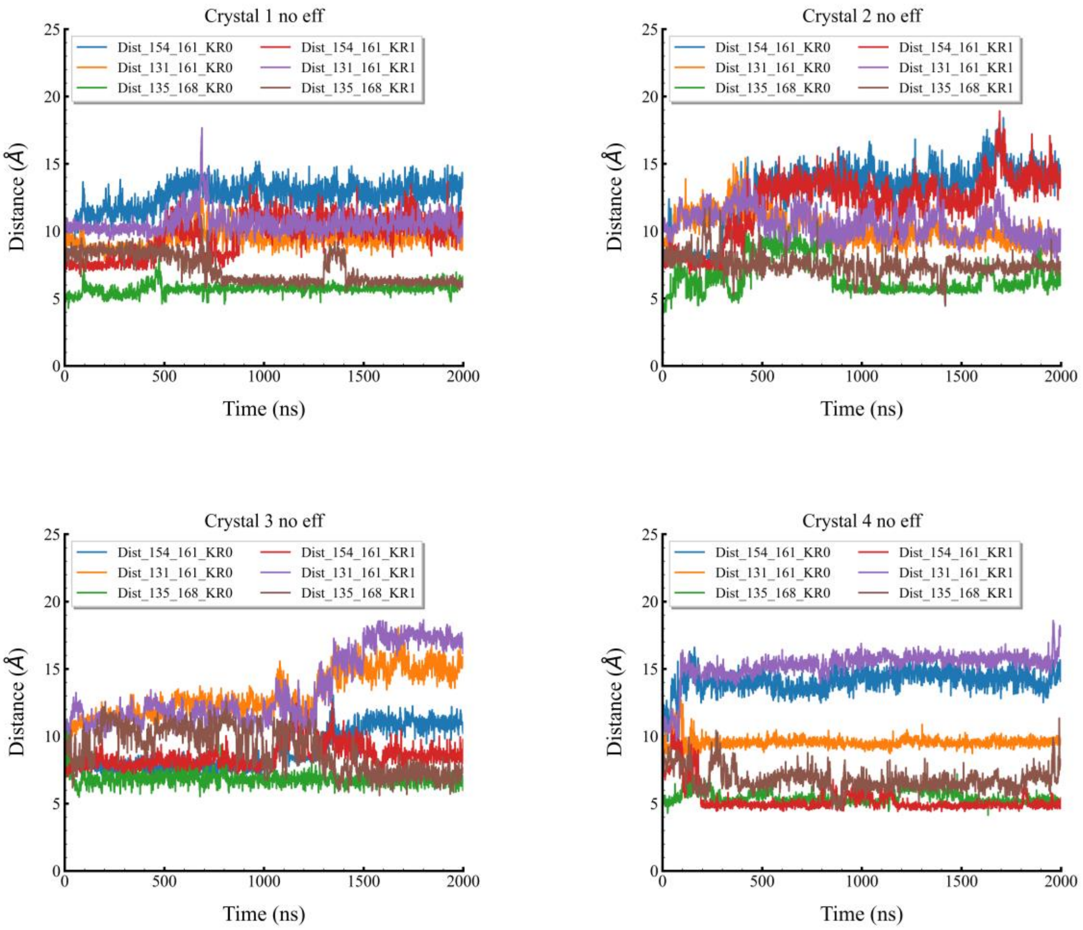


**Figure S17.** Distance of COM of 154-161, 131-161, and 135-168 amino acids for 5VQ2 systems without effectors (Crystal without effectors #1-4).

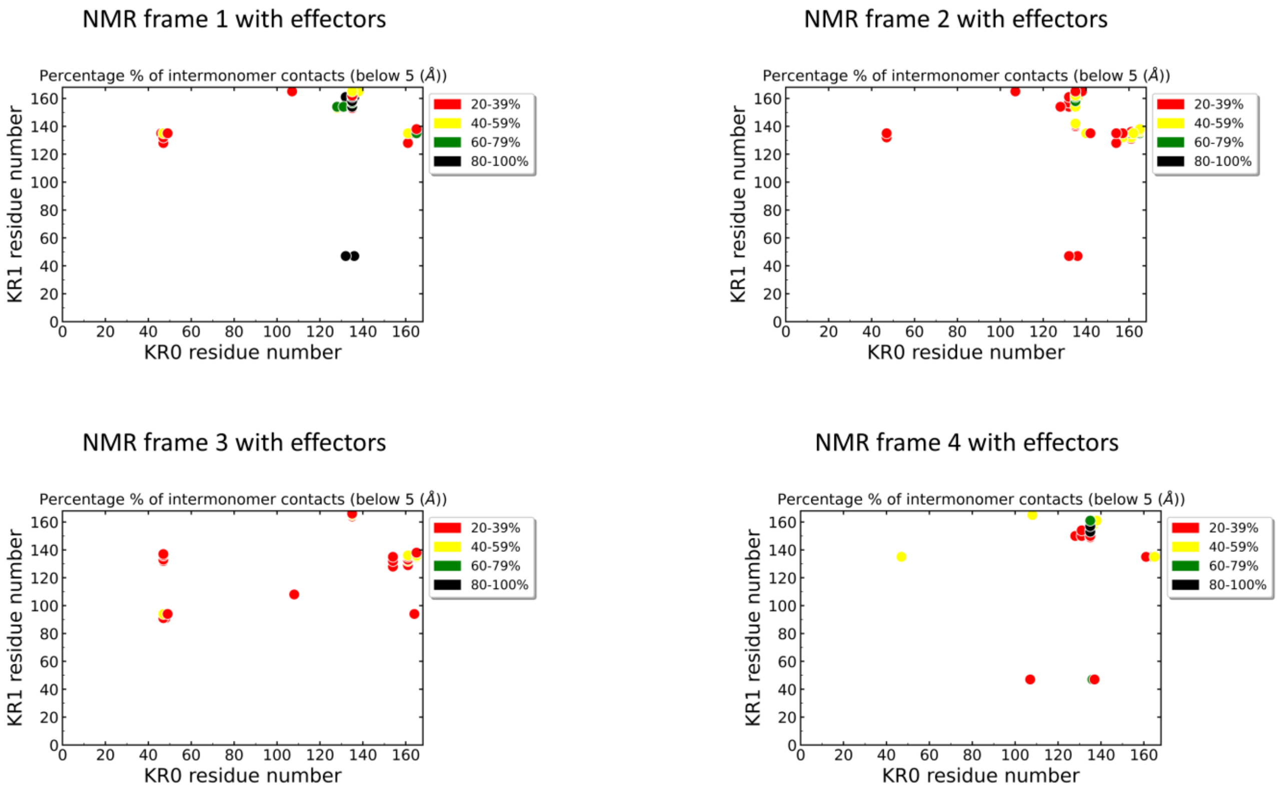


**Figure S18.** Frequency of the inter-monomer interactions between the amino acids of the two monomers K-Ras4B monomer 0 (KR0) and K-Ras4B monomer 1 (KR1) for the structure based on PDB ID: 6W4E with effectors.

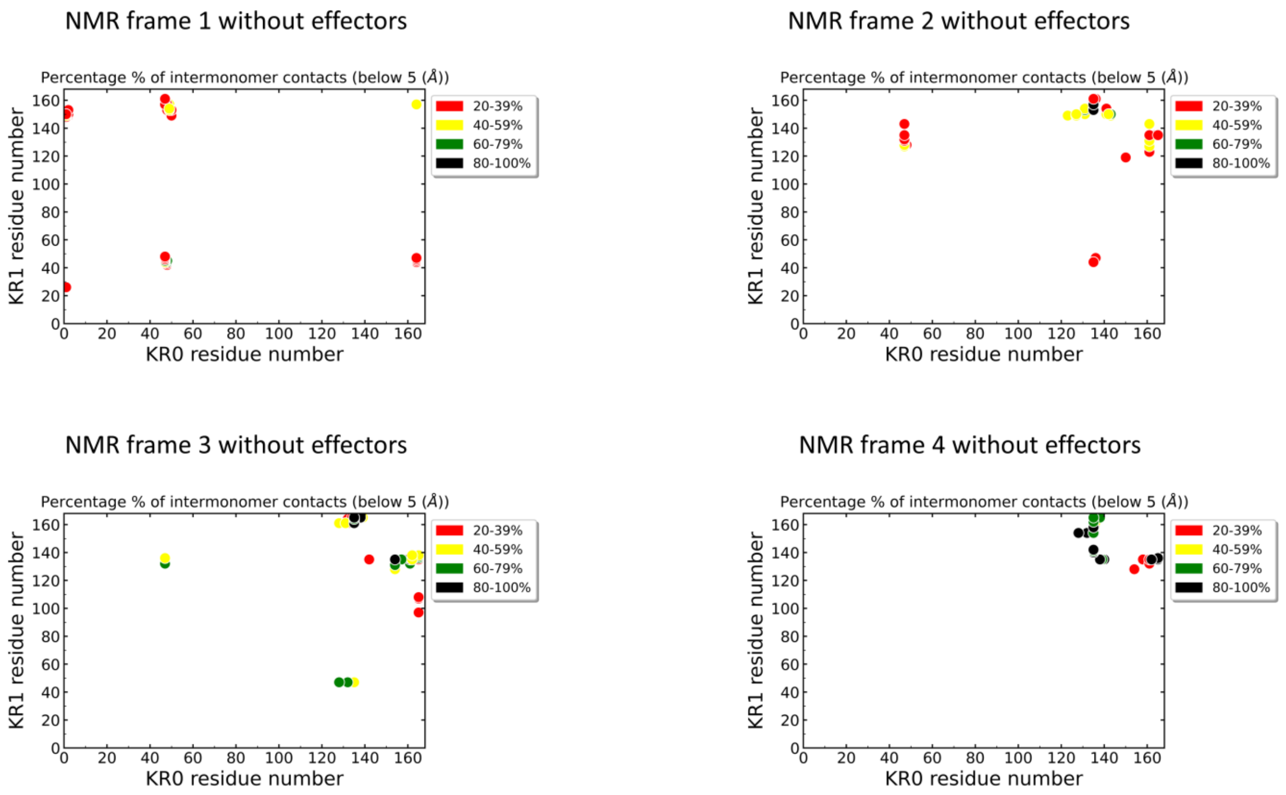


**Figure S19.** Frequency of the inter-monomer interactions between the amino acids of the two monomers K-Ras4B monomer 0 (KR0) and K-Ras4B monomer 1 (KR1) for the structure based on PDB ID: 6W4E without effectors.

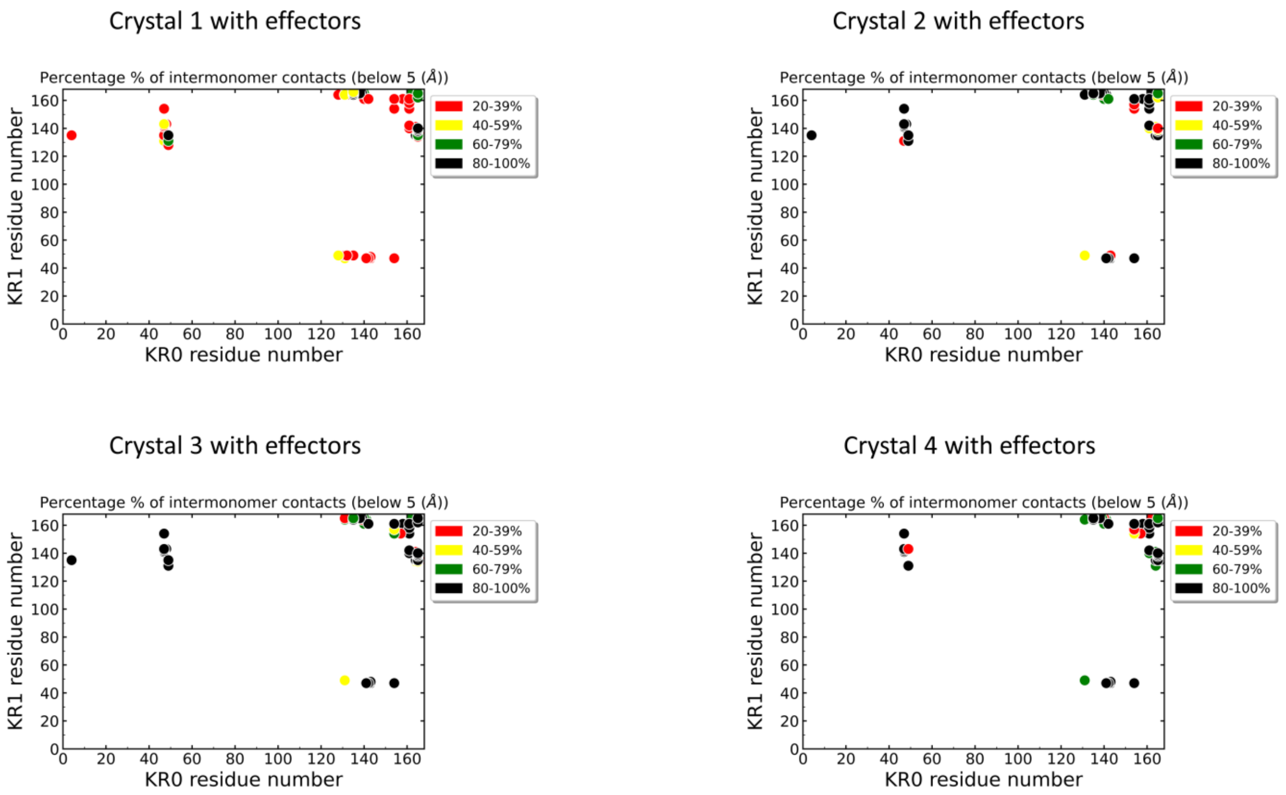


**Figure S20.** Frequency of the inter-monomer interactions between the amino acids of the two monomers K-Ras4B monomer 0 (KR0) and K-Ras4B monomer 1 (KR1) for the structure based on PDB ID: 5VQ2 with effectors.

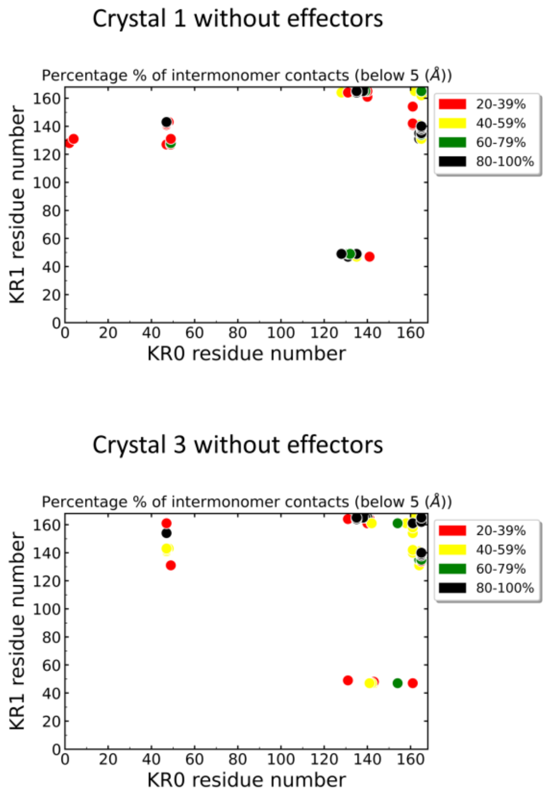


**Figure S21.** Frequency of the inter-monomer interactions between the amino acids of the two monomers K-Ras4B monomer 0 (KR0) and K-Ras4B monomer 1 (KR1) for the structure based on PDB ID: 5VQ2 without effectors.

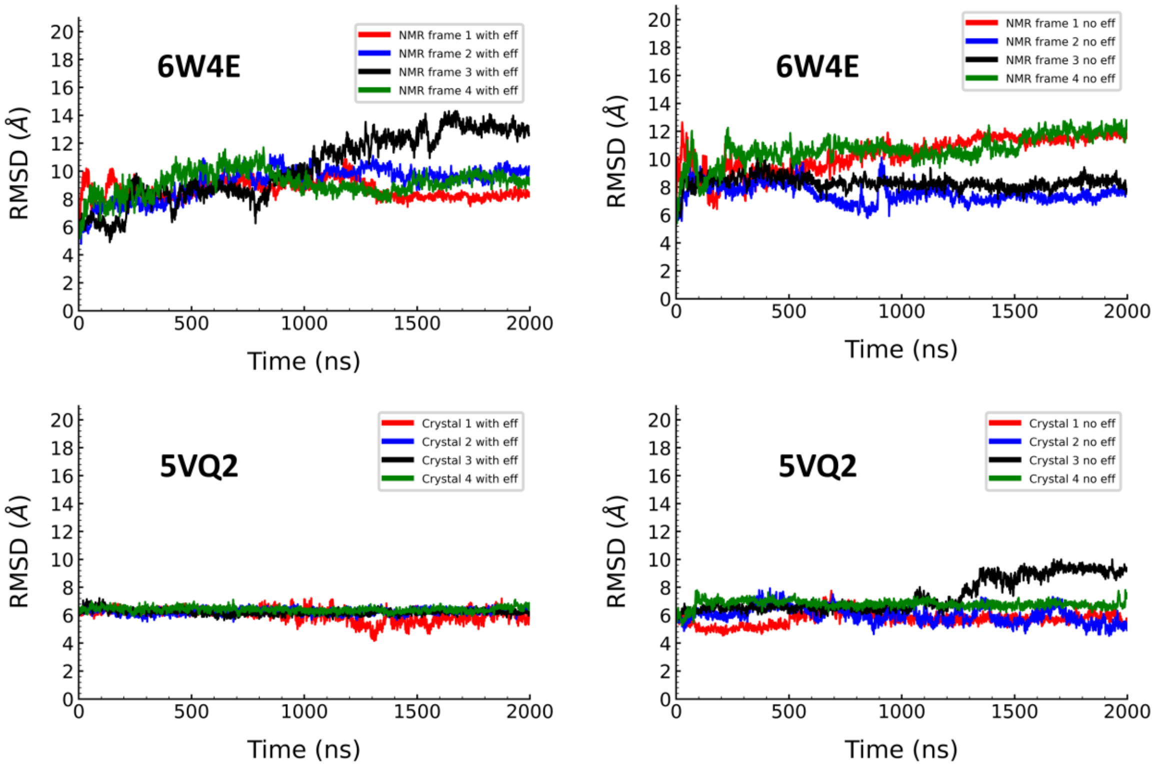


**Figure S22.** RMSD time series comparison for the 5VQ2 structure (bottom panel) versus the 6W4E (upper panel) simulations. 5VQ2 simulations were plotted, taking as reference NMR frame 1 with effectors and 6W4E simulations were plotted, taking as reference Crystal 1 simulations with effectors.

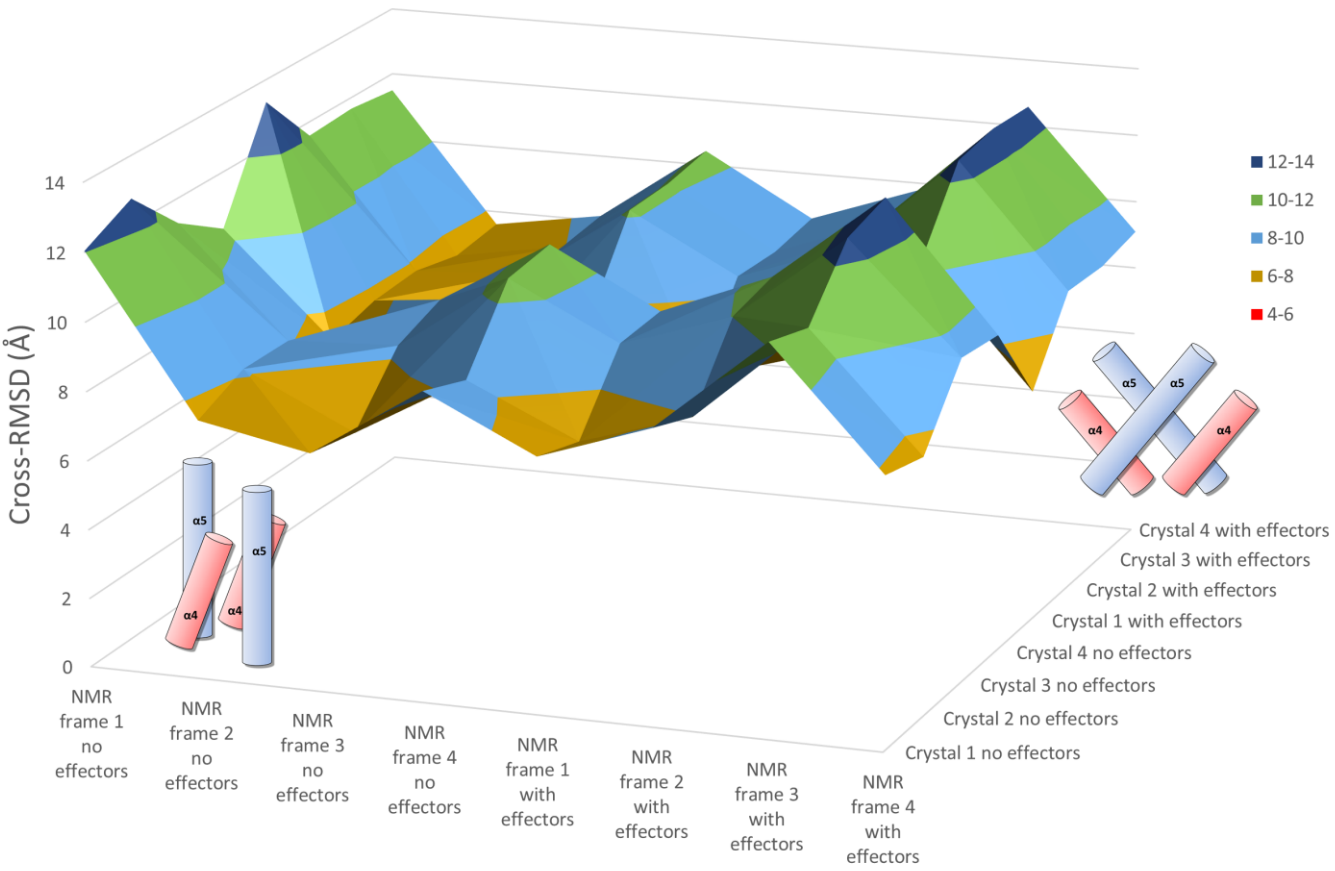


**Figure S23.** Cross-RMSD calculations of the 5VQ2 and 6W4E cluster representatives.

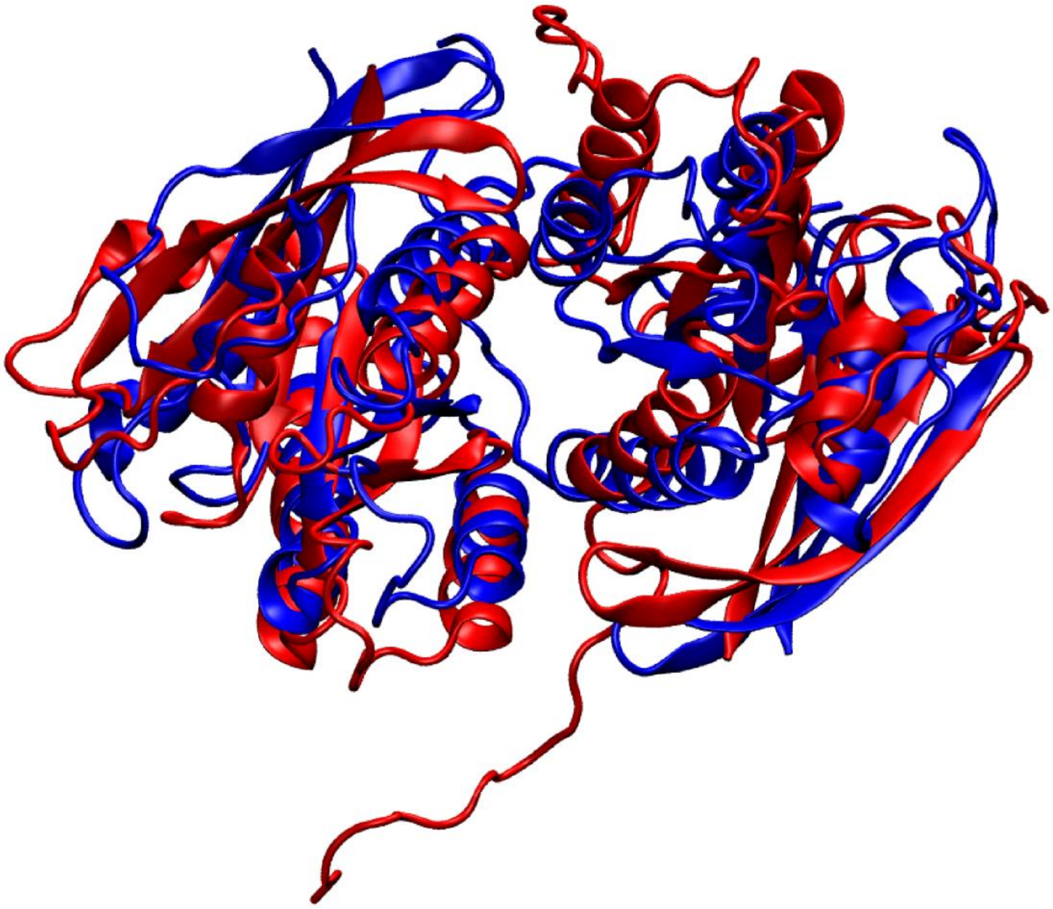

**Figure S24.** Alignment of the cluster representatives NMR frame 3 no effectors (blue) and Crystal 1 with effectors (red) (Raf effectors are omitted for clarity).

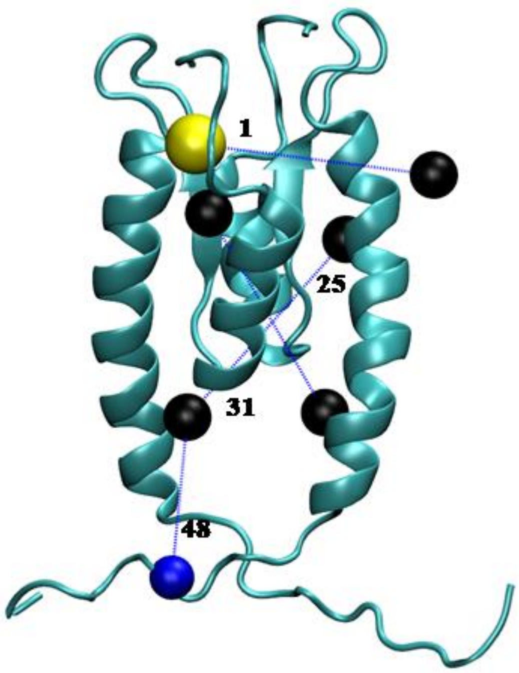


**Figure S25.** Numbering of the PRE examined distances. For clarity, only four distance numbers are depicted (on the α5-α4 interface).

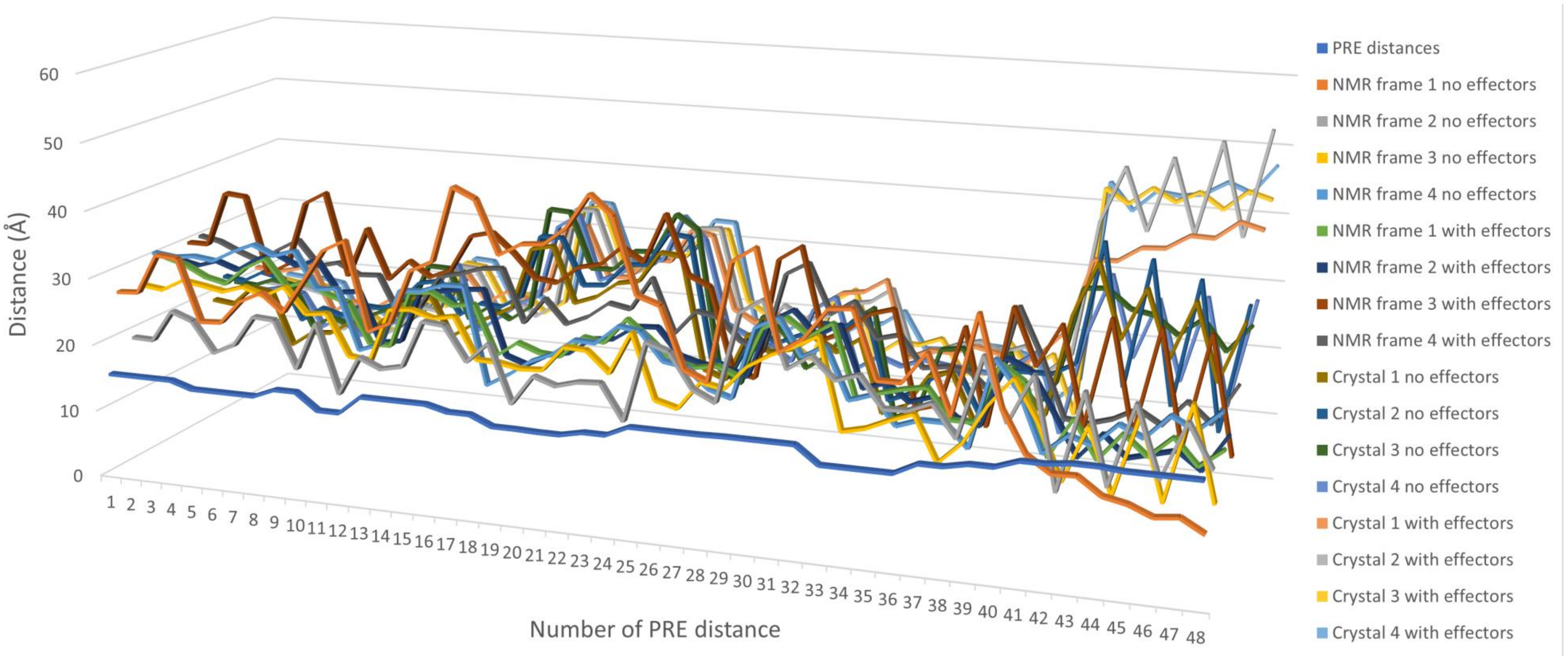


**Figure S26.** Average of the 48 calculated PRE distances for all simulations in comparison to the experimental values.

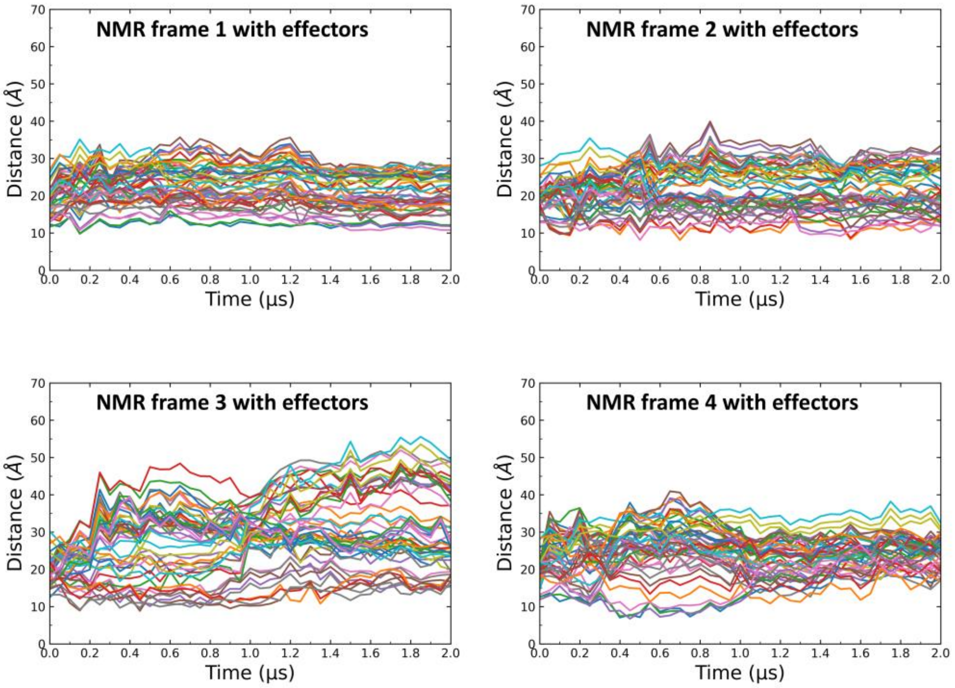


**Figure S27.** Time series of PRE distances obtained from the literature as calculated from the simulations for the 6W4E systems with effectors (NMR frame with effectors #1-4).

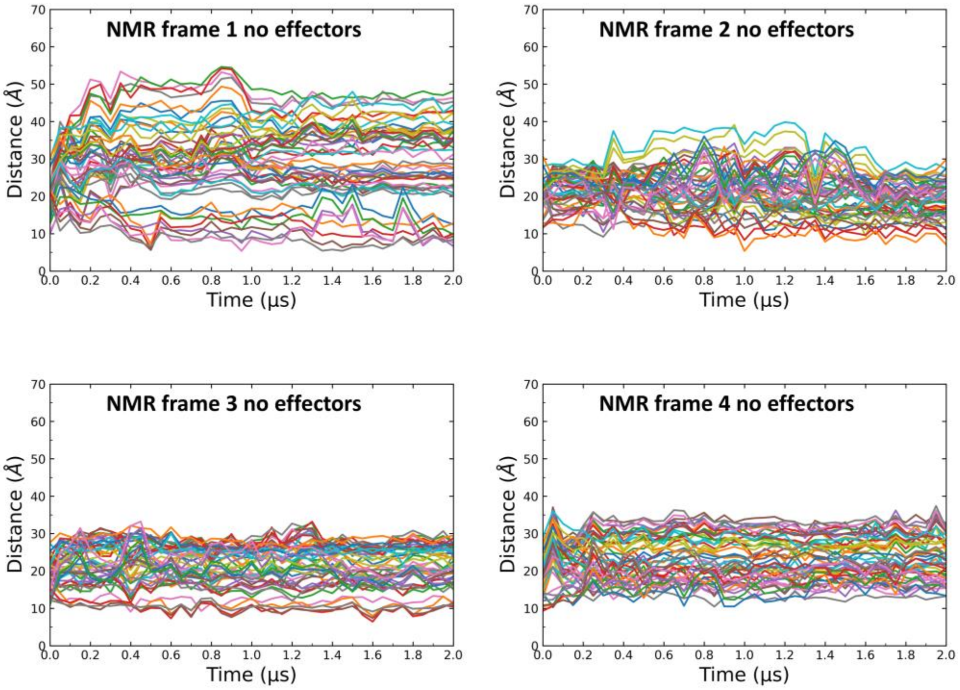


**Figure S28.** Time series of PRE distances obtained from literature as calculated from the simulations for the 6W4E systems without effectors (NMR frame without effectors #1-4).

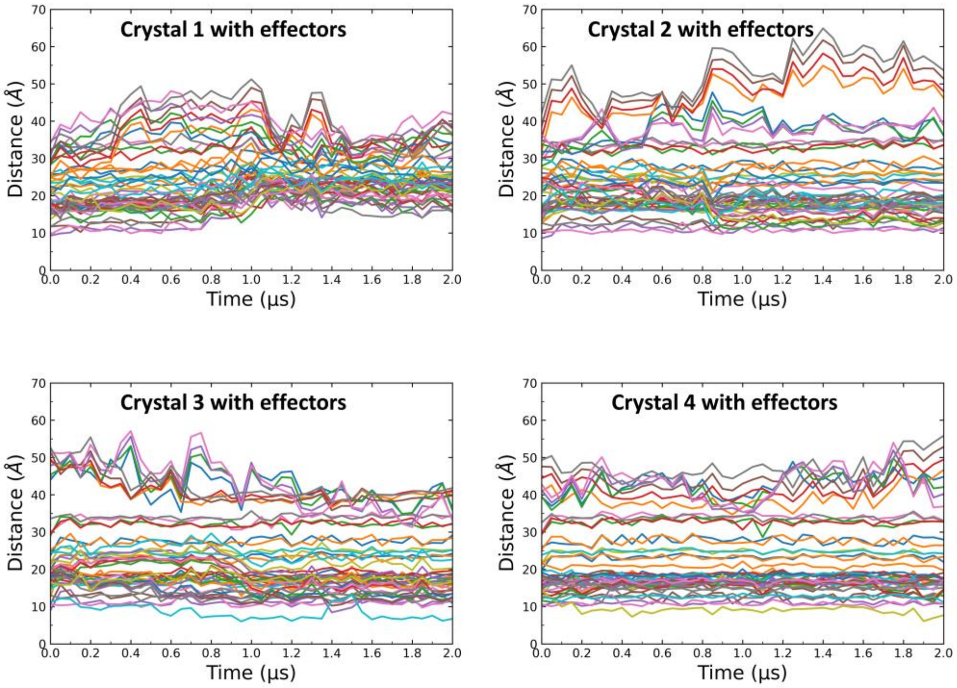


**Figure S29.** Time series of PRE distances obtained from literature as calculated from the simulations for the 5VQ2 systems with effectors (Crystal with effectors #1-4).

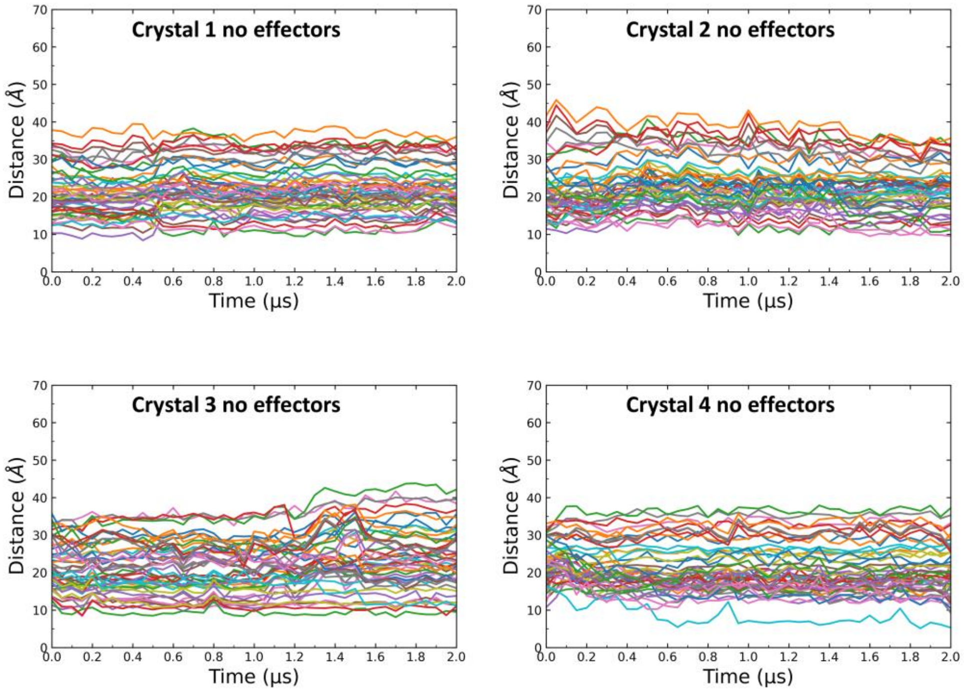


**Figure S30.** Time series of PRE distances obtained from literature as calculated from the simulations for the 5VQ2 systems without effectors (Crystal without effectors #1-4).

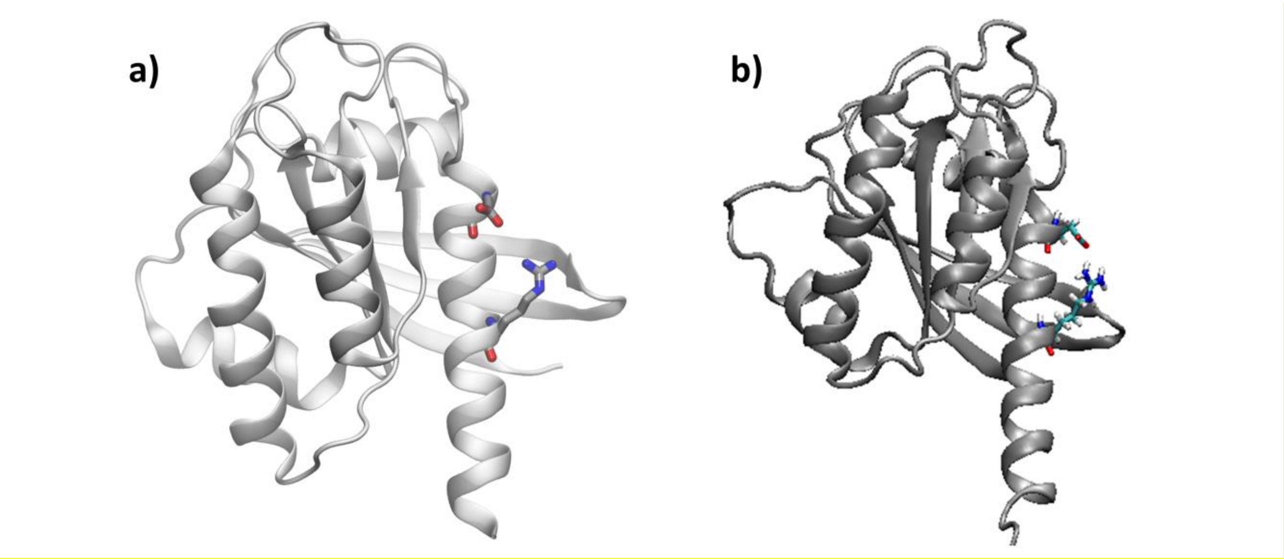


**Figure S31. a)** Monomer K-Ras4B crystal structure (PDB: 6GOF), **b)** Monomer K-Ras4B from our simulations. It is observed that the Asp154 and Arg161 of the K-Ras4B α5 helix interact through the formation of an intramolecular salt bridge.

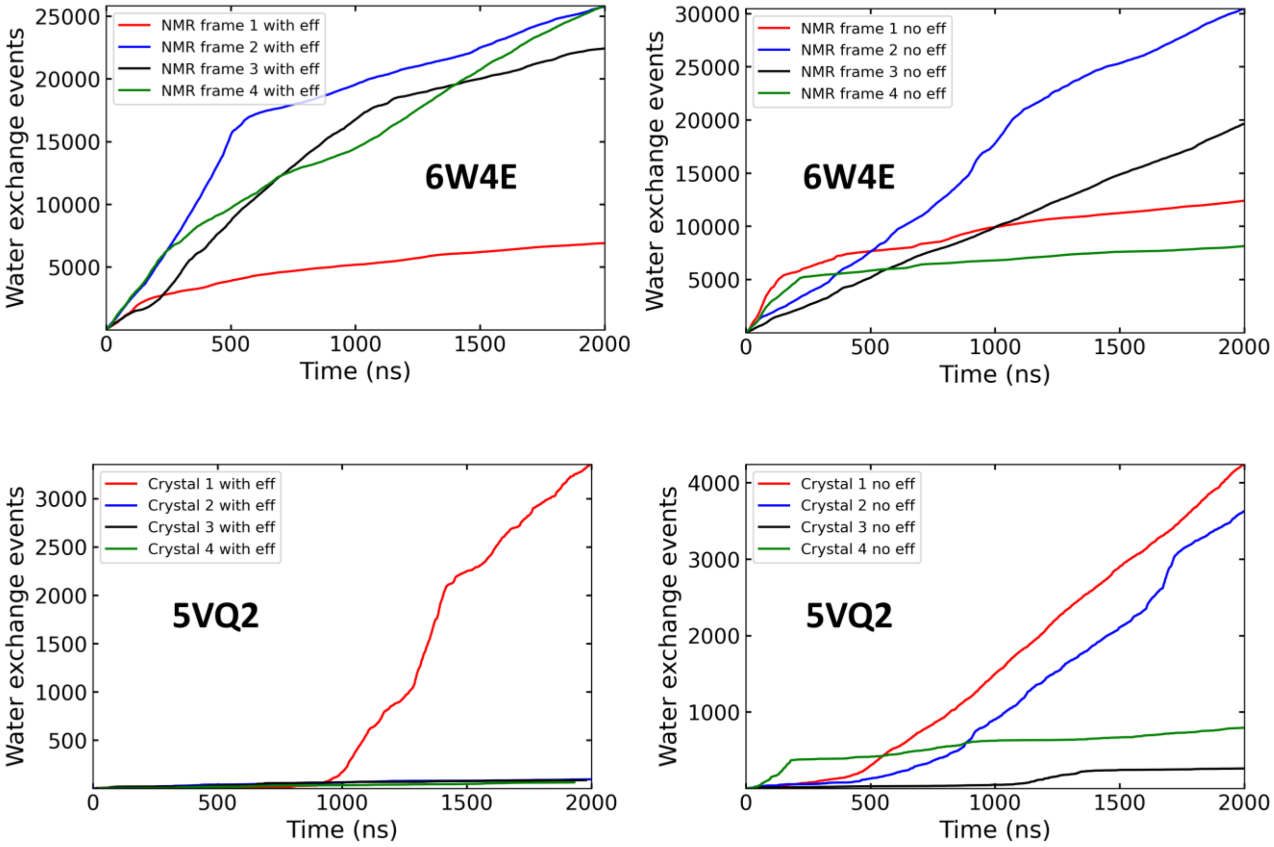


**Figure S32.** Water exchange events for the systems based on 6W4E and 5VQ2 with and without effectors.

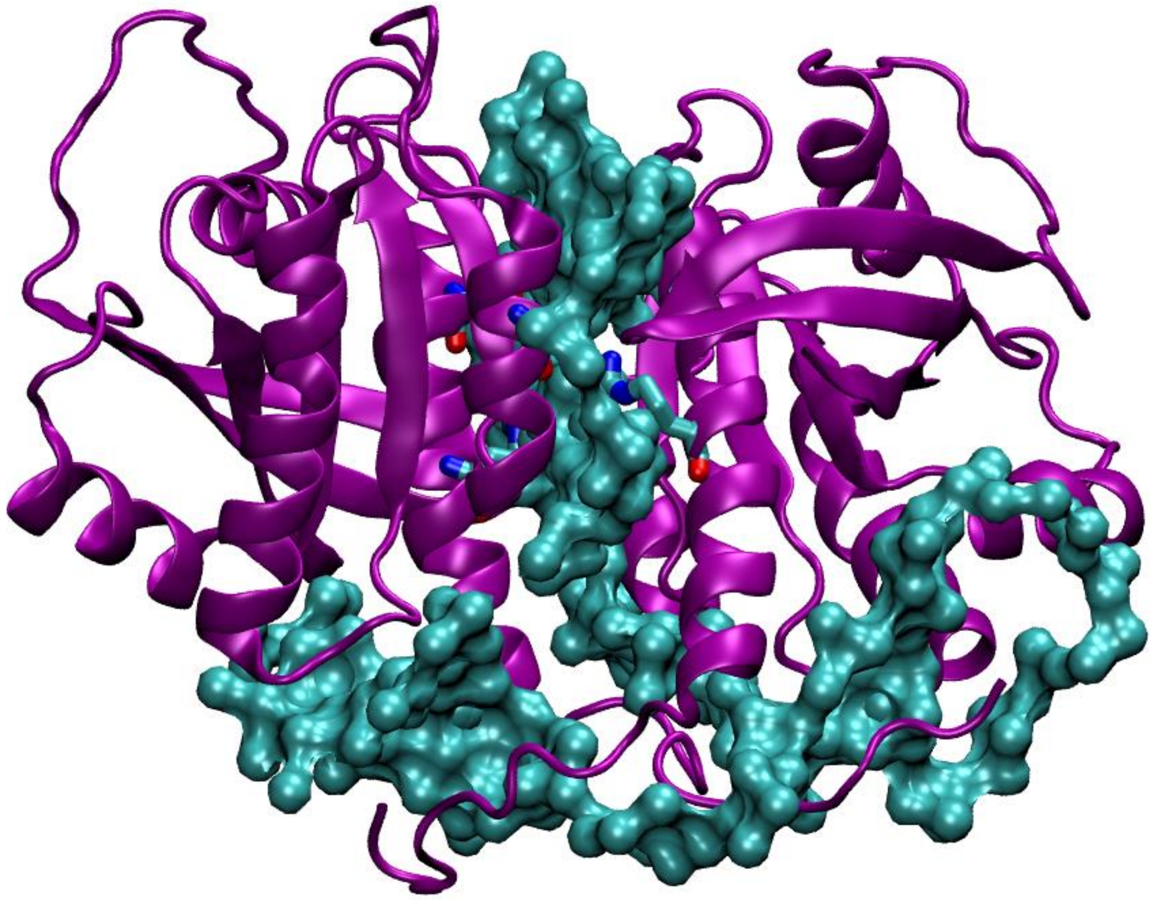

**Figure S33.** The water interface observed in the simulations based on the PDB ID 5VQ2 (Figure 5a) is replaced by a water interface in the simulations based on the PDB ID 6W4E.

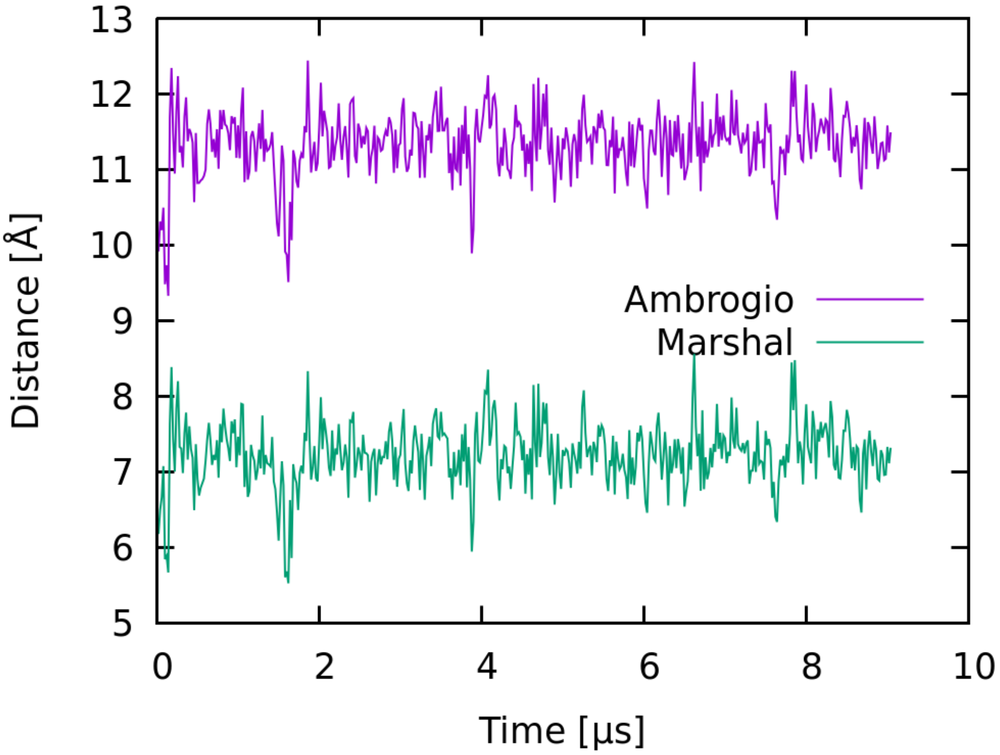


**Figure S34.** Unbiased MARTINI coarse-grained simulations. RMSD time series comparing the dimer structure from our coarse-grained simulations with PDB ID 5VQ2 (purple) and PDB ID 6W4E (green).

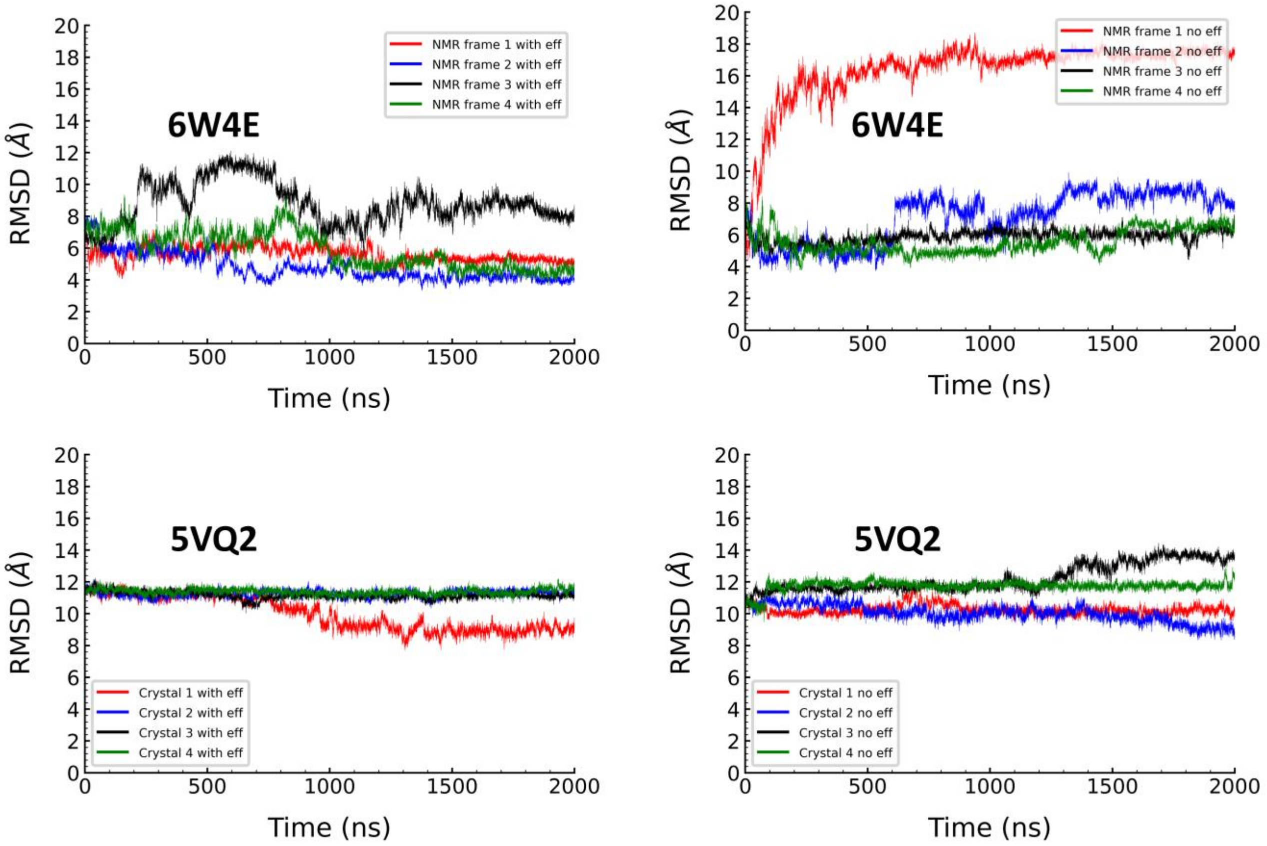


**Figure S35.** Full-time series of RMSD comparison of the dimer atomistic simulations, taking as reference the final frame of a 10 μs unbiased CG simulation.

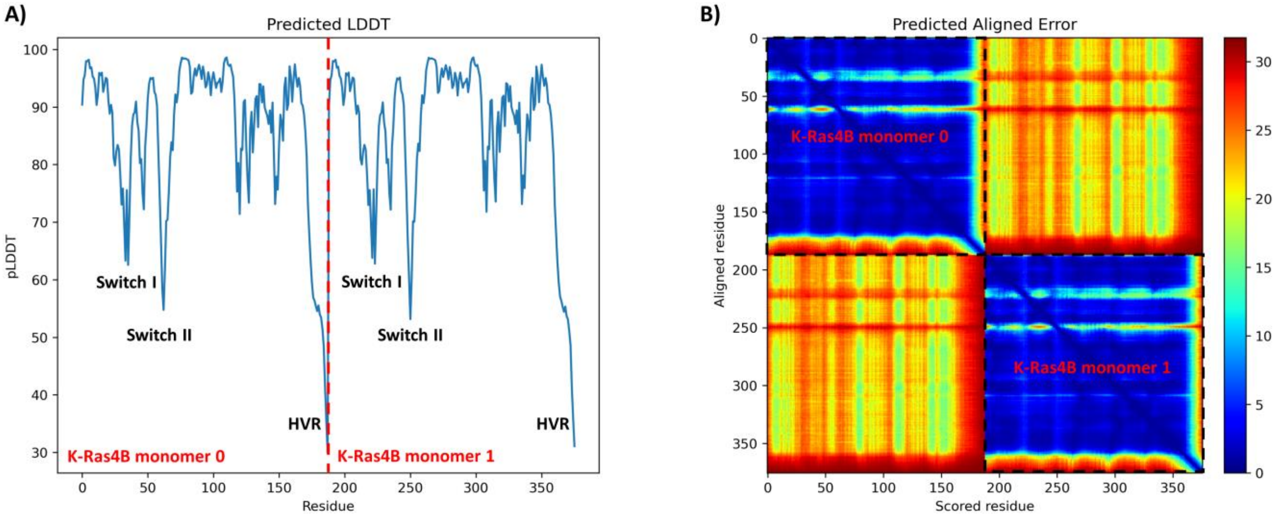


**Figure S36.** A) The pLDDT score shows that the predictions are accurate (pLDDT > 70) except for the Switch I, Switch II, and HVR regions. B) The Predicted Aligned Error (PAE) plot shows that the relative positions between the two monomers are uncertain, meaning that the protein-protein interface prediction is inaccurate.

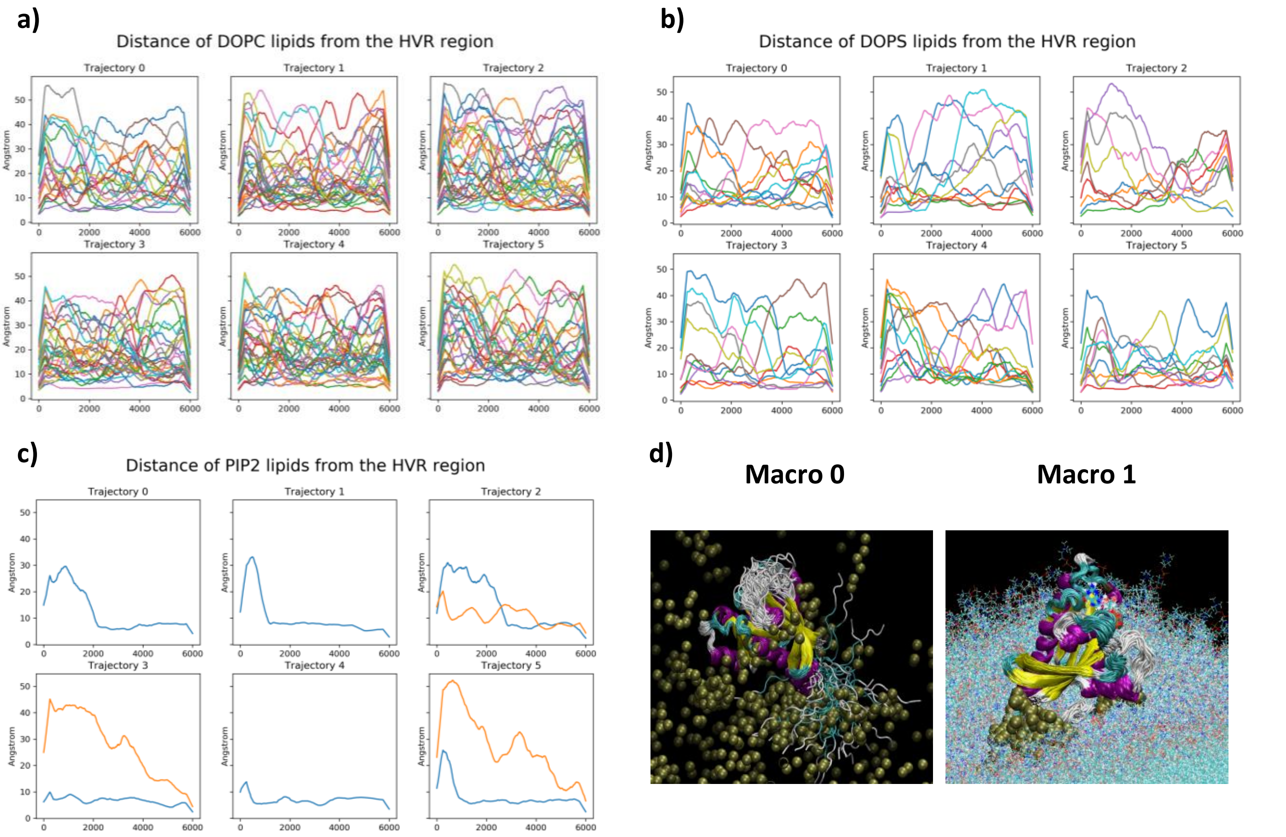


**Figure S37.** The minimum distance of each head group of a) DOPC, b) DOPS, and c) PIP2 lipids from the HVR tail in the six WT-GTP monomer K-Ras4B simulations. In the plots are presented only lipids whose heads stayed closer than 12 Å to the protein Cα atoms for more than 5% of the total simulation time of all six trajectories. (d) The two macrostates produced by the Markov state models.

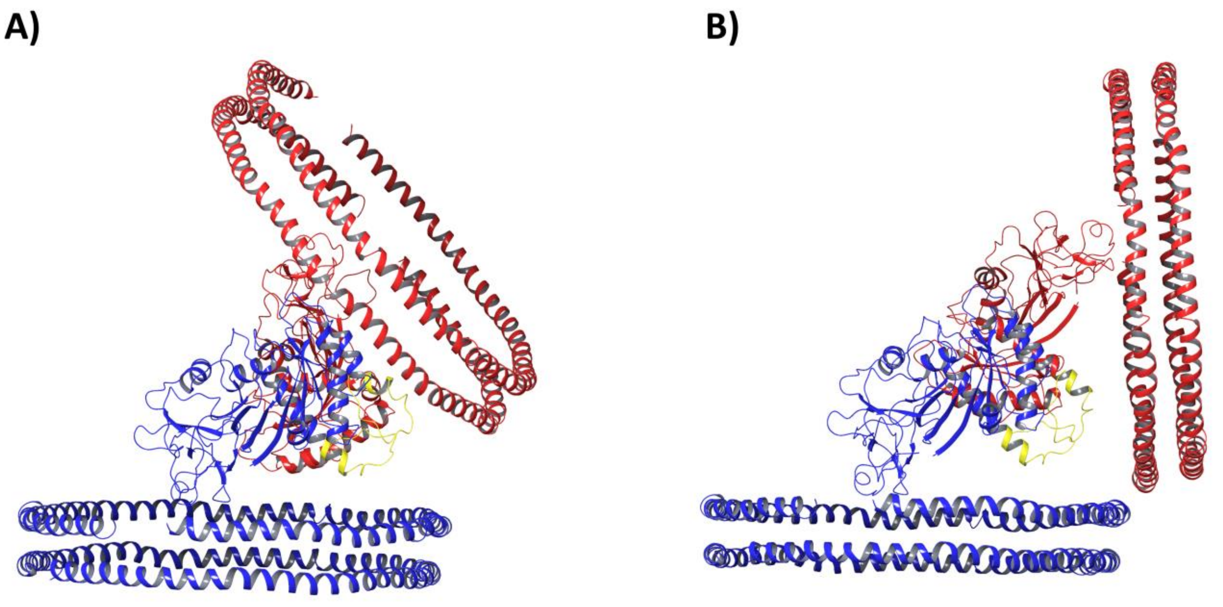


**Figure 38.** The 6PTW monomer structure aligned to A) 5VQ2 and B) 6W4E. The 6PTW structure was duplicated and each 6PTW structure was aligned to a different chain of the dimer. The different 6PTW structures are colored in red and blue and the HVRs in yellow.

**Supporting Tables**

**Table S1.** List of simulations.

| **System: Monomer K-Ras4B in solution** | | |
| --- | --- | --- |
| | **Simulation #** | **Simulation length (μs)** |
| **In solution** | GTP-bound Monomer #1 | 0.3 |
| | GTP-bound Monomer #2 | 0.3 |
| | GTP-bound Monomer #3 | 0.3 |
| | GTP-bound Monomer #4 | 0.3 |
| | GTP-bound Monomer #5 | 0.3 |
| | GTP-bound Monomer #6 | 0.3 |
| | GTP-bound Monomer #7 | 0.3 |
| | GTP-bound Monomer #8 | 0.3 |
| | GDP-bound Monomer #1 | 0.3 |
| | GDP-bound Monomer #2 | 0.3 |
| | GDP-bound Monomer #3 | 0.3 |
| | GDP-bound Monomer #4 | 0.3 |
| | GDP-bound Monomer #5 | 0.3 |
| | GDP-bound Monomer #6 | 0.3 |
| | GDP-bound Monomer #7 | 0.3 |
| | GDP-bound Monomer #8 | 0.3 |

| **System: Monomer K-Ras4B bound to the membrane** | | |
| --- | --- | --- |
| **Membrane composition** | **Simulation #** | **Simulation length (μs)** |
| **78% DOPC, 20% DOPS, 2% PIP2** | Monomer #1 | 0.6 |
| | Monomer #2 | 0.6 |
| | Monomer #3 | 0.6 |
| | Monomer #4 | 0.6 |
| | Monomer #5 | 0.6 |

| | | |
|---|---|---|
| | Monomer #6 | 0.6 |

| **System: Dimer K-Ras4B (5VQ2) bound to the membrane without Raf effectors** | | |
|---|---|---|
| **Membrane composition** | **Replica #** | **Simulation length (μs)** |
| **78% DOPC, 20% DOPS, 2% PIP2** | Crystal 1 no effectors | 2 |
| | Crystal 2 no effectors | 2 |
| | Crystal 3 no effectors | 2 |
| | Crystal 4 no effectors | 2 |

| **System: Dimer K-Ras4B (6W4E) bound to the membrane without Raf effectors** | | |
|---|---|---|
| **Membrane composition** | **Frame #** | **Simulation length (μs)** |
| **78% DOPC, 20% DOPS, 2% PIP2** | NMR frame 1 no effectors | 2 |
| | NMR frame 2 no effectors | 2 |
| | NMR frame 3 no effectors | 2 |
| | NMR frame 4 no effectors | 2 |

| **System: Dimer K-Ras4B (5VQ2) bound to the membrane with Raf effectors** | | |
|---|---|---|
| **Membrane composition** | **Replica #** | **Simulation length (μs)** |
| **78% DOPC, 20% DOPS, 2% PIP2** | Crystal 1 with effectors | 2 |
| | Crystal 2 with effectors | 2 |
| | Crystal 3 with effectors | 2 |
| | Crystal 4 with effectors | 2 |

| **System: Dimer K-Ras4B (6W4E) bound to the membrane with Raf effectors** | | |
|---|---|---|
| **Membrane composition** | **Frame #** | **Simulation length (μs)** |
| **78% DOPC, 20% DOPS, 2% PIP2** | NMR frame 1 with effectors | 2 |
| | NMR frame 2 with effectors | 2 |

| | | |
|---|---|---|
| | NMR frame 3 with effectors | 2 |
| | NMR frame 4 with effectors | 2 |

**Table S2.** Average distances of key salt bridges for the structure based on PDB ID 5VQ2 and structure based on PDB ID 6W4E.

| Simulation on 78/20/2 membrane | Average salt-bridge distance (pair 1, Å) | Average salt-bridge distance (pair 2, Å) |
|---|---|---|
| **D154-R161 salt bridge for structure based on 5VQ2** | | |
| Crystal 1 no eff | 9.7 ± 1.6 | 6.9 ± 1.9 |
| Crystal 2 no eff | 10.3 ± 3.1 | 9.8 ± 2.7 |
| Crystal 3 no eff | 6.2 ± 2.4 | 6.1 ± 1.7 |
| Crystal 4 no eff | 11.4 ± 1.1 | 5 ± 0.8 |
| Crystal 1 with eff | 9.3 ± 4.4 | 8.7 ± 3.6 |
| Crystal 2 with eff | 4.6 ± 0.7 | 4.3 ± 0.9 |
| Crystal 3 with eff | 4.2 ± 0.5 | 4.6 ± 0.7 |
| Crystal 4 with eff | 4.4 ± 0.6 | 4.8 ± 0.8 |
| **R135-E168 salt bridge for structure based on 6W4E** | | |
| NMR Frame 1 no eff | 16.5 ± 2.4 | - |
| NMR Frame 2 no eff | 4.5 ± 2.2 | 18.4 ± 3 |
| NMR Frame 3 no eff | 17.4 ± 2.1 | 7.2 ± 3.3 |
| NMR Frame 4 no eff | 12.7 ± 2 | 18.7 ± 3.1 |
| NMR Frame 1 with eff | 8.3 ± 3.3 | 16.2 ± 1.7 |
| NMR Frame 2 with eff | 12.3 ± 4.6 | 15.5 ± 4 |
| NMR Frame 3 with eff | 11.6 ± 6.1 | 16.8 ± 12.5 |
| NMR Frame 4 with eff | 7.8 ± 5.2 | 19.1 ± 3.3 |

**Table S3.** The interface pairs of amino acids with correlated motions that were found in the NMR (6W4E) and Crystal (5VQ2) trajectories with and without effectors, the interface pair mean correlation over all four trajectories, and the interface pair correlation in each trajectory separately.

| NMR (6W4E) with effectors | | | | | |
|---|---|---|---|---|---|
| **Interface pair** | **Mean correlation** | **Frame 1** | **Frame 2** | **Frame 3** | **Frame 4** |
| **R135-D154** | **0.38** | **0.77** | **0** | **0** | **0.74** |
| **D105-K180** | **0.36** | **0.72** | **0** | **0** | **0.72** |
| **R161-R135** | **0.22** | **0** | **0.86** | **0** | **0** |
| **S136-R161** | **0.2** | **0.81** | **0** | **0** | **0** |
| **S181-D173** | **0.2** | **0** | **0.81** | **0** | **0** |
| **D132-R161** | **0.2** | **0.81** | **0** | **0** | **0** |
| **D105-K182** | **0.2** | **0** | **0.8** | **0** | **0** |
| **R135-T158** | **0.2** | **0.8** | **0** | **0** | **0** |
| **R135-D153** | **0.19** | **0** | **0** | **0** | **0.78** |
| **R135-R161** | **0.19** | **0.78** | **0** | **0** | **0** |
| **R135-Y157** | **0.19** | **0** | **0** | **0** | **0.77** |
| **D105-K178** | **0.19** | **0** | **0** | **0** | **0.76** |
| **D173-K169** | **0.18** | **0.73** | **0** | **0** | **0** |
| **K177-D108** | **0.17** | **0.7** | **0** | **0** | **0** |
| **NMR (6W4E) without effectors** | | | | | |
| **Interface pair** | **Mean correlation** | **Frame 1** | **Frame 2** | **Frame 3** | **Frame 4** |
| **K172-R135** | **0.22** | **0.88** | **0** | **0** | **0** |
| **E168-R135** | **0.22** | **0** | **0.87** | **0** | **0** |
| **G174-R135** | **0.21** | **0.85** | **0** | **0** | **0** |
| **K179-E107** | **0.21** | **0.84** | **0** | **0** | **0** |
| **K175-R135** | **0.21** | **0.83** | **0** | **0** | **0** |
| **K165-S136** | **0.20** | **0** | **0** | **0** | **0.79** |
| **E162-R135** | **0.20** | **0** | **0** | **0** | **0.79** |
| **R161-S136** | **0.20** | **0** | **0** | **0.78** | **0** |
| **R135-Y157** | **0.19** | **0** | **0.77** | **0** | **0** |
| **T158-R135** | **0.19** | **0** | **0** | **0.77** | **0** |
| **R135-D153** | **0.19** | **0** | **0.77** | **0** | **0** |
| **K180-D105** | **0.19** | **0** | **0** | **0** | **0.77** |
| **R161-R135** | **0.19** | **0** | **0** | **0.77** | **0** |
| **K104-K178** | **0.19** | **0** | **0.76** | **0** | **0** |
| **D105-K178** | **0.19** | **0** | **0.76** | **0** | **0** |
| **G138-R135** | **0.19** | **0** | **0** | **0** | **0.75** |
| **D105-K179** | **0.19** | **0** | **0.74** | **0** | **0** |
| **D154-R135** | **0.19** | **0** | **0** | **0.74** | **0** |
| **D105-K182** | **0.18** | **0** | **0.73** | **0** | **0** |
| **R102-K182** | **0.18** | **0** | **0.73** | **0** | **0** |

| | | | | | |
|---|---|---|---|---|---|
| **R135-P140** | **0.18** | **0** | **0** | **0** | **0.73** |
| **R102-T183** | **0.18** | **0** | **0.73** | **0** | **0** |
| **P140-R135** | **0.18** | **0** | **0** | **0** | **0.73** |
| **R135-D154** | **0.18** | **0** | **0.72** | **0** | **0.00** |
| **R135-I142** | **0.18** | **0** | **0** | **0** | **0.72** |
| **K169-D108** | **0.17** | **0** | **0** | **0.69** | **0** |
| **R135-T158** | **0.17** | **0** | **0** | **0** | **0.66** |
| **M170-K179** | **0.16** | **0** | **0** | **0.64** | **0** |
| **D173-S181** | **0.16** | **0** | **0** | **0.64** | **0** |
| **D173-K182** | **0.16** | **0** | **0** | **0.64** | **0** |
| **D173-K179** | **0.16** | **0** | **0** | **0.64** | **0** |
| **D132-D154** | **0.16** | **0** | **0** | **0.00** | **0.63** |
| **D173-K178** | **0.16** | **0** | **0** | **0.62** | **0** |
| **D173-K180** | **0.16** | **0** | **0** | **0.62** | **0** |
| **G174-K182** | **0.15** | **0** | **0** | **0.62** | **0** |
| **M170-K177** | **0.15** | **0** | **0** | **0.62** | **0** |
| **M170-K178** | **0.15** | **0** | **0** | **0.61** | **0** |
| **R135-K165** | **0.15** | **0** | **0** | **0.59** | **0** |
| **R135-R161** | **0.15** | **0** | **0** | **0.59** | **0** |
| **G138-K165** | **0.14** | **0** | **0** | **0.57** | **0** |
| **K128-D154** | **0.14** | **0** | **0** | **0** | **0.55** |
| **D108-K169** | **0.14** | **0** | **0** | **0.55** | **0** |
| **Crystal (5VQ2) with effectors** | | | | | |
| **Interface pair** | **Mean correlation** | **Replica 1** | **Replica 2** | **Replica 3** | **Replica 4** |
| **K165-G138** | **0.74** | **0.85** | **0.74** | **0.64** | **0.72** |
| **R164-R135** | **0.73** | **0.86** | **0.7** | **0.63** | **0.73** |
| **G138-K165** | **0.64** | **0.82** | **0.57** | **0.54** | **0.63** |
| **D47-E143** | **0.58** | **0** | **0.74** | **0.72** | **0.85** |
| **G48-E143** | **0.56** | **0** | **0.72** | **0.68** | **0.84** |
| **K165-P140** | **0.55** | **0.84** | **0** | **0.63** | **0.73** |
| **K165-I139** | **0.54** | **0.85** | **0** | **0.63** | **0.7** |
| **E168-R135** | **0.54** | **0** | **0.72** | **0.64** | **0.81** |
| **E143-G48** | **0.53** | **0** | **0.74** | **0.65** | **0.74** |
| **I142-D47** | **0.53** | **0** | **0.77** | **0.61** | **0.74** |
| **E143-D47** | **0.52** | **0** | **0.75** | **0.61** | **0.73** |
| **F141-D47** | **0.52** | **0** | **0.75** | **0.6** | **0.73** |
| **K165-R135** | **0.51** | **0** | **0.7** | **0.62** | **0.72** |
| **R161-D154** | **0.5** | **0** | **0.66** | **0.63** | **0.71** |
| **R135-K165** | **0.5** | **0.8** | **0.56** | **0** | **0.63** |
| **R161-R161** | **0.48** | **0** | **0.62** | **0.61** | **0.69** |
| **E162-E162** | **0.47** | **0** | **0.61** | **0.6** | **0.68** |
| **D154-R161** | **0.46** | **0** | **0.6** | **0.6** | **0.66** |
| **D47-F141** | **0.45** | **0** | **0.56** | **0.51** | **0.72** |
| **T158-R161** | **0.44** | **0** | **0.58** | **0.54** | **0.65** |

| | | | | | |
|---|---|---|---|---|---|
| R161-T158 | 0.44 | 0 | 0.59 | 0.54 | 0.64 |
| D47-I142 | 0.43 | 0 | 0.52 | 0.5 | 0.7 |
| K167-R135 | 0.36 | 0 | 0 | 0.64 | 0.8 |
| R135-R164 | 0.36 | 0.8 | 0 | 0 | 0.64 |
| E49-Q131 | 0.32 | 0 | 0.68 | 0.59 | 0 |
| K165-E162 | 0.31 | 0 | 0 | 0.58 | 0.66 |
| I142-R161 | 0.3 | 0 | 0 | 0.56 | 0.65 |
| R135-D173 | 0.29 | 0 | 0 | 0.47 | 0.7 |
| Y4-R135 | 0.28 | 0 | 0.62 | 0.51 | 0 |
| M170-K128 | 0.17 | 0 | 0 | 0 | 0.7 |
| D154-D47 | 0.17 | 0 | 0 | 0 | 0.69 |
| R135-E168 | 0.17 | 0 | 0 | 0 | 0.69 |
| E168-D132 | 0.17 | 0 | 0.69 | 0 | 0 |
| D47-D154 | 0.17 | 0 | 0.69 | 0 | 0 |
| M170-Q131 | 0.17 | 0 | 0 | 0 | 0.67 |
| R135-K167 | 0.16 | 0 | 0 | 0 | 0.65 |
| K169-R135 | 0.16 | 0 | 0 | 0.62 | 0 |
| I139-K165 | 0.14 | 0 | 0.56 | 0 | 0 |
| Q131-R164 | 0.14 | 0 | 0.56 | 0 | 0 |
| P140-K165 | 0.14 | 0 | 0.55 | 0 | 0 |
| D132-D173 | 0.13 | 0 | 0 | 0.52 | 0 |
| Crystal (5VQ2) without effectors | | | | | |
| Interface pair | Mean correlation | Replica 1 | Replica 2 | Replica 3 | Replica 4 |
| M170-R135 | 0.73 | 0.74 | 0.76 | 0.78 | 0.62 |
| D173-R135 | 0.7 | 0.7 | 0.74 | 0.75 | 0.59 |
| R135-E168 | 0.64 | 0.64 | 0.71 | 0.7 | 0.5 |
| K165-G138 | 0.6 | 0.78 | 0.81 | 0.82 | 0 |
| K165-I139 | 0.6 | 0.78 | 0.8 | 0.82 | 0 |
| K165-P140 | 0.59 | 0.77 | 0.8 | 0.81 | 0 |
| K167-R135 | 0.59 | 0.78 | 0.78 | 0.81 | 0 |
| E168-R135 | 0.57 | 0.79 | 0.79 | 0 | 0.69 |
| G138-K165 | 0.53 | 0.66 | 0.74 | 0.74 | 0 |
| R135-K165 | 0.5 | 0.62 | 0.69 | 0.7 | 0 |
| R135-R164 | 0.45 | 0.62 | 0.69 | 0 | 0.48 |
| R164-R135 | 0.39 | 0.77 | 0.79 | 0 | 0 |
| K165-R135 | 0.39 | 0.76 | 0.79 | 0 | 0 |
| S136-E168 | 0.36 | 0.68 | 0.76 | 0 | 0 |
| R164-Q131 | 0.34 | 0.67 | 0.67 | 0 | 0 |
| R135-E49 | 0.33 | 0.61 | 0.69 | 0 | 0 |
| Q131-D47 | 0.31 | 0.65 | 0 | 0 | 0.6 |
| M170-Q131 | 0.3 | 0.65 | 0 | 0 | 0.56 |
| R161-R161 | 0.2 | 0 | 0 | 0.79 | 0 |
| E162-E162 | 0.2 | 0 | 0 | 0.78 | 0 |
| K165-E162 | 0.19 | 0 | 0 | 0.78 | 0 |

| | | | | | |
|---|---|---|---|---|---|
| **K165-K165** | **0.19** | **0** | **0** | **0.76** | **0** |
| **S136-K169** | **0.19** | **0** | **0** | **0.74** | **0** |
| **I139-K165** | **0.18** | **0** | **0** | **0.74** | **0** |
| **D132-K169** | **0.18** | **0** | **0** | **0.73** | **0** |
| **P140-K165** | **0.18** | **0** | **0** | **0.73** | **0** |
| **D132-M170** | **0.18** | **0** | **0** | **0.72** | **0** |
| **D132-S171** | **0.18** | **0** | **0** | **0.71** | **0** |
| **F141-G48** | **0.17** | **0** | **0** | **0** | **0.7** |
| **R135-K169** | **0.17** | **0** | **0** | **0.7** | **0** |
| **R135-K167** | **0.17** | **0** | **0** | **0.69** | **0** |
| **R135-M170** | **0.17** | **0** | **0** | **0.69** | **0** |
| **F141-D47** | **0.17** | **0** | **0** | **0** | **0.68** |
| **S171-K128** | **0.17** | **0.68** | **0** | **0** | **0** |
| **K165-F141** | **0.17** | **0** | **0** | **0** | **0.67** |
| **D132-E168** | **0.17** | **0.67** | **0** | **0** | **0** |
| **P140-D47** | **0.17** | **0** | **0** | **0** | **0.67** |
| **K165-I142** | **0.17** | **0** | **0** | **0** | **0.66** |
| **E49-K128** | **0.16** | **0.66** | **0** | **0** | **0** |
| **R161-D153** | **0.16** | **0** | **0** | **0** | **0.65** |
| **R161-D154** | **0.16** | **0** | **0** | **0** | **0.65** |
| **K165-D154** | **0.16** | **0** | **0** | **0** | **0.65** |
| **Q131-E49** | **0.16** | **0** | **0** | **0** | **0.64** |
| **M170-K128** | **0.15** | **0** | **0** | **0** | **0.59** |
| **K165-T158** | **0.15** | **0** | **0** | **0** | **0.59** |
| **P140-R161** | **0.15** | **0** | **0** | **0** | **0.58** |
| **K167-Q131** | **0.14** | **0** | **0** | **0** | **0.57** |
| **M170-D132** | **0.14** | **0** | **0** | **0** | **0.56** |
| **S136-K177** | **0.13** | **0** | **0** | **0** | **0.54** |
| **Q131-R164** | **0.12** | **0** | **0** | **0** | **0.49** |
| **R97-K177** | **0.12** | **0** | **0** | **0** | **0.48** |
| **D47-G151** | **0.11** | **0** | **0** | **0** | **0.42** |
| **D47-Q150** | **0.1** | **0** | **0** | **0** | **0.4** |
| **G48-Q150** | **0.1** | **0** | **0** | **0** | **0.39** |

**Table S4.** Cross-RMSD of the cluster representatives of the most populated cluster of 5VQ2 and 6W4E systems

| Cross-RMSD (Å) | NMR frame 1 no effectors | NMR frame 2 no effectors | NMR frame 3 no effectors | NMR frame 4 no effectors | NMR frame 1 with effectors | NMR frame 2 with effectors | NMR frame 3 with effectors | NMR frame 4 with effectors | Crystal 1 no effectors | Crystal 2 no effectors | Crystal 3 no effectors | Crystal 4 no effectors | Crystal 1 with effectors | Crystal 2 with effectors | Crystal 3 with effectors | Crystal 4 with effectors |
|---|---|---|---|---|---|---|---|---|---|---|---|---|---|---|---|---|
| **NMR frame 1 no effectors** | | 13.37 | 15.53 | 17.6 | 16.37 | 17.18 | 16.14 | 16.09 | 12.04 | 12.76 | 11.27 | 10.32 | 13.33 | 11.58 | 11.66 | 11.49 |
| **NMR frame 2 no effectors** | 13.37 | | 6.82 | 7.75 | 6.34 | 7.32 | 11.86 | 6.27 | 7.48 | 8.12 | 8.11 | 7.3 | 7.67 | 7.76 | 7.43 | 7.61 |
| **NMR frame 3 no effectors** | 15.53 | 6.82 | | 4.76 | 4.25 | 3.73 | 8.42 | 4.99 | 6.85 | 6.57 | 8.27 | 8.8 | 5.83 | 7.98 | 7.98 | 8.15 |
| **NMR frame 4 no effectors** | 17.6 | 7.75 | 4.76 | | 3.1 | 2.56 | 10.19 | 3.55 | 8.94 | 8.41 | 10.53 | 10.72 | 7.78 | 10.17 | 10.2 | 10.43 |
| **NMR frame 1 with effectors** | 16.37 | 6.34 | 4.25 | 3.1 | | 2.75 | 10.58 | 2.61 | 7.39 | 6.95 | 8.94 | 9.05 | 6.27 | 8.47 | 8.41 | 8.64 |
| **NMR frame 2 with effectors** | 17.18 | 7.32 | 3.73 | 2.56 | 2.75 | | 9.15 | 3.59 | 8.51 | 7.97 | 10.02 | 10.35 | 7.22 | 9.61 | 9.57 | 9.79 |

| | | | | | | | | | | | | | | | | |
|---|---|---|---|---|---|---|---|---|---|---|---|---|---|---|---|---|
| **NMR frame 3 with effectors** | 16.14 | 11.86 | 8.42 | 10.19 | 10.58 | 9.15 | | 10.65 | 11.24 | 10.81 | 12.56 | 12.94 | 10.35 | 12.49 | 12.69 | 12.59 |
| **NMR frame 4 with effectors** | 16.09 | 6.27 | 4.99 | 3.55 | 2.61 | 3.59 | 10.65 | | 7.82 | 7.46 | 9.43 | 9.25 | 6.77 | 8.91 | 8.84 | 9.09 |
| **Crystal 1 no effectors** | 12.04 | 7.48 | 6.85 | 8.94 | 7.39 | 8.51 | 11.24 | 7.82 | | 1.88 | 2.68 | 3.24 | 2.65 | 2.57 | 3.02 | 2.87 |
| **Crystal 2 no effectors** | 12.76 | 8.12 | 6.57 | 8.41 | 6.95 | 7.97 | 10.81 | 7.46 | 1.88 | | 3.55 | 4.3 | 2.02 | 3.36 | 3.89 | 3.79 |
| **Crystal 3 no effectors** | 11.27 | 8.11 | 8.27 | 10.53 | 8.94 | 10.02 | 12.56 | 9.43 | 2.68 | 3.55 | | 2.67 | 4.5 | 1.56 | 2.29 | 2.01 |
| **Crystal 4 no effectors** | 10.32 | 7.3 | 8.8 | 10.72 | 9.05 | 10.35 | 12.94 | 9.25 | 3.24 | 4.3 | 2.67 | | 5.06 | 2.79 | 3.18 | 2.86 |
| **Crystal 1 with effectors** | 13.33 | 7.67 | 5.83 | 7.78 | 6.27 | 7.22 | 10.35 | 6.77 | 2.65 | 2.02 | 4.5 | 5.06 | | 4.04 | 4.29 | 4.22 |
| **Crystal 2 with effectors** | 11.58 | 7.76 | 7.98 | 10.17 | 8.47 | 9.61 | 12.49 | 8.91 | 2.57 | 3.36 | 1.56 | 2.79 | 4.04 | | 1.98 | 1.64 |
| **Crystal 3 with effectors** | 11.66 | 7.43 | 7.98 | 10.2 | 8.41 | 9.57 | 12.69 | 8.84 | 3.02 | 3.89 | 2.29 | 3.18 | 4.29 | 1.98 | | 1.62 |
| **Crystal 4 with effectors** | 11.49 | 7.61 | 8.15 | 10.43 | 8.64 | 9.79 | 12.59 | 9.09 | 2.87 | 3.79 | 2.01 | 2.86 | 4.22 | 1.64 | 1.62 | |

**Table S5.** Average of the 48 simulated distances versus the experimental values

| Distance # | PRE distances | NMR frame 1 no eff | NMR frame 2 no eff | NMR frame 3 no eff | NMR frame 4 no eff | NMR frame 1 with eff | NMR frame 2 with eff | NMR frame 3 with eff | NMR frame 4 with eff | Crystal 1 no eff | Crystal 2 no eff | Crystal 3 no eff | Crystal 4 no eff | Crystal 1 with eff | Crystal 2 with eff | Crystal 3 with eff | Crystal 4 with eff |
|---|---|---|---|---|---|---|---|---|---|---|---|---|---|---|---|---|---|
| 1 | 15.4 ±3.0 | 27.0 ± 2.8 | 19.2 ± 4.0 | 26.4 ± 1.3 | 30.6 ± 2.2 | 29.1 ± 2.9 | 28.0 ± 3.1 | 29.9 ± 4.2 | 30.2 ± 4.0 | 19.2 ± 2.5 | 22.2 ± 2.6 | 20.5 ± 3.2 | 18.9 ± 1.1 | 21.3 ± 3.2 | 18.0 ± 1.0 | 18.0 ± 1.0 | 17.8 ± 0.8 |
| 2 | 15.4 ±3.0 | 27.2 ± 3.2 | 19.1 ± 3.9 | 26.0 ± 1.5 | 30.2 ± 2.3 | 28.2 ± 3.1 | 27.3 ± 3.5 | 29.9 ± 4.2 | 29.6 ± 4.3 | 18.8 ± 2.5 | 21.7 ± 2.5 | 20.2 ± 3.6 | 18.6 ± 1.1 | 20.8 ± 3.0 | 17.8 ± 1.2 | 17.5 ± 1.2 | 17.5 ± 1.0 |
| 3 | 15.4 ±3.0 | 33.1 ± 2.6 | 23.8 ± 2.4 | 27.7 ± 2.0 | 30.6 ± 2.2 | 26.6 ± 1.5 | 28.3 ± 2.6 | 38.1 ± 7.7 | 28.2 ± 1.8 | 21.7 ± 1.7 | 23.4 ± 2.2 | 21.4 ± 3.4 | 19.3 ± 1.4 | 21.5 ± 3.3 | 17.7 ± 0.6 | 17.5 ± 0.8 | 17.6 ± 0.7 |
| 4 | 15.4 ±3.0 | 32.8 ± 2.7 | 22.6 ± 2.7 | 27.2 ± 2.1 | 30.2 ± 2.3 | 25.9 ± 2.0 | 27.2 ± 2.7 | 37.8 ± 7.8 | 27.0 ± 2.1 | 21.2 ± 2.0 | 22.7 ± 2.2 | 20.8 ± 3.3 | 18.6 ± 1.7 | 22.4 ± 3.4 | 17.0 ± 0.8 | 17.1 ± 1.1 | 17.1 ± 0.9 |
| 5 | 14.4 ±3.0 | 23.7 ± 3.4 | 18.1 ± 4.4 | 26.6 ± 1.6 | 31.9 ± 2.4 | 28.6 ± 3.1 | 29.2 ± 4.1 | 26.1 ± 4.2 | 29.1 ± 4.0 | 13.2 ± 2.4 | 16.3 ± 2.9 | 14.4 ± 4.0 | 13.5 ± 1.2 | 15.1 ± 3.7 | 11.3 ± 0.9 | 11.3 ± 1.1 | 10.9 ± 0.7 |
| 6 | 14.4 ±3.0 | 23.9 ± 3.5 | 19.6 ± 4.3 | 27.6 ± 1.6 | 33.2 ± 2.4 | 30.3 ± 3.1 | 30.2 ± 4.1 | 27.9 ± 4.1 | 30.9 ± 4.1 | 15.4 ± 2.4 | 18.5 ± 3.0 | 16.2 ± 4.0 | 15.5 ± 1.2 | 17.2 ± 3.8 | 13.3 ± 1.0 | 13.2 ± 1.2 | 12.9 ± 0.8 |
| 7 | 14.4 ±3.0 | 27.4 ± 3.1 | 24.2 ± 2.3 | 26.5 ± 1.8 | 31.8 ± 2.4 | 26.3 ± 1.7 | 28.7 ± 3.1 | 37.6 ± 10.3 | 27.1 ± 2.1 | 15.7 ± 1.5 | 17.6 ± 2.3 | 15.0 ± 3.6 | 12.7 ± 1.5 | 15.9 ± 3.9 | 10.8 ± 0.6 | 10.8 ± 0.8 | 10.9 ± 0.7 |
| 8 | 14.4 ±3.0 | 28.9 ± 3.2 | 24.0 ± 2.8 | 28.2 ± 1.9 | 32.8 ± 2.5 | 27.4 ± 1.7 | 29.9 ± 3.2 | 39.4 ± 9.9 | 27.9 ± 2.1 | 17.7 ± 1.7 | 19.6 ± 2.3 | 17.1 ± 3.6 | 14.1 ± 1.7 | 18.0 ± 4.0 | 12.7 ± 0.7 | 12.7 ± 0.9 | 12.8 ± 0.7 |
| 9 | 15.7 ±3.0 | 26.3 ± 2.6 | 17.0 ± 3.8 | 24.4 ± 1.3 | 27.3 ± 2.2 | 25.7 ± 2.9 | 25.2 ± 3.2 | 27.0 ± 3.6 | 26.3 ± 4.4 | 16.7 ± 2.0 | 19.4 ± 2.1 | 18.5 ± 2.8 | 15.8 ± 0.9 | 18.8 ± 2.4 | 16.4 ± 0.7 | 16.5 ± 0.7 | 16.2 ± 0.5 |

| | | | | | | | | | | | | | | | | |
|---|---|---|---|---|---|---|---|---|---|---|---|---|---|---|---|---|
| **10** | **15.7 ±3.0** | **31.1 ± 2.4** | **24.2 ± 1.5** | **24.8 ± 1.9** | **27.7 ± 1.9** | **24.7 ± 1.6** | **25.6 ± 2.5** | **34.5 ± 8.2** | **26.5 ± 1.9** | **19.6 ± 1.3** | **20.6 ± 1.9** | **18.8 ± 2.7** | **18.0 ± 1.2** | **19.5 ± 2.8** | **15.9 ± 0.7** | **15.9 ± 0.5** | **16.0 ± 0.4** |
| **11** | **13.3 ±3.0** | **36.0 ± 3.6** | **13.9 ± 2.2** | **18.7 ± 1.5** | **18.7 ± 1.6** | **18.8 ± 2.2** | **18.1 ± 1.7** | **27.0 ± 4.4** | **20.2 ± 3.1** | **22.3 ± 1.7** | **24.0 ± 1.4** | **25.9 ± 2.6** | **23.2 ± 0.9** | **23.4 ± 1.1** | **23.6 ± 0.8** | **23.3 ± 0.6** | **23.2 ± 0.6** |
| **12** | **13.3 ±3.0** | **37.7 ± 3.3** | **19.7 ± 1.7** | **18.3 ± 2.0** | **19.9 ± 1.5** | **18.8 ± 1.5** | **18.6 ± 1.6** | **30.1 ± 4.3** | **22.5 ± 2.2** | **23.5 ± 1.3** | **24.6 ± 1.4** | **25.8 ± 2.6** | **25.6 ± 0.8** | **23.3 ± 1.0** | **23.4 ± 0.7** | **22.9 ± 0.7** | **23.0 ± 0.6** |
| **13** | **16 ±3.0** | **24.8 ± 2.6** | **18.3 ± 3.5** | **26.3 ± 1.3** | **28.3 ± 1.6** | **27.2 ± 2.5** | **26.5 ± 2.1** | **27.9 ± 2.6** | **27.2 ± 3.5** | **17.0 ± 1.4** | **19.6 ± 2.0** | **18.1 ± 1.9** | **15.4 ± 0.7** | **19.1 ± 2.3** | **17.2 ± 0.7** | **17.0 ± 0.7** | **16.6 ± 0.5** |
| **14** | **16 ±3.0** | **26.3 ± 2.7** | **19.0 ± 3.5** | **26.6 ± 1.2** | **28.5 ± 1.6** | **27.5 ± 2.5** | **26.8 ± 2.1** | **29.3 ± 2.9** | **27.9 ± 3.4** | **19.1 ± 1.5** | **21.7 ± 2.0** | **20.4 ± 2.0** | **17.7 ± 0.7** | **21.1 ± 2.2** | **19.2 ± 0.7** | **19.1 ± 0.7** | **18.7 ± 0.5** |
| **15** | **16 ±3.0** | **34.2 ± 2.3** | **26.0 ± 1.3** | **25.6 ± 2.0** | **29.7 ± 1.7** | **25.8 ± 1.3** | **27.4 ± 2.1** | **34.7 ± 7.3** | **29.0 ± 1.6** | **20.8 ± 1.2** | **21.4 ± 1.4** | **18.6 ± 2.1** | **20.2 ± 0.8** | **20.1 ± 2.9** | **16.6 ± 0.6** | **16.3 ± 0.6** | **16.5 ± 0.5** |
| **16** | **16 ±3.0** | **35.8 ± 2.3** | **25.7 ± 1.5** | **26.2 ± 2.0** | **29.8 ± 1.6** | **26.3 ± 1.3** | **27.7 ± 2.1** | **35.5 ± 6.6** | **29.3 ± 1.5** | **22.5 ± 1.2** | **23.3 ± 1.5** | **20.9 ± 2.2** | **21.8 ± 0.8** | **22.0 ± 2.7** | **18.8 ± 0.6** | **18.3 ± 0.5** | **18.5 ± 0.5** |
| **17** | **15.2 ±3.0** | **46.7 ± 4.6** | **20.7 ± 1.7** | **20.1 ± 1.5** | **15.4 ± 1.7** | **18.9 ± 1.8** | **18.0 ± 2.5** | **32.7 ± 4.5** | **21.3 ± 3.0** | **31.8 ± 1.5** | **33.0 ± 1.6** | **36.4 ± 2.3** | **33.0 ± 1.0** | **32.3 ± 1.9** | **34.2 ± 1.1** | **33.6 ± 1.2** | **34.1 ± 0.8** |
| **18** | **15.2 ±3.0** | **45.2 ± 5.0** | **23.8 ± 2.4** | **20.0 ± 1.9** | **17.3 ± 2.1** | **21.2 ± 1.9** | **16.9 ± 2.1** | **29.2 ± 6.6** | **25.3 ± 3.3** | **32.5 ± 1.3** | **33.0 ± 1.6** | **36.2 ± 2.5** | **35.1 ± 1.1** | **32.6 ± 1.7** | **34.3 ± 0.8** | **34.1 ± 0.7** | **33.9 ± 0.7** |
| **19** | **13.8 ±3.0** | **37.6 ± 3.8** | **15.2 ± 2.3** | **19.4 ± 1.2** | **19.6 ± 1.6** | **20.0 ± 2.0** | **18.7 ± 1.5** | **28.8 ± 4.6** | **21.7 ± 3.1** | **24.1 ± 1.9** | **26.0 ± 1.4** | **27.8 ± 2.7** | **25.2 ± 0.7** | **25.0 ± 1.1** | **25.3 ± 0.8** | **24.8 ± 0.7** | **24.8 ± 0.6** |
| **20** | **13.8 ±3.0** | **39.4 ± 3.5** | **19.6 ± 1.5** | **19.6 ± 1.9** | **21.0 ± 1.6** | **19.8 ± 1.5** | **19.4 ± 1.5** | **31.4 ± 4.2** | **23.1 ± 2.2** | **25.3 ± 1.4** | **26.3 ± 1.5** | **27.9 ± 2.8** | **26.5 ± 0.7** | **25.1 ± 1.2** | **25.1 ± 0.7** | **24.6 ± 0.7** | **24.7 ± 0.6** |

| | | | | | | | | | | | | | | | | | |
|---|---|---|---|---|---|---|---|---|---|---|---|---|---|---|---|---|---|
| 21 | 13.7 ±3.0 | 39.7 ± 4.0 | 18.5 ± 2.4 | 23.3 ± 1.4 | 23.4 ± 1.8 | 23.0 ± 2.2 | 21.8 ± 1.9 | 32.1 ± 5.3 | 25.5 ± 3.2 | 27.7 ± 2.4 | 29.7 ± 1.8 | 31.0 ± 3.2 | 29.0 ± 1.1 | 28.3 ± 1.5 | 28.0 ± 1.1 | 27.7 ± 1.0 | 27.8 ± 1.1 |
| 22 | 13.7 ±3.0 | 42.3 ± 3.8 | 19.4 ± 2.3 | 23.1 ± 2.0 | 23.3 ± 1.8 | 22.5 ± 1.6 | 22.1 ± 1.6 | 35.0 ± 4.5 | 24.9 ± 2.4 | 28.3 ± 1.9 | 30.0 ± 1.9 | 31.2 ± 3.1 | 29.1 ± 1.2 | 28.2 ± 1.7 | 28.2 ± 1.1 | 27.4 ± 1.0 | 27.6 ± 1.0 |
| 23 | 14.4 ±3.0 | 47.2 ± 5.3 | 19.6 ± 3.7 | 20.1 ± 2.5 | 26.1 ± 2.5 | 26.3 ± 2.4 | 24.3 ± 2.6 | 33.0 ± 6.9 | 29.9 ± 3.4 | 32.9 ± 2.9 | 35.0 ± 2.0 | 37.2 ± 4.1 | 36.2 ± 1.1 | 33.3 ± 1.7 | 32.8 ± 1.2 | 32.0 ± 1.0 | 32.6 ± 0.9 |
| 24 | 14.4 ±3.0 | 44.4 ± 5.0 | 14.4 ± 4.8 | 26.4 ± 2.0 | 25.6 ± 2.4 | 23.7 ± 2.1 | 24.5 ± 2.4 | 40.4 ± 5.6 | 21.8 ± 2.7 | 32.6 ± 2.2 | 34.7 ± 2.2 | 35.1 ± 2.0 | 32.8 ± 1.0 | 32.7 ± 1.8 | 33.1 ± 1.1 | 32.1 ± 0.9 | 32.5 ± 0.9 |
| 25 | 15.9 ±3.0 | 33.6 ± 3.6 | 26.2 ± 5.3 | 16.9 ± 1.2 | 21.6 ± 1.8 | 20.9 ± 2.5 | 20.6 ± 2.5 | 31.7 ± 10.4 | 25.5 ± 1.9 | 15.9 ± 1.5 | 16.4 ± 1.3 | 14.1 ± 1.0 | 20.1 ± 1.1 | 21.6 ± 2.0 | 20.1 ± 3.1 | 21.1 ± 3.1 | 17.1 ± 1.2 |
| 26 | 15.9 ±3.0 | 32.5 ± 3.6 | 25.1 ± 5.0 | 16.0 ± 1.2 | 21.1 ± 1.6 | 20.1 ± 2.4 | 19.7 ± 2.2 | 30.2 ± 10.4 | 24.9 ± 1.9 | 14.6 ± 1.4 | 15.1 ± 1.2 | 12.1 ± 1.0 | 18.6 ± 1.1 | 19.9 ± 2.0 | 18.0 ± 3.1 | 19.1 ± 2.9 | 14.8 ± 1.2 |
| 27 | 15.9 ±3.0 | 23.8 ± 2.4 | 20.1 ± 2.4 | 20.2 ± 1.3 | 18.2 ± 1.9 | 20.7 ± 1.4 | 19.5 ± 1.8 | 18.1 ± 2.7 | 21.7 ± 2.9 | 21.8 ± 0.7 | 21.7 ± 1.3 | 22.8 ± 2.7 | 17.4 ± 2.5 | 21.3 ± 2.0 | 22.0 ± 1.2 | 15.3 ± 1.6 | 17.2 ± 0.5 |
| 28 | 15.9 ±3.0 | 22.4 ± 2.3 | 18.5 ± 2.3 | 19.5 ± 1.4 | 17.2 ± 1.8 | 19.3 ± 1.4 | 18.5 ± 1.7 | 17.4 ± 3.0 | 20.3 ± 3.0 | 19.4 ± 0.7 | 19.9 ± 1.0 | 20.5 ± 2.7 | 15.3 ± 2.4 | 19.9 ± 1.9 | 20.0 ± 1.1 | 12.9 ± 1.6 | 15.0 ± 0.5 |
| 29 | 16 ±3.0 | 39.5 ± 3.9 | 31.6 ± 4.4 | 23.1 ± 1.3 | 27.8 ± 1.6 | 26.8 ± 2.2 | 27.0 ± 2.0 | 35.1 ± 11.7 | 32.2 ± 2.3 | 20.2 ± 1.5 | 21.4 ± 1.3 | 15.5 ± 0.9 | 23.6 ± 1.0 | 23.4 ± 2.3 | 20.8 ± 3.4 | 22.3 ± 2.1 | 16.1 ± 1.4 |
| 30 | 16 ±3.0 | 41.7 ± 3.9 | 33.7 ± 4.5 | 25.1 ± 1.3 | 29.5 ± 1.7 | 28.7 ± 2.3 | 28.9 ± 2.1 | 37.3 ± 11.8 | 34.1 ± 2.3 | 22.3 ± 1.6 | 23.5 ± 1.3 | 17.8 ± 0.9 | 25.8 ± 1.0 | 25.6 ± 2.3 | 23.1 ± 3.5 | 24.5 ± 2.2 | 18.2 ± 1.4 |
| 31 | 16 ±3.0 | 27.6 ± 2.6 | 24.3 ± 2.3 | 26.6 ± 1.6 | 24.2 ± 1.9 | 26.3 ± 1.3 | 25.4 ± 1.6 | 24.0 ± 2.9 | 26.9 ± 3.2 | 20.4 ± 1.0 | 23.3 ± 1.2 | 23.2 ± 3.1 | 15.2 ± 3.1 | 26.1 ± 2.1 | 23.7 ± 1.3 | 14.5 ± 1.8 | 18.6 ± 0.6 |

| | | | | | | | | | | | | | | | | | |
|---|---|---|---|---|---|---|---|---|---|---|---|---|---|---|---|---|---|
| **32** | **16 ±3.0** | **29.8 ± 2.7** | **26.5 ± 2.3** | **28.4 ± 1.6** | **26.1 ± 1.9** | **28.5 ± 1.3** | **27.4 ± 1.7** | **25.7 ± 2.8** | **29.1 ± 3.1** | **22.7 ± 1.1** | **25.5 ± 1.2** | **25.6 ± 3.1** | **17.4 ± 3.2** | **28.2 ± 2.1** | **26.0 ± 1.2** | **16.9 ± 1.7** | **20.9 ± 0.6** |
| **33** | **13.5 ±3.0** | **34.2 ± 3.3** | **23.4 ± 4.9** | **15.2 ± 1.5** | **18.7 ± 2.1** | **17.9 ± 2.5** | **18.5 ± 2.4** | **28.4 ± 12.0** | **23.9 ± 2.7** | **12.0 ± 1.8** | **12.9 ± 1.6** | **9.3 ± 0.9** | **16.2 ± 1.3** | **16.5 ± 2.2** | **15.8 ± 3.9** | **16.8 ± 4.2** | **15.3 ± 1.4** |
| **34** | **13.5 ±3.0** | **34.5 ± 3.5** | **24.5 ± 5.0** | **16.0 ± 1.4** | **19.8 ± 2.0** | **19.1 ± 2.5** | **19.6 ± 2.4** | **29.4 ± 11.8** | **24.8 ± 2.5** | **13.3 ± 1.7** | **14.3 ± 1.4** | **10.7 ± 0.8** | **17.5 ± 1.1** | **18.1 ± 2.2** | **17.0 ± 3.7** | **18.1 ± 3.8** | **15.5 ± 1.3** |
| **35** | **13.5 ±3.0** | **25.0 ± 1.8** | **20.1 ± 3.1** | **17.5 ± 1.8** | **15.7 ± 2.4** | **20.0 ± 1.6** | **17.2 ± 2.0** | **15.2 ± 2.7** | **20.3 ± 3.7** | **18.2 ± 1.0** | **17.1 ± 1.9** | **20.3 ± 1.8** | **17.0 ± 1.6** | **18.0 ± 2.1** | **18.4 ± 1.4** | **13.1 ± 2.2** | **16.6 ± 0.5** |
| **36** | **13.5 ±3.0** | **25.0 ± 1.8** | **20.2 ± 2.9** | **18.8 ± 1.7** | **16.9 ± 2.2** | **20.6 ± 1.6** | **18.3 ± 1.9** | **16.2 ± 2.7** | **20.8 ± 3.5** | **19.1 ± 0.8** | **18.6 ± 1.5** | **20.8 ± 2.1** | **16.7 ± 2.0** | **19.3 ± 2.1** | **19.2 ± 1.5** | **13.3 ± 2.1** | **16.7 ± 0.6** |
| **37** | **15.2 ±3.0** | **29.5 ± 3.3** | **21.6 ± 5.6** | **12.2 ± 1.1** | **16.9 ± 1.8** | **16.3 ± 2.5** | **15.6 ± 2.7** | **27.8 ± 9.8** | **20.5 ± 1.8** | **11.9 ± 1.3** | **12.1 ± 1.2** | **12.2 ± 1.0** | **16.4 ± 1.1** | **18.6 ± 1.9** | **17.8 ± 2.8** | **18.7 ± 3.5** | **16.5 ± 1.1** |
| **38** | **15.2 ±3.0** | **21.1 ± 2.0** | **17.0 ± 3.1** | **15.4 ± 1.3** | **13.6 ± 1.9** | **16.3 ± 1.5** | **14.8 ± 1.9** | **13.7 ± 2.7** | **17.3 ± 2.9** | **20.6 ± 0.6** | **19.1 ± 1.5** | **20.8 ± 2.1** | **18.1 ± 1.6** | **16.9 ± 2.0** | **19.1 ± 1.5** | **14.6 ± 1.5** | **15.4 ± 0.5** |
| **39** | **15.9 ±3.0** | **35.5 ± 4.0** | **28.9 ± 3.6** | **21.1 ± 1.3** | **26.6 ± 1.3** | **24.9 ± 2.0** | **24.9 ± 1.6** | **31.3 ± 11.1** | **30.5 ± 2.5** | **17.9 ± 1.3** | **19.2 ± 1.2** | **12.0 ± 0.9** | **20.4 ± 0.9** | **19.6 ± 2.3** | **15.6 ± 3.0** | **17.3 ± 1.3** | **9.3 ± 1.4** |
| **40** | **15.9 ±3.0** | **22.9 ± 2.6** | **19.9 ± 2.7** | **25.2 ± 1.5** | **22.3 ± 1.7** | **22.7 ± 1.3** | **23.4 ± 1.6** | **23.1 ± 3.5** | **23.5 ± 3.1** | **14.1 ± 1.0** | **19.2 ± 1.9** | **16.9 ± 3.2** | **8.5 ± 3.0** | **23.1 ± 2.3** | **18.4 ± 1.6** | **8.4 ± 1.9** | **12.7 ± 0.6** |
| **41** | **17.1 ±3.0** | **16.9 ± 2.9** | **26.9 ± 4.0** | **19.5 ± 2.4** | **14.0 ± 2.2** | **12.4 ± 0.9** | **17.7 ± 2.1** | **29.4 ± 5.2** | **15.0 ± 5.8** | **29.1 ± 1.0** | **21.1 ± 2.5** | **31.2 ± 1.9** | **23.6 ± 2.0** | **33.8 ± 3.3** | **38.7 ± 4.0** | **42.9 ± 4.3** | **43.1 ± 2.8** |
| **42** | **17.1 ±3.0** | **14.7 ± 2.8** | **11.1 ± 2.8** | **11.4 ± 1.4** | **16.9 ± 2.3** | **18.3 ± 1.8** | **11.7 ± 1.5** | **15.0 ± 2.5** | **14.6 ± 2.3** | **36.9 ± 1.2** | **39.0 ± 3.3** | **31.5 ± 3.0** | **32.7 ± 1.8** | **33.9 ± 4.0** | **46.9 ± 4.8** | **40.8 ± 2.9** | **39.1 ± 3.0** |

| | | | | | | | | | | | | | | | | | |
|---|---|---|---|---|---|---|---|---|---|---|---|---|---|---|---|---|---|
| **43** | **17.5 ±3.0** | **14.8 ± 2.9** | **25.3 ± 3.9** | **19.6 ± 3.1** | **15.9 ± 2.4** | **12.8 ± 1.0** | **15.5 ± 1.8** | **30.9 ± 4.7** | **15.6 ± 5.7** | **26.1 ± 1.1** | **18.4 ± 2.4** | **28.8 ± 2.0** | **20.8 ± 2.0** | **36.1 ± 3.6** | **37.8 ± 3.3** | **43.4 ± 4.5** | **42.2 ± 3.4** |
| **44** | **17.5 ±3.0** | **12.4 ± 2.8** | **12.6 ± 2.9** | **10.3 ± 1.5** | **19.2 ± 2.7** | **16.9 ± 1.9** | **12.4 ± 1.6** | **14.7 ± 2.3** | **17.4 ± 2.2** | **33.7 ± 1.1** | **37.0 ± 2.9** | **28.2 ± 3.2** | **29.7 ± 1.7** | **36.3 ± 4.0** | **48.7 ± 5.3** | **41.6 ± 2.7** | **41.8 ± 3.2** |
| **45** | **17.2 ±3.0** | **11.7 ± 3.1** | **24.4 ± 3.8** | **21.8 ± 4.0** | **17.6 ± 2.4** | **14.0 ± 1.5** | **13.3 ± 1.8** | **30.2 ± 4.1** | **15.4 ± 5.9** | **23.9 ± 1.2** | **16.3 ± 2.3** | **25.6 ± 2.2** | **18.0 ± 2.1** | **38.4 ± 3.9** | **37.9 ± 3.0** | **43.3 ± 5.9** | **42.3 ± 4.1** |
| **46** | **17.2 ±3.0** | **10.3 ± 2.7** | **14.3 ± 2.4** | **10.1 ± 1.1** | **21.1 ± 3.2** | **17.1 ± 1.6** | **14.2 ± 1.8** | **13.5 ± 2.9** | **19.2 ± 2.3** | **32.5 ± 1.1** | **34.7 ± 3.0** | **28.3 ± 2.8** | **30.6 ± 1.5** | **38.3 ± 4.6** | **51.5 ± 6.0** | **41.0 ± 3.6** | **44.3 ± 3.3** |
| **47** | **17.2 ±3.0** | **10.7 ± 3.2** | **22.6 ± 3.9** | **24.1 ± 3.8** | **19.3 ± 1.9** | **13.4 ± 1.9** | **11.7 ± 1.4** | **30.0 ± 3.9** | **16.3 ± 5.0** | **21.3 ± 1.3** | **13.2 ± 2.1** | **23.9 ± 1.9** | **14.9 ± 2.2** | **40.9 ± 3.8** | **38.0 ± 2.7** | **43.9 ± 7.2** | **42.9 ± 4.6** |
| **48** | **17.2 ±3.0** | **8.8 ± 2.4** | **16.4 ± 2.4** | **10.7 ± 0.8** | **22.4 ± 3.1** | **16.1 ± 1.1** | **17.1 ± 2.1** | **13.0 ± 2.3** | **22.2 ± 2.5** | **30.2 ± 1.1** | **31.7 ± 3.2** | **27.9 ± 2.8** | **30.7 ± 1.6** | **40.0 ± 5.3** | **53.6 ± 6.4** | **42.9 ± 4.1** | **47.2 ± 3.3** |
